# Systematic engineering of synthetic serine cycles in Pseudomonas putida uncovers emergent topologies for methanol assimilation

**DOI:** 10.1101/2025.02.17.638773

**Authors:** Òscar Puiggené, Jaime Muñoz-Triviño, Laura Civil-Ferrer, Line Gille, Helena Schulz-Mirbach, Daniel Bergen, Tobias J. Erb, Birgitta E. Ebert, Pablo I. Nikel

**Affiliations:** The Novo Nordisk Foundation Center for Biosustainability, Technical University of Denmark, Kongens Lyngby, Denmark; Department of Biochemistry and Synthetic Metabolism, Max Planck Institute for Terrestrial Microbiology, Marburg, Germany; Australian Institute for Bioengineering and Nanotechnology (AIBN), The University of Queensland, Brisbane, Queensland, Australia; Advanced Engineering Biology Future Science Platform, CSIRO, Brisbane, Queensland, Australia; Center for Synthetic Microbiology (SYNMIKRO), Marburg, Germany

**Keywords:** *Pseudomonas putid*, *a* Growth-coupled selection, Metabolic engineering, Methylotrophy, Methanol, Serine cycle

## Abstract

The urgent need for a circular carbon economy has driven research into sustainable substrates, including one-carbon (C_1_) compounds. The non-pathogenic soil bacterium *Pseudomonas putida* is a promising host for exploring synthetic methylotrophy due to its versatile metabolism. In this work, we implemented synthetic serine cycle variants in *P. putida* for methanol assimilation combining modular engineering and growth-coupled selection, whereby methanol assimilation supported biosynthesis of the essential amino acid serine. The serine cycle forms acetyl-coenzyme A from C_1_ molecules without carbon loss but has bottlenecks that hinder engineering efforts. We adopted three synthetic variants (serine-threonine cycle, homoserine cycle, and modified serine cycle) that yield serine in a methanol-dependent fashion to overcome these challenges. By dividing these metabolic designs into functional modules, we systematically compared their performance for implementation *in vivo*. Additionally, we harnessed native pyrroloquinoline quinone-dependent dehydrogenases for engineering methylotrophy. Recursive rewiring of synthetic and native activities revealed novel metabolic topologies for methanol utilization, termed enhanced serine-threonine cycle, providing a blueprint for engineering C_1_ assimilation in non-model heterotrophic bacteria.

**GRAPHICAL ABSTRACT:** 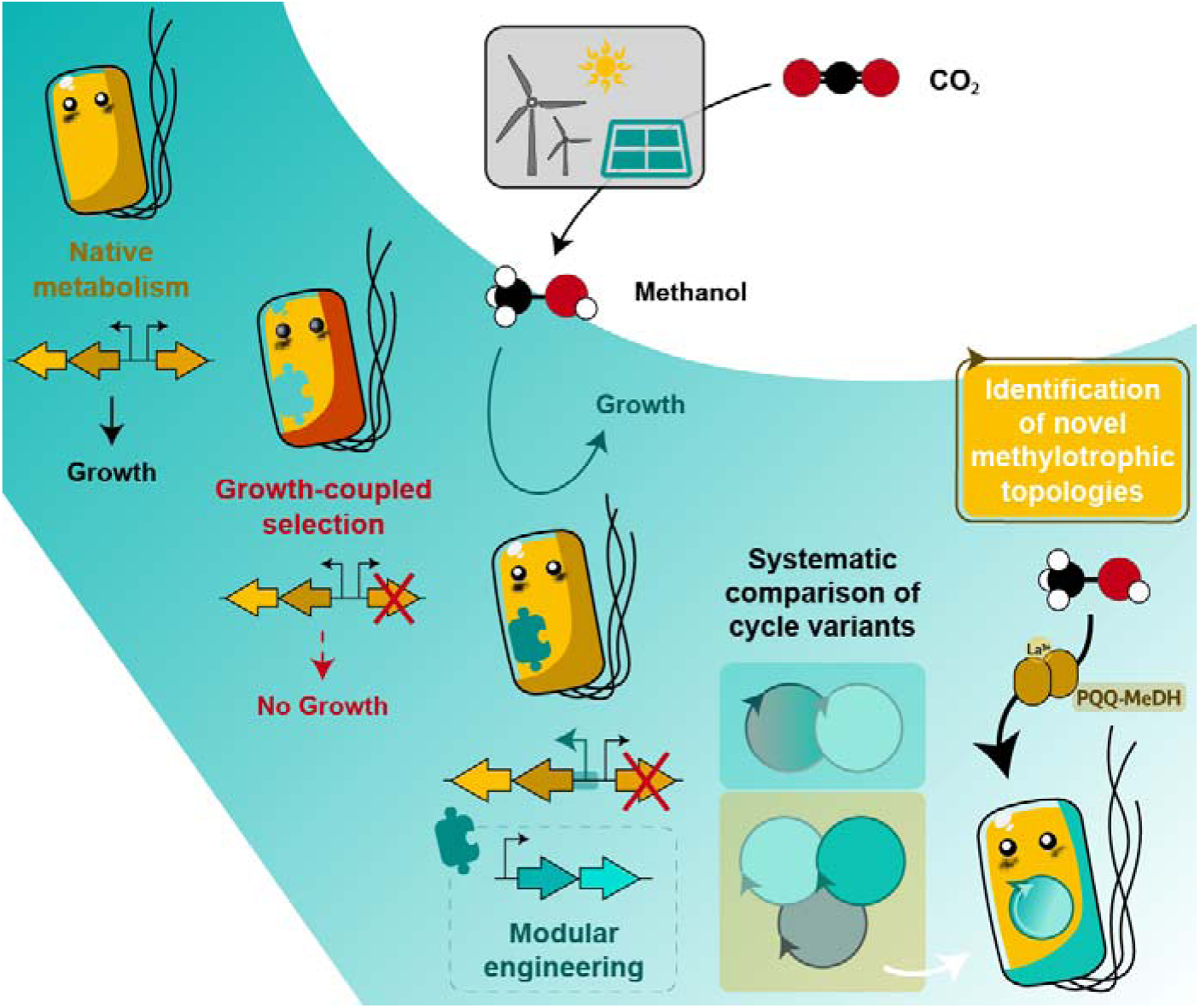

## INTRODUCTION

Climate change, driven by anthropogenic greenhouse gas emissions, poses an existential threat to virtually all species inhabiting planet Earth. Despite recurrent warnings from leading experts, a prevailing economy of overconsumption and affluence supported by inaction of governments and public institutions, especially in developed countries, exacerbated the climate crisis [1]. Mitigating this global crisis requires reducing current emissions, developing innovative CO_2_ capture technologies, and efficiently transitioning to renewable energy sources [2,3]. The urgent need for global decarbonization and rising oil prices have spurred substantial research into alternative, sustainable substrates for industrial production. One-carbon (C_1_) compounds, e.g., methanol and formate, are promising substrates for microbial bioproduction, as they can be derived from methane or CO_2_ using renewable electricity or sunlight [4–9]. However, the model organisms typically used in bioprocesses, e.g., *Escherichia coli* or *Saccharomyces cerevisiae*, do not grow on C_1_ compounds naturally (with the exception of the methylotroph *Komagataella phaffii*) [10]. Methylotrophs, while capable of utilizing C_1_ substrates, are notoriously difficult to metabolically engineer, and current product titers through natural C_1_ assimilation pathways are far from industrial competitiveness [11–13]. The last decade witnessed major breakthroughs to achieve synthetic methylotrophy in a few model bacterial species (*E. coli* and *Cupriavidus necator*) [14-20] owing to the extensive knowledge about their metabolism, established genetic engineering toolsets, and potential for industrial applications [21]. Despite impressive progress in synthetic C_1_ assimilation, designing and implementing efficient pathways for formatotrophy and methylotrophy remains a largely artisanal endeavor. This challenge is further compounded when engineering pathways in non-canonical organisms, for which knowledge of central carbon metabolism is limited.

The soil bacterium *Pseudomonas putida* is a consolidated biotechnological host due to its enhanced tolerance to physicochemical stresses [22–25], availability of tools for genetic and genomic engineering [26], and potential for producing fine chemicals (e.g., new-to-nature products [27–29]) while valorizing waste streams [30]. Although *P. putida* does not naturally assimilate C_1_ feedstocks, we recently showed that methanol, formaldehyde, and formate are oxidized through a set of convergent and peripheral dehydrogenases to generate reducing power [31], and we engineered synthetic formatotrophy in this species through the reductive glycine pathway (rGlyP) [32]. The versatile metabolism of *P*. *putida*, reflecting its extreme adaptability to diverse environments, makes this bacterium an ideal candidate for exploring synthetic methylotrophy, a capability that has yet to be realized. Unlike other bacteria previously engineered for methanol assimilation, *P*. *putida* encodes pyrroloquinoline quinone (PQQ)-dependent alcohol dehydrogenases (MeDHs) [33] that also recognize methanol. However, selecting an adequate set of reactions for methanol assimilation in a non-canonical bacterium is a daunting task. The architecture of the central carbon metabolism in *P*. *putida*, comprising the EDEMP cycle, the tricarboxylic acid (TCA) cycle, and the pyruvate shunt, provides an attractive biochemical template to engineer cyclic C_1_ assimilation pathways [34–36].

The serine cycle (SC) is a natural oxygen-insensitive pathway mediating acetyl-coenzyme A (CoA) synthesis directly from C_1_ molecules (**Fig. 1**a) without carbon loss [37–39]. It was the first pathway identified in methylotrophic bacteria for methanol-dependent growth, yet the last to be fully characterized [40]. Several features make the SC a prime target for engineering C_1_ assimilation in strictly aerobic heterotrophs (e.g., *P*. *putida*), including (i) autocatalytic nature, which could support substantial growth, (ii) operation at ambient CO_2_ conditions, (iii) oxygen tolerance, and (iv) small number of heterologous genes required to establish the full cycle [41]. Additionally, a theoretical study concluded that the SC displays high energetic efficiency [42]. However, compared to other C_1_ assimilation pathways, the SC is ATP- and NADH-demanding, yields the toxic intermediate 3-hydroxypyruvate [43], and requires some reactions (e.g., malyl-CoA synthetase and lyase) that could interfere with the native metabolism of the host [44]. Synthetic SC variants have been designed to overcome these hurdles (**Fig. 1**b–d). These include the homoserine cycle (HSC), the most energy-efficient but requiring assimilation of two formaldehyde molecules (**Fig. 1**b); the modified serine cycle (mSC), encompassing fewer enzymes and consuming less reducing power (**Fig. 1**c); and the serine-threonine cycle (STC) that, in spite of requiring more energy (**Fig. 1**d), highly resembles the CCM of most model organisms and could sustain formatotrophy in *E. coli* upon extended adaptive laboratory evolution (ALE) [41,45,46]. These three variants require less ATP and NAD(P)H than the natural cycle and bypass 3-hydroxypyruvate. However promising, these synthetic cycles have been only implemented in *E*. *coli* to different degrees of completion [45–47]. A concerted effort to systematically compare their operativity is missing, essential for guiding the engineering of non-canonical heterotrophs into methylotrophs.

**Figure 1.**
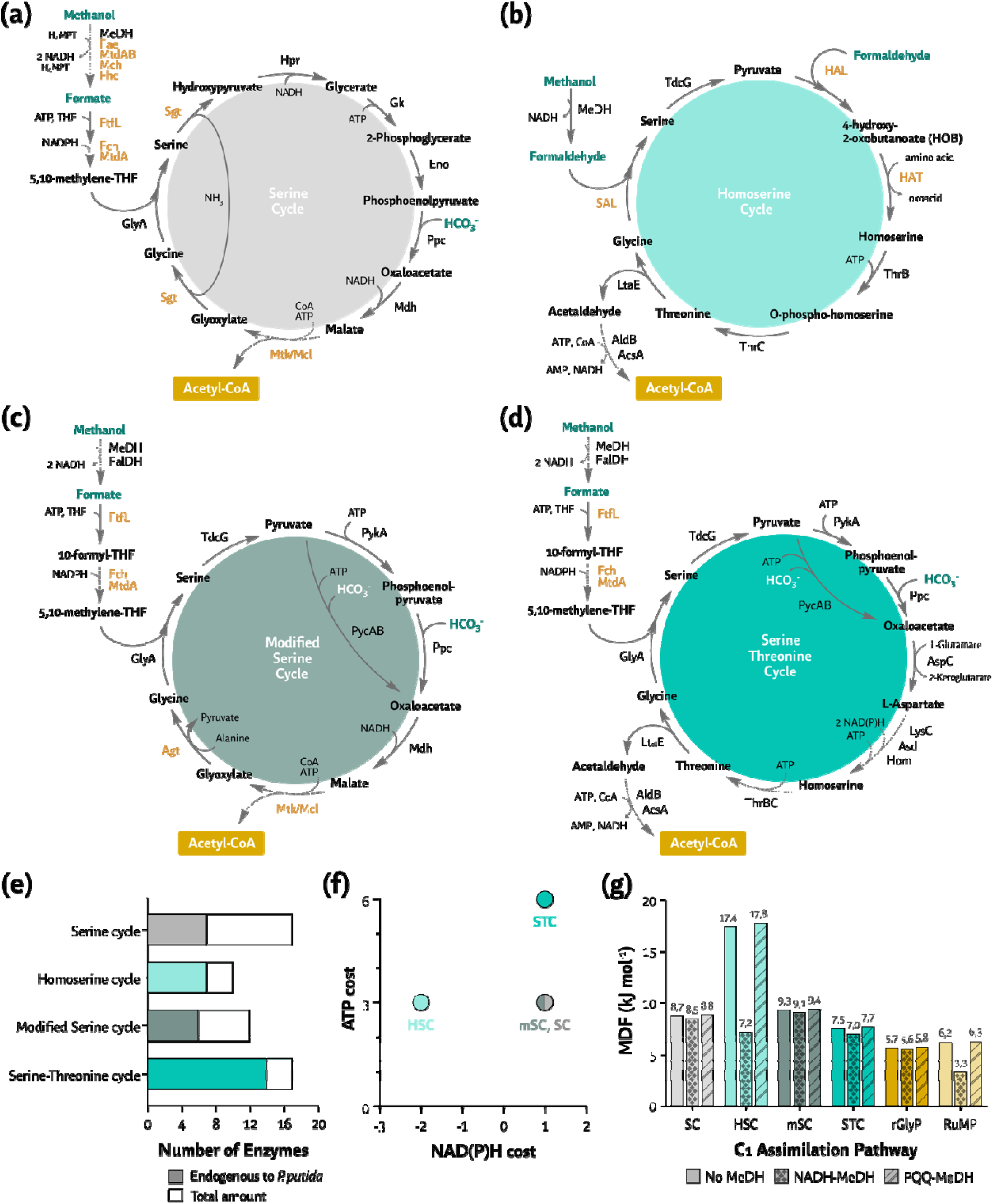
The native serine cycle and its synthetic variants for methanol assimilation. The (a) native serine cycle (SC) found in methylotrophic bacteria has been engineered as (b) the homoserine cycle (HSC), (c) the modified serine cycle (mSC), and (d) the serine-threonine cycle (STC). For enzyme abbreviations, see **Table S1**. Dashed lines represent pooled reactions; orange-colored enzymes represent reactions exogenous to *P*. *putida*. (e) Number of enzymes (or reactions) [endogenous as darker colors and total amount as lighter and darker combined in each cycle] and (f) NAD(P)H and ATP costs calculated from methanol to acetyl-CoA *via* the shortest route (PycAB instead of PykA and Ppc) in the metabolic context of *P*. *putida*. One NADH equivalent has been factored to replenish the amino donor for each transamination event in the three modified serine cycles. (g) Max-min driving force (MDF) of the SC, HSC, mSC, and STC in comparison to the reductive glycine pathway (rGlyP) and ribulose monophosphate (RuMP) pathway. MDF were calculated with eQuilibrator-API [89] with and without NADH- or PQQ-dependent methanol dehydrogenases (no MeDH *versus* NADH-MeDH or PQQ-MeDH). No MeDH indicates the usage of formaldehyde as a substrate, thereby bypassing the thermodynamic constraints of methanol oxidation (especially with NAD-MeDH). Details on MDF calculations are provided in **Table S2**.

To address these challenges, in this work we implemented all synthetic variants of the SC in *P. putida* for methanol assimilation using modular engineering and growth-coupled selection. By dividing the three synthetic SC designs into functional modules, we systematically compared their performance to identify the most promising variant for *in vivo* implementation. Furthermore, we demonstrate that key extant biochemical activities, e.g., PQQ-dependent dehydrogenases and robust gluconeogenesis, can be harnessed for rational engineering of methylotrophy. This recursive rewiring of synthetic and native activities uncovered novel metabolic topologies, dubbed enhanced STC (eSTC), for efficient methanol utilization, providing a blueprint for engineering C_1_ assimilation in non-model heterotrophic bacteria.

## RESULTS

### *In silico* analysis highlights the relevance of serine cycles for efficient methanol conversion

To investigate the potential of all SC variants for synthetic methylotrophy in *P*. *putida*, we examined the pathway demands in terms of enzymes requirements (**Fig. 1**e, **Table S1**), ATP and NAD(P)H costs (**Fig. 1**f), thermodynamics under physiological conditions, using maximum-minimum driving force (MDF) analysis (**Note S1**), and predicted specific growth rates (μ) using flux balance analysis (FBA; **Fig. 1**g, **Tables S2**-**S3**). We compared the SC variants with each other and other pathways engineered for methanol assimilation, i.e., the rGlyP [14,32,48] and the ribulose monophosphate (RuMP) pathway [49,50]. All six pathways were modelled in the biochemical context of *P. putida* KT2440 (**Fig. 2**), considering either methanol or formaldehyde as C_1_ substrate. For methanol oxidation, either NADH- or PQQ-dependent MeDHs (with Δ_r_G′ = 34.2 and −24.8 kJ mol^−1^, respectively) were included in the simulation (**Fig. 1**g and **Table S2**).

**Figure 2.**
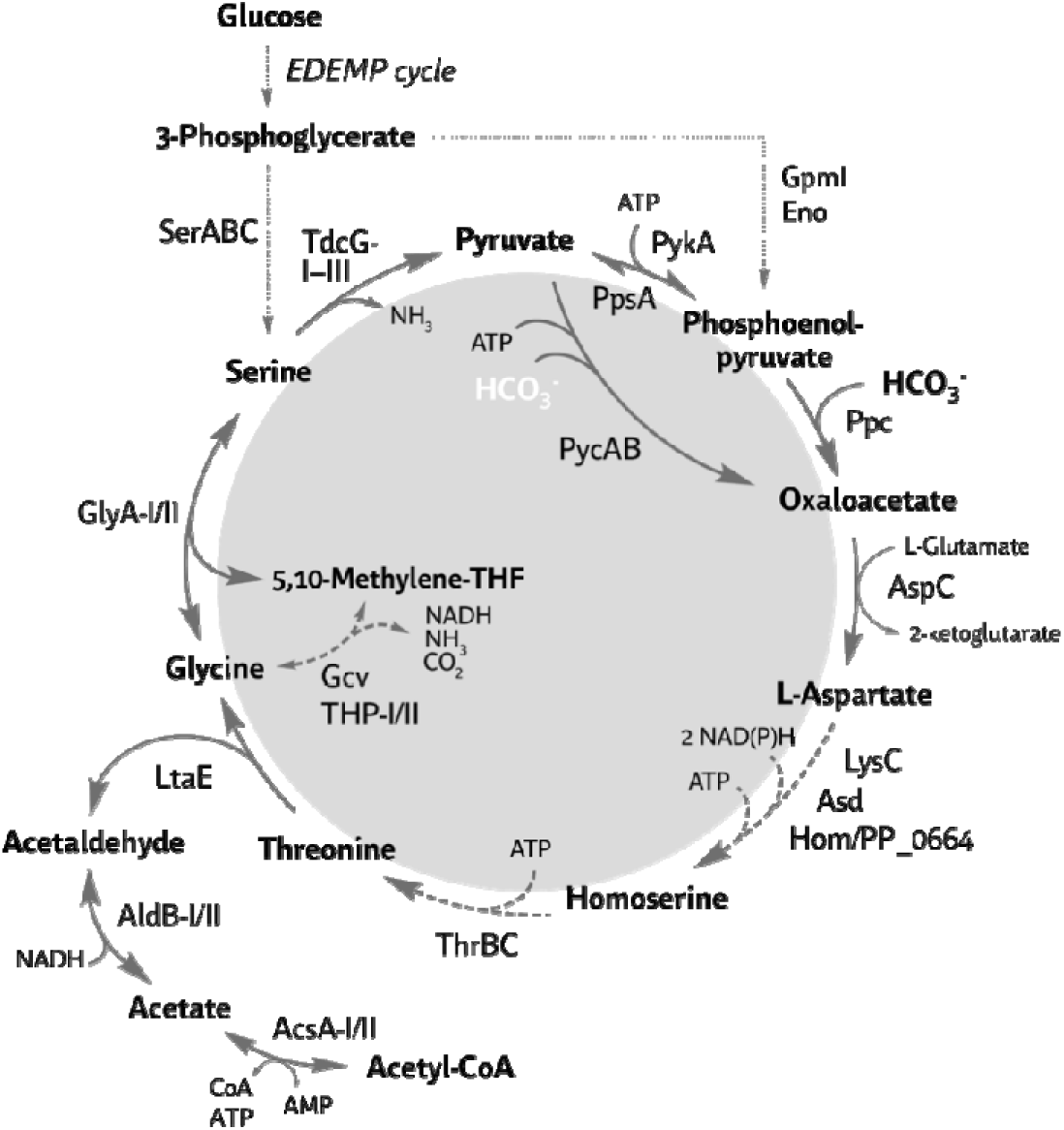
Native metabolism of *P. putida* growing on glucose. The diagram shows the reactions around the amino acids serine, glycine, and threonine, critical for the serine cycles in this study. Enzyme abbreviations are indicated in **Table S1**.

The simulations showed a broader, favorable thermodynamic range for all SC variants (MDFs ∼7.5-9 kJ mol^−1^) compared to the rGlyP and RuMP pathways (MDFs ∼6-7 kJ mol^−1^). The HSC was the most thermodynamically favorable variant, whether using formaldehyde (i.e. No MeDH in **Fig. 1**g) or methanol with PQQ-dependent MeDHs, surpassing all other pathways by at least 2-fold (**Fig. 1**g and **Table S2**). Due to decreased alcohol oxidation performance by NADH-dependent MeDH, some pathways (e.g., the HSC or RuMP pathway) appeared less feasible. We also used MDF to identify thermodynamic bottlenecks in the SC variants (**Fig. S1**). The primary bottleneck for all routes was predicted to be the condensation of glycine and 5,10-methylene-tetrahydrofolate (5,10-methylene-THF). For the HSC, the condensation of pyruvate and formaldehyde was the main thermodynamic barrier. Replenishing glycine from C_3_ intermediates was another bottleneck for the STC, with oxaloacetate transamination and subsequent reactions being thermodynamically unfavorable (**Fig. 1**d and **Fig. S1**). These biochemical bottlenecks were addressed in designing the functional modules for synthetic C_1_ assimilation.

Next, FBA was used to investigate intrinsic differences among the SC variants to support C_1_-trophic growth (**Tables S3-S4**). To this end, we curated the latest genome-scale metabolic model of *P*. *putida* KT2440 [51,52] and added the synthetic reactions for C_1_ assimilation in each pathway [53]. The natural SC and the mSC, with similar energy and NAD(P)H demand (**Fig. 1**e–f), predicted similar specific growth rates that correlated with their favorable thermodynamics (**Table S3**). Formate oxidation (i.e., *metabolic turnover*, with CO_2_ emission as proxy) reflected the energetic demand of the corresponding cycle variant, as formate oxidation was only required for NAD(P)H generation. The STC supported the slowest growth among all variants tested, reflecting its unfavorable thermodynamics. Furthermore, the HSC, as the most energy-efficient variant, surpassed the performance of the natural SC (**Table S3**). These results indicate that engineering the SC and its synthetic variants in *P*. *putida* is not only possible but also favored by the presence of native PQQ-dependent alcohol oxidation that could facilitate methylotrophy. A detailed FBA comparison of individual modules is discussed in the corresponding sections below.

### Growth-coupled selection schemes and modular testing of serine cycles *P.inputida*

To standardize the comparison of each SC, we employed growth-coupled selections [54] and modular engineering in strain SEM11 [55], a genome-reduced derivative of wild-type *P. putida* KT2440 [56]. We first identified structural commonalities among the three SC variants to generate different functional modules (**Fig. 3**). Module 1 (M_1_) encompasses the assimilation of a C_1_ moiety from glycine to serine, whereas module 2 (M_2_) closes the cycle by transforming pyruvate back to glycine, including another round of carbon assimilation (**Fig. 3**). Serine and glycine auxotrophs were designed to test C_1_ assimilation by M_1_, while serine, acetyl-CoA, or homoserine auxotrophies were implemented to select M_2_. All selection schemes were designed *in silico*, utilizing FBA to explore the long-term stability of the metabolic interventions prior to engineering efforts (**Table S5**). A detailed roadmap of all modifications implemented in *P*. *putida* SEM11, including gene deletions, gene insertions, and overexpression of genes from the chromosome or in plasmids, is presented in **Fig. S2**.

**Figure 3.**
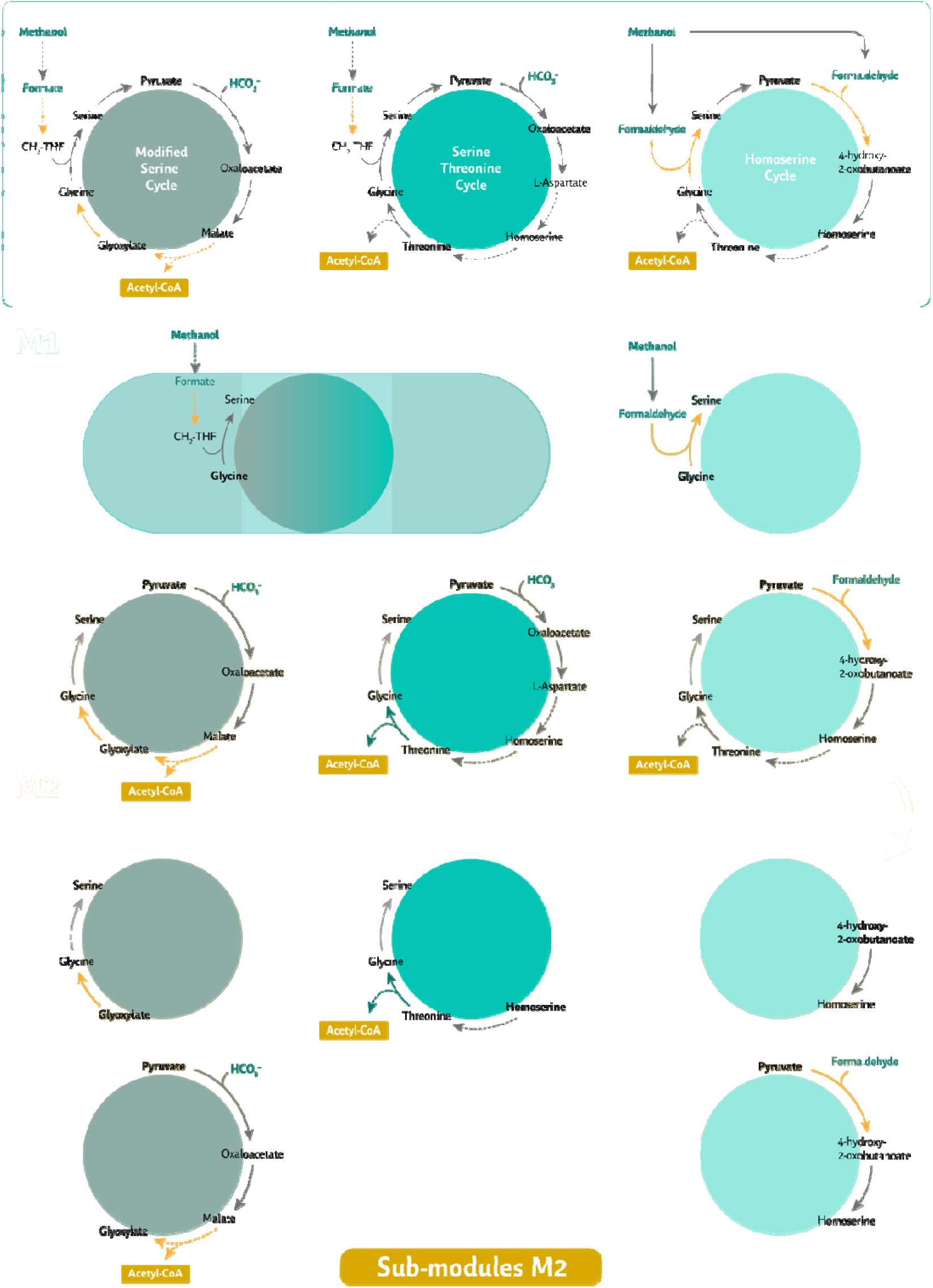
Metabolic maps for the three modified serine cycles implemented for synthetic C_1_ assimilation. The modified serine cycle (mSC, left), the serine-threonine cycle (STC, middle), and the homoserine cycle (HSC, right) have been split into two modules. Module 1 (M_1_) comprises the assimilation from glycine to serine, and Module 2 (M_2_) involves the production of glycine from pyruvate. M_2_ was also split into sub-modules, which are represented in the lower part of the figure. All reactions and their cognate genes are listed in **Table S1**. Dashed lines represent pooled reactions; yellow-colored arrows represent reactions exogenous to *P. putida*.

### Designing, building, and testing M_1_ — Formate and formaldehyde assimilation inLt-oserine

The assimilation of C_1_ moieties *via* THF requires expression of *ftfL*(formate-THF ligase), *fch* (methenyl-THF cyclohydrolase), and *mtdA* (5,10-methylene-THF dehydrogenase) from *Methylobacterium extorquens* AM1, producing 5,10-methylene-THF [32]. M_1_ requires 1 NADPH and 1 ATP for each formate assimilated as part of the mSC and the STC (**Fig. 1**c–d and **Fig. 3**). Threonine aldolases can utilize formaldehyde for glycine aldolization to serine (serine aldolase), required for the HSC [20], in an ATP and NAD(P)H-independent fashion. Since methanol (rather than formate) was our target substrate, MeDH genes should be overexpressed to provide formaldehyde, generating one reducing equivalent (NADH or PQQH_2_) per methanol oxidized. Endogenous oxidation activities in *P*. *putida* [31] catalyze the subsequent oxidation of formaldehyde to formate, as required for the mSC or STC. *In silico* analysis indicated that M_1_ could support methylotrophic growth *via* the mSC/STC and HSC configurations with similar performance with glucose as cosubstrate (**Tables S6** and **S7**). At high methanol uptake rates, M_1_ of the HSC (henceforth referred to as M_1_·HSC) was predicted to outperform the M_1_·mSC/STC (**Table S4**). Based on these designs, we constructed a strain that can support C1 assimilation through M_1_·HSC.

### A low specificity l-threonine aldolase [_1_M·HSC] restores growth of a serine and glycine auxotroph

We constructed a serine and glycine auxotroph ( serA glyA) to select a l-serine aldolase activity, further deleting *frmAC* (*PP_1616-7*) to avoid formaldehyde detoxification at low methanol concentrations [31]. The resulting strain (*P*. putida serA glyA frmAC) was transformed with a low-copy-number plasmid (pSEVA621 [57,58]) constitutively expressing ^Ec^*ltaE* and the CT4-1 methanol dehydrogenase gene from *C. necator*(^Cn^*mdh*), engineered to display enhanced activity [59]. Control plasmids harboring only ^Ec^*ltaE* or ^Cn^*mdh* were also tested (**Fig. 4**b). The serine pool in the serA glyA frmAC strain incubated in de Bont minimal (DBM) medium with 20 mM glucose, 10 mM glycine, and 250-500 mM methanol was replenished, relieving the auxotrophy, when ^Ec^*ltaE and*^Cn^*mdh* were overexpressed (**Fig. 4**), supporting μ ∼ 0.3 h^−1^ (**Fig. 4**c). As expected, plasmid pSEVA621·^Cn^*mdh* alone did not revert the auxotrophy. Expressing ^Ec^*ltaE* alone supported growth when serine was supplied (**Fig. 4**b), and we concluded that ^Ec^LtaE can cleave serine to glycine and formaldehyde. Furthermore, overexpressing ^Ec^*ltaE* alone promoted growth at high methanol concentrations (**Fig. 4**b– c), which we attribute to several endogenous, broad-range alcohol dehydrogenases acting on methanol at > 250-500 mM [31]. Encouraged by the activity of ^Ec^LtaE as l-serine aldolase, we assessed whether LtaEs from other species could provide higher catalytic activities.

**Figure 4.**
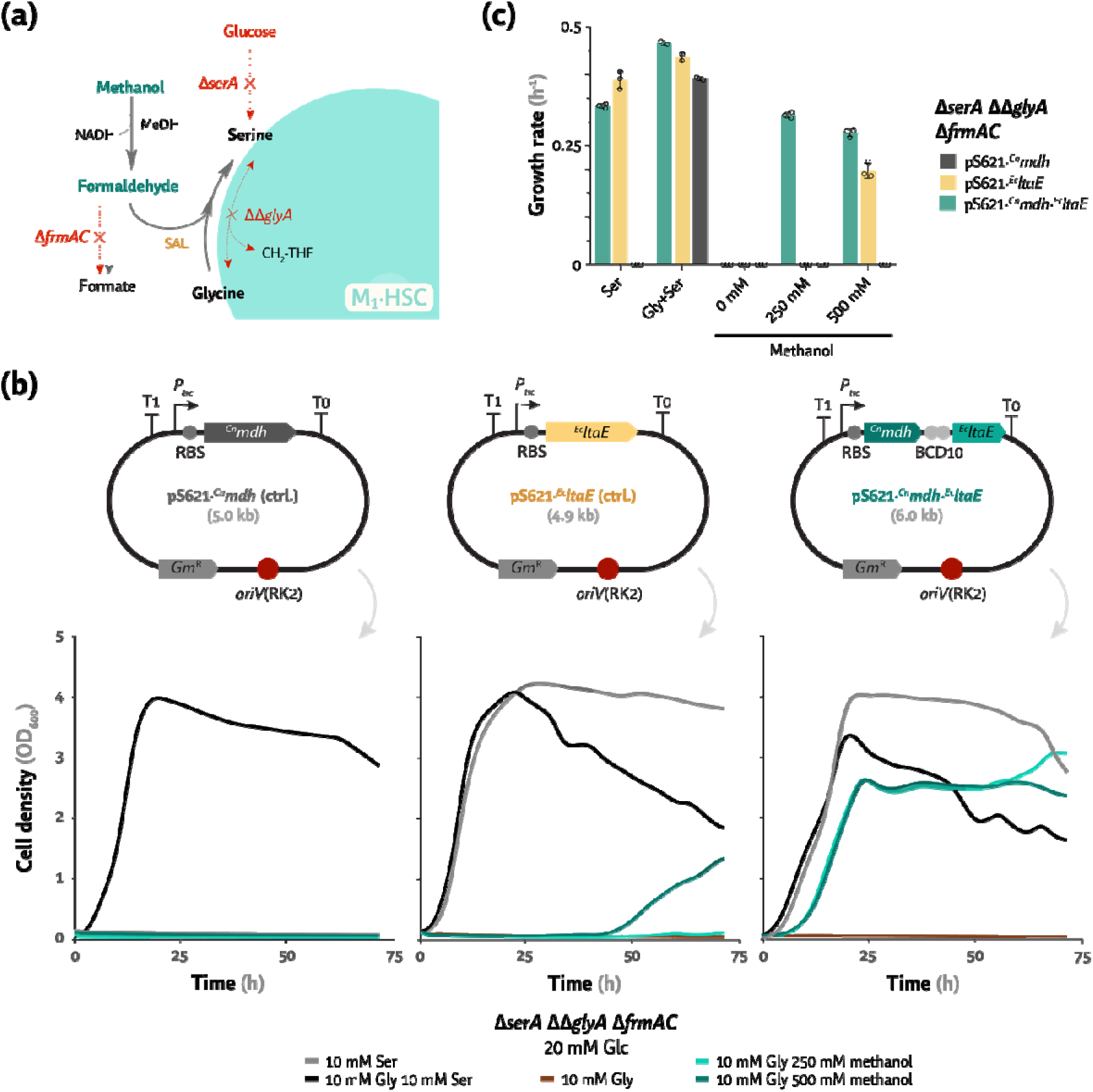
Testing M_1_ of the homoserine cycle in *P. putida*. M_1_·HSC relies on the formaldehyde-assimilative l-serine aldolase (SAL) reaction via the side-reaction of LtaE from E. coli. (a) The auxotrophic strain serA glyA frmAC was employed to test the capacity of ^Ec^ltaE to assimilate formaldehyde and glycine into serine. (b) Structure of plasmid maps used to relieve the auxotrophy (pS621·^Cn^mdh·^Ec^ltaE), including the control vectors pS621·^Cn^mdh and pS621·^Ec^ltaE, and growth profile of the serA glyA frmAC strain carrying such plasmids in DBM medium supplemented with 20 mM glucose and serine, glycine, or methanol as indicated. A canonical RBS was used (5’-AGG AGG AAA AAC AT-3’) in these constructs. Both (b) cell densities (estimated as the optical density measured at 600 nm, OD_600_) and (c) specific growth rates (μ, h^−1^) are indicated as average values ± standard deviation of three biological replicates. Individual data points are shown whenever relevant.

### ^Ec^LtaE outperforms a library of aldolase variants for formaldehyde assimilation

We used *P*. *putida* serA glyA frmAC as selection strain to test a library of low-specificity l-threonine aldolases, encompassing variants from heterotrophic bacteria (*E. coli, P. putida*, and *Vibrio natriegens*), C_1_-trophic bacteria (*M. extorquens* AM1 and *C. necator*) and yeast (*K. phaffii*), and heterotrophic yeasts (*S. cerevisiae* and *Yarrowia lipolytica*)(**Fig. S3**). The *ltaE* variants were individually expressed in a pSEVA621 backbone alongside ^Cn^*mdh*, and the ΔserA glyA frmAC strain harboring the plasmid library was tested in DBM medium with 20 mM glucose, 10 mM glycine, and 60-500 mM methanol (**Fig. S3**). ^Ec^LtaE outperformed all other variants, and only the enzymes from *E. coli*, *V. natriegens*, and *Y. lipolytica* supported l-serine aldolase activity (**Fig. S3**). Moreover, all LtaE variants executed the reverse aldolase reaction (serine to glycine and formaldehyde), enabling growth when serine was supplied. Hence, ^Ec^LtaE was retained for further engineering SC variants in *P. putida*.

### Rewiring the transcriptional regulation of genes encoding PQQ-dependent, broad-range alcohol dehydrogenases

The growth of the engineered *P*. *putida* strain at high methanol concentrations in the absence of exogenous MeDH*s* expressed (**Fig. 4**) could be attributed to endogenous broad-range alcohol dehydrogenases. This oxidative activity has been previously linked to PedE and PedH, two periplasmic, PQQ-dependent MeDHs that utilize calcium (Ca^2+^) and rare-earth elements (REEs, usually lanthanides, La^3+^) as cofactors, respectively [33]. The expression of the *ped* cluster is tightly regulated and activated by several alcohols and the presence of REEs [60–62]. PedE and PedH exhibit complementary functions depending on cofactor availability: *pedE* is expressed when short-chain organic alcohols are supplemented in the absence of REEs [62], whereas *pedH* is expressed in the presence of La^3+^ (**Fig. 5**b). YiaY (PP_2682) has been recently proposed to act as a transcriptional activator for the *ped* cluster alongside downstream neighboring histidine kinase, *PP_2683* [61]. Together, they resemble the signaling system for ethanol oxidation in *P. aeruginosa*, the two-component system ErcAS [61]. To test if we could use the endogenous MeDHs for synthetic methylotrophy, we investigated the transcriptional regulation of *pedE* and *pedH* in a strain constitutively expressing *yiaY* (**Fig. 5**a). We transformed wild-type *P*. *putida* and its P_yiaY_::P_14G_(BCD10) derivative with reporter plasmids harboring the promoters of *pedE*, *pedH*, and *yiaY* cloned in front of a monomeric superfolder GFP (*msfGFP*) gene (**Fig. 5**c). By recording msfGFP fluorescence under different cultivation regimes, we aimed to understand the regulatory effect of YiaY on the transcription of the *pedE*, *pedH*, and *yiaY* genes.

**Figure 5.**
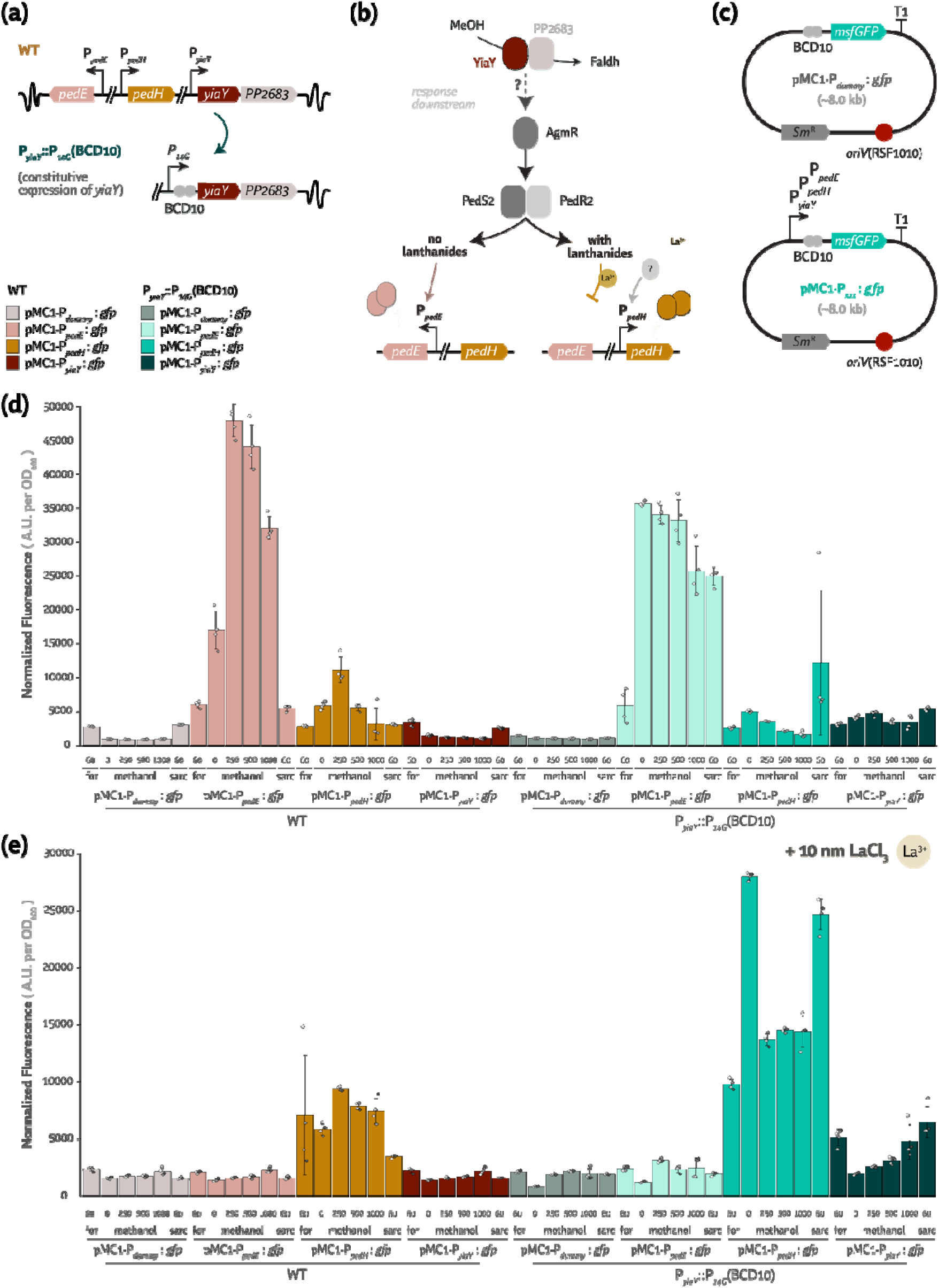
Constitutive expression of native PQQ-dependent alcohol dehydrogenase genes via yiaY overexpression. (a) Organization of the ped cluster, including the two PQQ-dependent MeDHs pedE and pedH, a putative NAD-MeDH, and the yiaY regulator with its downstream neighbor PP_2683. Constitutive expression of yiaY was achieved by promoter engineering. (b) Simplified regulatory model scheme for the ped cluster based on Wehrmann et al. [62] and Bator et al. [61], including the potential regulatory function of YiaY as a methanol sensor. Downstream in the signaling cascade, the presence of rare-earth elements (REEs, e.g., lanthanide salts, La^3+^) modulates the activity of PedE and PedH. (c) Biosensor plasmids, harboring a monomeric superfolder GFP gene (msfGFP) expressed by the native promoters of yiaY, pedE or pedH, or a dummy promoter (negative control). (d) Normalized fluorescence plots (arbitrary units, A.U., normalized to the optical density at 600 nm, OD_600_) for strains EM42 P_yiaY_::P_14G_(BCD10), with yiaY overexpression, or the wild-type (WT) control, harboring the biosensor plasmids shown in panel (c). All strains were grown in DBM medium with 20 mM glucose supplemented with formate (for), methanol, or sarcosine (sarc, used as a formaldehyde donor). (e) Same as in panel (d), but with the additional supplementation of LaCl_3_ at 10 nM. Average values for normalized fluorescence ± 95% confidence intervals of at least three biological replicates are represented in all cases, together with individual data points. Statistical analysis indicated P-values < 0.05 for all pairwise comparisons where error bars do not overlap.

The reporter strains were grown in DBM medium with 20 mM glucose and varying concentrations of C_1_ molecules (i.e., formate, methanol, or formaldehyde derived from sarcosine) in the presence or absence of LaCl_3_ as REE (**Fig. 5**d–e). In the wild-type strain, *pedE* expression strongly depended on the methanol or formaldehyde (sarcosine) concentration when REEs were absent (**Fig. 5**d). However, *pedE* was expressed at high levels with constitutive *yiaY* expression, regardless of the methanol or formaldehyde concentration (**Fig. 5**d). A similar trend was observed for *pedH* in the presence of 10 nM LaCl_3_, where constitutive *yiaY* expression resulted in a 1.5- to 3-fold increase in *pedH* expression with varying methanol or formaldehyde levels (**Fig. 5**e). Hence, YiaY (and possibly PP_2683) orchestrates a transcriptional response that regulates PQQ-MeDHs in a C_1_ substrate-dependent manner, overcoming the regulatory effect of REEs. Hence, constitutively overexpressing *yiaY* upregulated PQQ-MeDH– encoding genes independently of the C_1_ substrate, which provides a basis for further engineering efforts.

### Endogenous PQQ-dependent methanol dehydrogenases provide formaldehyd_1_·eST[MC·mSC]

To implement M_1_ of the STC, shared with the mSC (**Fig. 1**c–d), we integrated the C_1_ assimilation module in the *phaC1ZC2DFI* locus of strain SEM11, resulting in *P*. *putida* serA gcvTHP *pha::ftfL-mtdA-fch*(termed ΔSGG-C_1_; **Fig. 6**a). In initial experiments, we assessed if the pool of methylated THF and, subsequently, serine could be restored in strain ΔSGG-C_1_ incubated in DBM medium with 20 mM glucose, 10 mM glycine, and varying formate concentrations (**Fig. 6**b). This was indeed the case, with specific growth rates and maximum cell density (optical density at 600 nm, OD_600_) of μ ∼ 0.6 h^−1^ and 2.5-3, respectively; similar to the values observed when supplementing glycine and serine (positive control, **Fig. 6**c).

**Figure 6.**
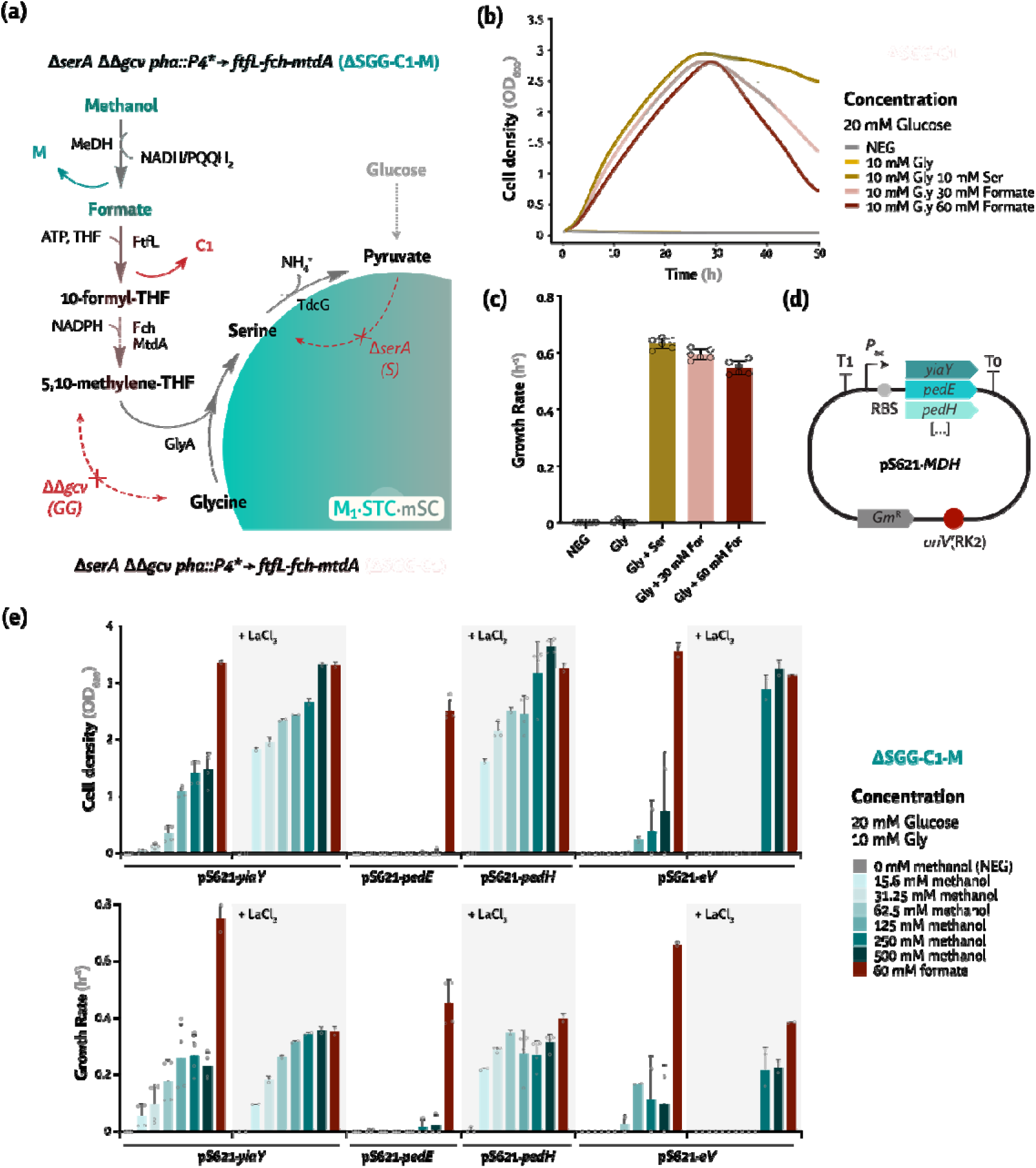
Testing the assimilative M_1_ of the serine-threonine and modified serine cycles via the C_1_ moiety-carrier molecule tetrahydrofolate (THF). (a) The auxotrophic serA gcvTHP strain (termed ΔSGG) was employed to test one-carbon (shown as C_1_) assimilation via FtfL, Fch, and MtdA. Such assimilation pathway was introduced in the genome at the phaC1ZC2DFI locus under the expression of a strong P_4_ promoter (indicated as P_4_*). Methanol oxidation was facilitated by plasmid-based overexpression of different endogenous methanol dehydrogenases (MeDH, indicated as M). (b) Growth profile of strain ΔSGG-C_1_ in DBM medium supplemented with 20 mM glucose, 10 mM glycine and, when appropriate, 10 mM serine (positive control) or 30-60 mM formate. (c) Specific growth rates for the experiments shown in panel (b). (d) Plasmid maps for overexpressing methanol dehydrogenases (MeDH) genes in the ΔSGG-C_1_ strain (yielding ΔSGG-C_1_-M). (e) Maximum cell density (optical density measured at 600 nm, OD_600_) and specific growth rates of strain ΔSGG-C_1_-M with MeDH overexpression (yiaY, pedE, and pedH) compared to an empty vector (eV). Strains were grown in DBM medium with 20 mM glucose and 10 mM glycine, as well as varying concentrations of methanol, or 60 mM formate as a positive control [as shown in panel (b)]. When indicated, 10 nM LaCl_3_ was tested as a cofactor for the PQQ-dependent alcohol dehydrogenase PedH. Average values for cell densities (OD_600_) and specific growth rates (μ, in h^−1^) ± standard deviation of three biological replicates are represented in all cases; individual data points are shown whenever relevant.

A library of native and heterologous MeDH genes, constitutively expressed from a pSEVA621 backbone as explained in the previous section, was tested in strain ΔSGG-C_1_ to explore synthetic methylotrophy at different substrate levels (**Fig. 6**d, **Fig. S4** d). Incubating these strains in DBM medium with 20 mM glucose, 10 mM glycine, and 15-500 mM methanol revealed that NADH-dependent MeDHs, e.g., ^Bs^Mdh, ^Cn^Mdh, ^Bm^Mdh2, or the native AdhP had no activity below 125 mM methanol, compatible with previous reports [63–66]. Moreover, strain ΔSGG-C_1_ harboring these NADH-dependent MeDHs grew slowly, with μ < 0.25 h^−1^ (**Fig. S4**d). In contrast, overexpressing *xoxFGJ*, encoding a PQQ-dependent MeDH from *M. extorquens* AM1 [67], resulted in higher cell densities (OD_600_ ∼ 3) at low methanol concentrations, with similar μ values as supported by NADH-MeDHs (**Fig. S4**).

The native PQQ-dependent, broad-range alcohol dehydrogenases [33] were also tested (**Fig. 6**d–e). We overexpressed *pedE*, *pedH*, and *yiaY* from a plasmid in strain ΔSGG-C_1_ (**Fig. 6**d–e), incubated in DBM medium with 20 mM glucose, 10 mM glycine, and 15-500 mM methanol. Strains overproducing PedH and YiaY outperformed all NADH-dependent MeDHs (**Fig. S4**) in the presence of LaCl_3_, with μ ∼ 0.35 h^−1^ and reaching OD_600_ > 3 (**Fig. 6**e). Whilst PedE did not promote growth, alcohol oxidation in the strain carrying the empty vector occurred at high methanol concentrations, especially when lanthanides were present. Without added LaCl_3_, *yiaY* overexpression still promoted similar specific growth rates but the cell density was lower (OD_600_ ∼ 1.5, **Fig. 6**e), indicative of *pedE* upregulation and *pedH* repression [33]. These experiments confirm that sufficient methanol oxidation to formaldehyde can be achieved through the endogenous PQQ-dependent dehydrogenases of *P. putida viayiaY* overexpression—even more efficiently than with NADH-dependent MeDHs, typically used in metabolic engineering of C_1_ assimilation.

### Designing, building, and testing M_2_ — C_3_ carboxylation and replenishment of the glycine pool by C_4_ cleavage

Given that l-serine deamination to pyruvate is an activity native to *P. putida*, we focused on implementing M_2_ for converting pyruvate to glycine. Serine or glycine auxotrophs (Δ*serA* or ΔΔ*glyA*) were used for selection, yet the glycine auxotroph had growth defects, and we adopted the serine auxotroph for testing the activity of M_2_. Glycine, formed from pyruvate by M_2_, is used for l-serine synthesis *via* the endogenous GcvTHP and GlyA activities. Hence, a serine auxotroph would only grow if M_2_ provides glycine from C_1_ assimilation. Also, carbon assimilation relies on pyruvate condensation with a C_1_ moiety. In the mSC and STC, this carboxylation could proceed with HCO_3_^−^ *via* the native pyruvate carboxylase (PycAB) and/or PEP carboxylase (Ppc). In the HSC, in contrast, a non-native reaction condenses pyruvate with formaldehyde to produce 4-hydroxy-2-oxobutanoate (HOB, **Fig. 1**b–d, **Fig. 3**). HOB is transaminated to homoserine, subsequently phosphorylated and hydrated to l-threonine (**Fig. 1**b, **Fig. 3**). This amino acid is broken down into glycine and acetaldehyde, and the C_2_ pool is used for growth *via* activation to acetyl-CoA. M_2_ of the HSC requires 3 ATP per acetyl-CoA formed.

The STC partly relies on the same circularization as indicated for the HSC (i.e., homoserine to glycine), except that homoserine is produced *via* l-aspartate, stemming from oxaloacetate. This pathway is endogenous to *P. putida* yet has a high energy and reducing power cost, with a total of 5 ATP and 2 NAD(P)H per acetyl-CoA formed (**Fig. 1**d, **Fig. 3**). The mSC resembles the natural SC, where PEP (or pyruvate) is carboxylated to oxalacetate and reduced to malate, reversing the TCA cycle. Malate is then activated to malyl-CoA and broken down into glyoxylate and acetyl-CoA (**Fig. 1**c, **Fig. 3**). While acetyl-CoA is assimilated into biomass, glyoxylate replenishes glycine by transamination. This M_2_ requires 2 NADH, but only 2 ATP per acetyl-CoA formed.

Our FBA simulations indicated that C_3_ carboxylation and glycine replenishment *via* the different M_2_ configurations could occur efficiently in all three SC designs (**Tables S6**-**S7**), with the mSC outperforming the STC and HSC. The predicted growth rate for the HSC was the slowest among the cycles due to enforcement of the homoserine auxotrophy (i.e., blocking anaplerotic pathways) that compromises glycine regeneration if intermediates within the module are drained to support growth (e.g., homoserine or l-threonine; **Tables S3**-**S4**). The next step was to compare the three module variants *in vivo* to regenerate glycine from the C_3_ pool.

### Glycine from the C_3_ pool via the endogenous serine-threonine cycle module [_2_M·STC]

Theoretically, *P. putida* could support the STC *via* oxaloacetate (carboxylating the C_3_ pool, e.g., pyruvate or PEP from the EDEMP cycle), l-aspartate, homoserine, l-threonine, and glycine *via* the endogenous LtaE. However, the serA or glyA auxotrophs could not grow in DBM medium with glucose, indicating that this pathway connectivity is either inactive or does not carry enough flux to support growth. Hence, we initially overexpressed the native l-threonine aldolase (^Pp^LtaE), encoding the final step in the cycle, in the serA or glyA strain (**Fig. 7**a). The native LtaE was unable to replenish the pool of serine (**Fig. 7**b–c) or glycine (**Fig. 7**d) at the endogenous expression levels in DBM medium with 20 mM glucose. Growth was restored when supplemented with 10 mM l-threonine or homoserine. Overexpressing ^Pp^*ltaE* supported growth of both Δ*serA* and ΔΔ*glyA* auxotrophs under all cultivation conditions (**Fig. 7**a–e). *P*. *putida* SEM11Δ*serA* overexpressing ^Pp^*ltaE* grew at μ ∼ 0.25 h^−1^ and reached OD_600_ ∼ 2 (**Fig. 7**b–c), while *P*. *putida*ΔΔ*glyA* transformed with the same plasmid grew as fast but reached half the maximum OD_600_. Hence, overexpression of the endogenous ^Pp^*ltaE* proved sufficient to promote flux from the C_3_ pool to glycine through M_2_·STC.

**Figure 7.**
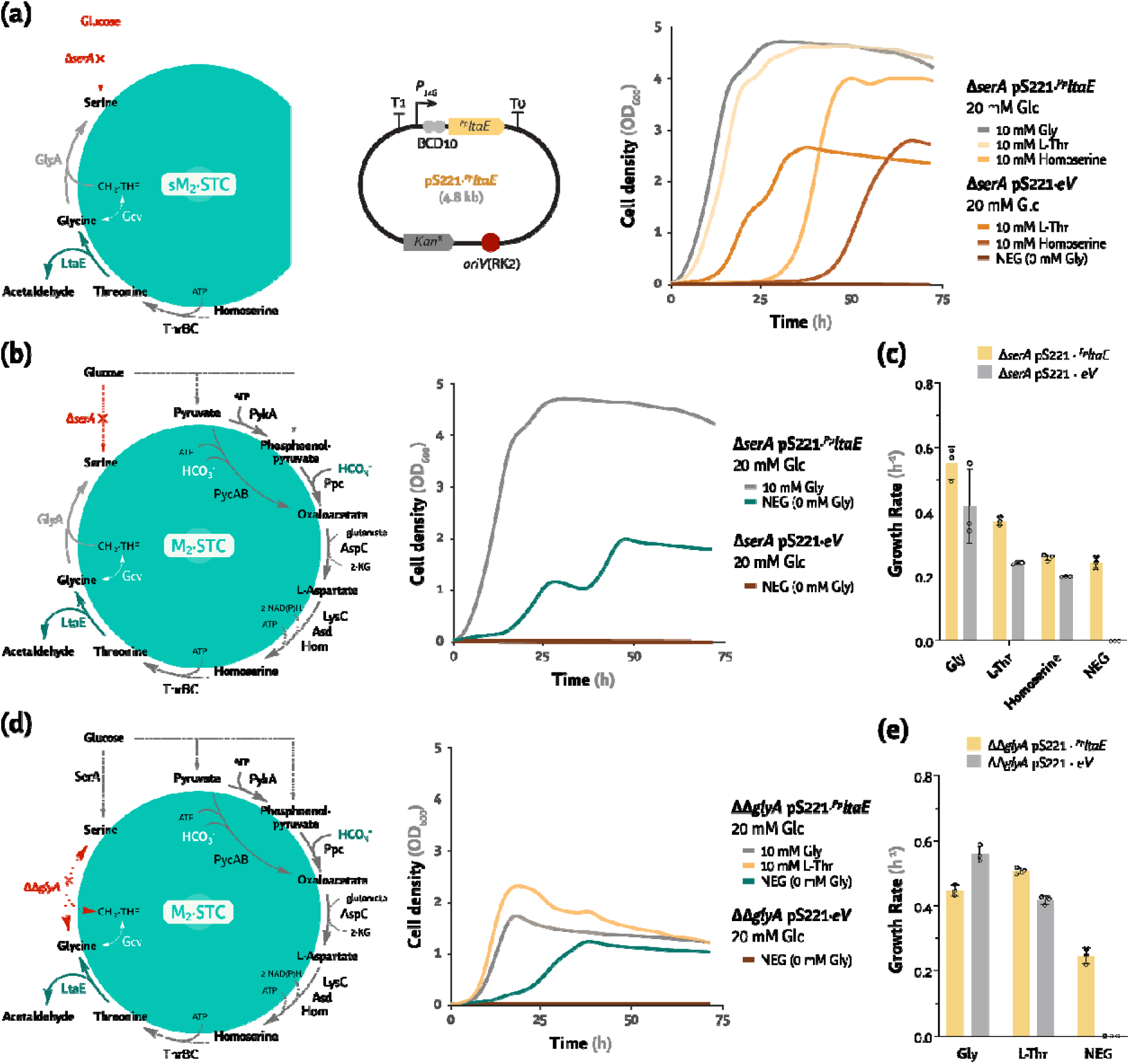
Testing M_2_ of the serine-threonine cycle to yield glycine or serine from the C_3_ and C_4_ pool. (a) Growth of a ΔserA mutant in DBM medium supplemented with 20 mM glucose to test serine production from glycine, l-threonine, and homoserine, with or without constitutive overexpression of ^Pp^ltaE in a pSEVA221 vector (pS221), compared to an empty vector (eV). (b) Testing the same conditions as in panel (a) but directly from glucose, which yields pyruvate or PEP. (c) Specific growth rates corresponding to the experiments plotted in panels (a) and (b). (d) Growth of a ΔΔglyA mutant in DBM medium supplemented with 20 mM glucose to test glycine production from l-threonine or glucose, with or without constitutive overexpression of ^Pp^ltaE compared to an empty vector (eV). (e) Specific growth rates corresponding to the experiments plotted in panel (d). Average values for bacterial growth (estimated as the optical density measured at 600 nm, OD_600_), and specific growth rate (μ, in h^−1^) ± standard deviation of three biological replicates are represented in all cases. Individual data points are shown in all cases. Except for glycine supplementation in panel (c), all comparisons were statistically significant with P-values < 0.01.

### HOB transaminase occurs endogenously inP. putida [M_2_·HSC]

M_2_ of the HSC produces HOB *via* a HOB aldolase (HAL), followed by transamination to homoserine by a HOB transaminase (HAT) (**Fig. 1**b, **Fig. 3**, **Fig. 8**). Homoserine is converted into glycine *via* l-threonine, yielding acetyl-CoA in the same manner as within the STC (**Fig. 7**). To test HAT and HAL activities, we constructed a homoserine auxotroph (**Fig. 8**a, **Fig. S5**). Rather than merely eliminating Asd (aspartate-semialdehyde dehydrogenase, which requires diaminopimelate supplementation [20]), we deleted *hom* and *PP_0664* (encoding homoserine dehydrogenase). The resulting *P*. *putida* strain, Δ*hom*Δ*PP_0664*, was auxotrophic for homoserine. With this selection strain, we tested whether HOB transaminase activity is endogenous to *P. putida*. We screened a set of genes encoding putative HATs (*aspC*, *alaC*^A142P Y275D^, and *ilvE* from *E. coli*; and *ilvE* from *P. putida* KT2440) constitutively expressed in a pSEVA221 vector as explained in the previous sections (**Fig. S5**a). This HAT library was introduced in *P*. *putida* SEM11 Δ*hom*Δ*PP_0664*, and the cells were grown in DBM medium with 20 mM glucose, supplemented with 2 mM homoserine or HOB (**Fig. S5**b). None of the overexpressed putative HAT genes supported a substantial increase in growth parameters, except for a slightly higher μ in cells transformed with the ^Ec^*aspC* or ^Ec^*ilvE* constructs (**Fig. S5**c). Moreover, overexpression of either *E. coli* gene was detrimental when homoserine was supplemented (**Fig. S5**b–c). In view of these results, we decided to proceed without HAT upregulation and rely on the endogenous transaminase activity.

**Figure 8.**
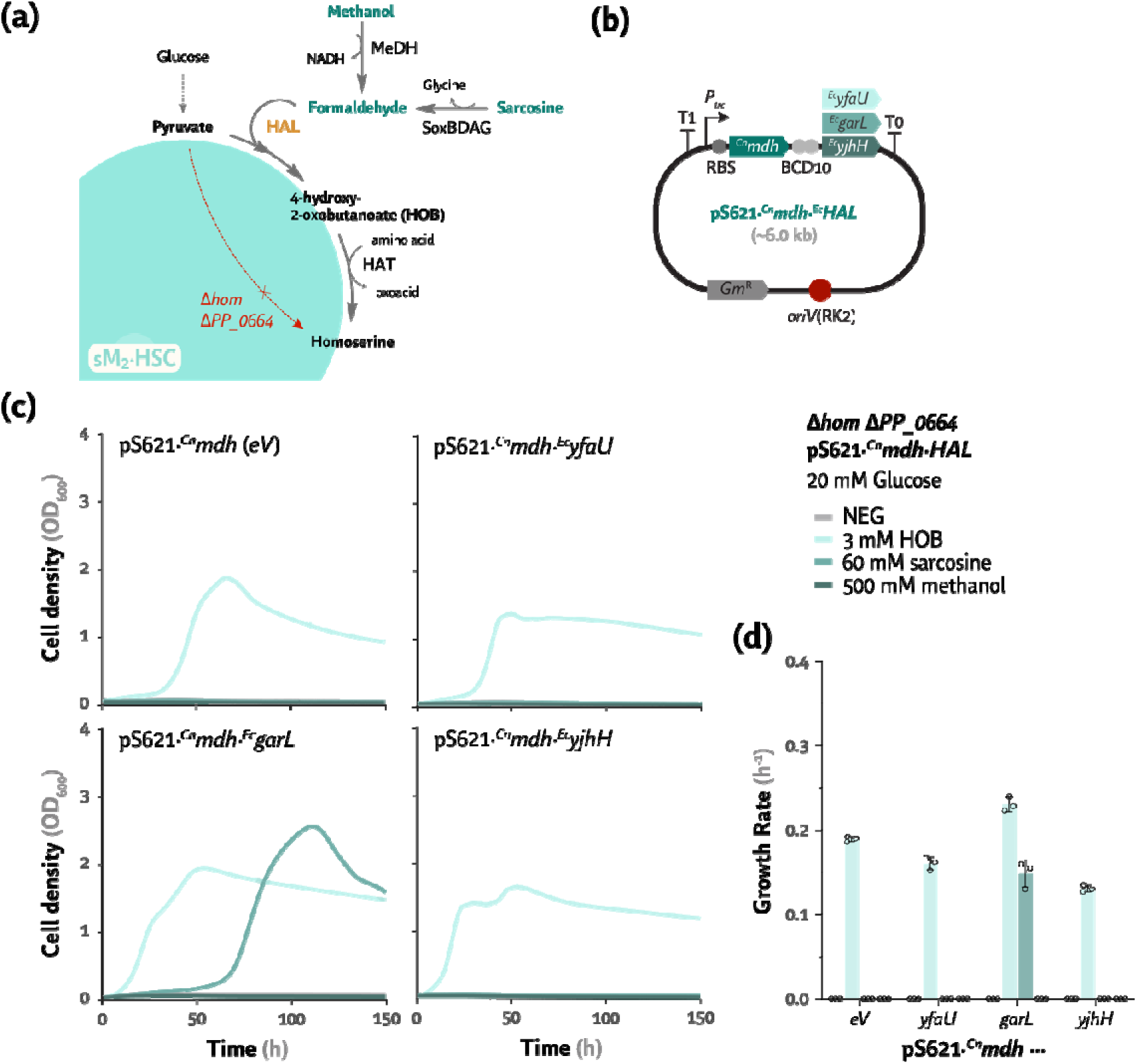
Screening of HOB aldolase activity. (a) Strain Δhom PP_0664 (lacking homoserine dehydrogenase activity) was employed as a homoserine auxotroph to identify candidates that perform the HOB aldolase (HAL) activity coupled to transamination. Formaldehyde can be produced from methanol via MeDHs or from sarcosine via the soxBDAG operon of P. putida. (b) Structure of the plasmids used to overexpress the CT4-1 MeDH from C. necator (^Cn^mdh) and three putative HOB aldolase genes (yfaU, garL, and yjhH from E. coli) as compared to an empty vector control. (c) Growth profiles of the selection strain Δhom PP_0664 in DBM medium with 20 mM glucose supplemented with 3 mM 4-hydroxy-2-oxobutanoate (HOB), 60 mM sarcosine, or 500 mM methanol. The tested strains harbor the pSEVA621 plasmids constitutively overexpressing ^Cn^mdh as well as the different HAT candidates or the empty vector (eV) control. (d) Growth rates corresponding to the growth profiles indicated in panel (c). Average values for bacterial growth (estimated as the optical density measured at 600 nm, OD_600_), and specific growth rate (μ, in h^−1^) ± standard deviation of three biological replicates are represented in all cases and individual data points are shown.

### GarL fromE. coli can act as a HOB aldolase inP. putida [M_2_·HSC]

The next step was selecting a suitable HAL by screening a library of putative HAL genes cloned in pSEVA621 and constitutively expressed alongside ^Cn^*mdh* (**Fig. S6**a, **Fig. 8**a–b). We tested aldolases that use pyruvate as a substrate from different heterotrophic bacteria, i.e., GarL, YfaU, YjhH, and DapA from *E. coli*; GllC, PanB, PP_1791, and Eda from *P. putida*; and HpaI from *P. aeruginosa*. *P*. *putida* Δhom PP_0664 containing the library of aldolase genes was grown on DBM medium with 20 mM glucose, supplemented with 3 mM HOB (positive control), 60 mM sarcosine (as a source of formaldehyde), or 500 mM methanol. Methylotrophic growth was only observed with ^Ec^*garL* overexpression (μ ∼ 0.15 h^−1^ and maximum OD_600_ ∼2.5) and partially for ^Pp^*panB* with 60 mM sarcosine (**Fig. S6**–b, **Fig. 8**c–d). No candidate showed growth in methanol (**Fig. 8**c). Since sarcosine, a source of intracellular formaldehyde, also yields glycine, we did not test the entirety of M_2_·HSC. Nevertheless, the homoserine-to-glycine submodule has already been shown to be active in M_2_·STC regardless of LtaE upregulation (**Fig. 7**a–c). Thus, combining the previous efforts could drive enough flux through the HOB aldolase from methanol, resulting in full activity of M_2_·HSC in *P. putida*.

### Heterologous expression of alanine-glyoxylate transaminase complements the glycine and serine auxotrophy [M_2_·mSC]

M_2_ of the mSC relies on carboxylation of pyruvate or PEP to oxalacetate, oxidation to malate, and activation to malyl-CoA by malate thiokinase (Mtk). Malyl-CoA is then broken down into acetyl-CoA and glyoxylate by malyl-CoA lyase (Mcl). Glyoxylate is finally transaminated into glycine by alanine-glyoxylate transaminases (Agt, **Fig. 1**c). Since Agt, Mtk, and Mcl are exogenous to *P. putida*, we first targeted the submodule for glyoxylate transamination and explored Agts from *H. sapiens*(^Hs^*agxt1*), *S. cerevisiae*(^Sc^*agx1*), and *P. denitrificans*(^Pd^*bhcA*), as well as *thiO* from *P. putida*(**Fig. 9**a). The four candidates were constitutively expressed in a pSEVA221 backbone in the glycine and serine auxotroph (*P*. *putida* Δ*serA*). The resulting strains were grown in DBM medium with 20 mM glucose, supplemented with 10 mM glycine or glyoxylate (**Fig. 9**b). Except for ^Pp^*thiO*, the three Agts complemented the glycine and serine auxotrophy. Among these, ^Sc^*agx1* performed best with μ ∼ 0.25 h^−1^ and a cell density of OD_600_ ∼ 2.5 with glyoxylate supplementation (**Fig. 9**b–c). We proceeded with this variant to implement the entire M_2_ of the mSC.

**Figure 9.**
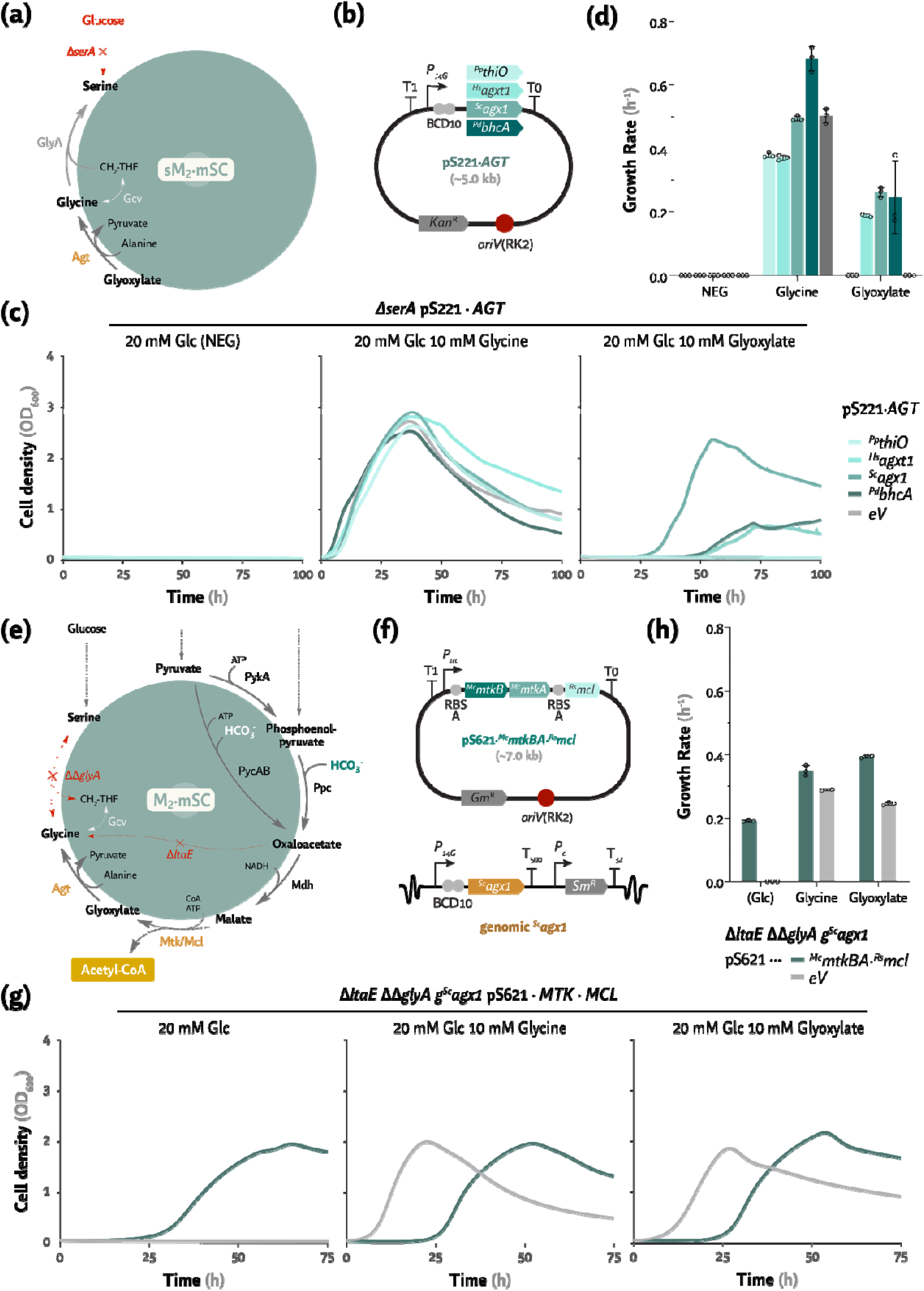
Heterologous transamination from glyoxylate to glycine, as well as expression of malate thiokinase and malyl-CoA lyase, rescues glycine and serine auxotrophy. (a) A ΔserA strain was employed as a serine-glycine auxotroph to identify AGT candidates that perform the alanine-glyoxylate transamination. (b) Structure of plasmids used to overexpress putative AGT genes (thiO from P. putida, agxt1 from H. sapiens, agx1 from S. cerevisiae, and bhcA from P. denitrificans). (c) Growth profiles of the ΔserA selection strain in DBM medium with 20 mM glucose supplemented with 10 mM glycine or 10 mM glyoxylate. The tested strains harbor pSEVA221 plasmids constitutively overexpressing AGT candidates or the empty vector (eV) control. (d) Growth rates corresponding to the growth profiles shown in panel (c). (e) Metabolic map of the auxotrophic strain ΔltaE glyA used as a glycine and C_1_ auxotroph to test malyl-CoA lyase (Mcl) and malate thiokinase (MtkAB) activities. (f) Structure of the plasmid used to overexpress mtkBA and mcl under control of the P_trc_ promoter and a strong RBS (RBS A, top), and genomic integration of the best AGT gene (^Sc^agx1) by Tn5 transposition (bottom). (g) Growth profiles of the ΔltaE glyA g^Sc^agx1 strain in DBM medium with 20 mM glucose supplemented with 10 mM glycine or 10 mM glyoxylate. A strain containing the empty vector (eV) is also plotted as a control. (h) Specific growth rates corresponding to the growth profiles shown in panel (g). Average values for bacterial growth (estimated as the optical density measured at 600 nm, OD_600_), and specific growth rate (μ, in h^−1^) ± standard deviation of three biological replicates are represented in all cases. Individual data points are also shown in panel (c).

### Glycine auxotrophy is rescued after adaptation and expression of malate thiokinase and malyl-CoA lyase as the complete module 2 of the modified serine cycle_2_·m[MSC]

Besides alanine-glyoxylate transamination, the mSC also relies on Mtk and Mcl (**Fig. 1**c). We tested the variants from *Methylococcus capsulatu* a*s* nd *Rhodobacter sphaeroide* in*s* a small expression library. The three genes (^Mc^*mtkB*, ^Mc^*mtkA*, and ^Rs^*mcl*) were constitutively expressed with different RBSs in an acetyl-CoA auxotrophic strain (Δ*aceEF*; **Fig. S7**a–b). *P. putida*Δ*aceEF* cannot synthetize acetyl-CoA or run the TCA cycle for biomass precursors and energy [68]. Expression of ^Mc^*mtkBA* and ^Rs^*mcl* complemented the Δ*aceEF* auxotrophy in DBM medium with 20 mM glucose, although with a low cell density (OD_600_ < 0.3, **Fig. S7**c). To reduce the carbon demand of this submodule and implement the full M_2_ of the mSC, we added the Agt activity, randomly integrating the ^Sc^*agx1* gene *via* Tn*5* transposition [69] into a glycine and C_1_ auxotroph (Δ*ltaE* ΔΔ*glyA*, **Fig. 9**e–f). After a few selective passages, the Δ*ltaE* ΔΔ*glyA g*^Sc^*agx1* strain grew in DBM medium with 20 mM glucose and 10 mM glyoxylate (**Fig. 9**a–d).

Next, the strongest RBSs for expression of *mtk-mc* c*l* loned in a pSEVA621 backbone was tested in the Δ*ltaE* ΔΔ*glyA g*^Sc^*agx1* strain for its ability to produce glyoxylate (**Fig. 9**e–f). When these combined activities were tested in DBM medium with 20 mM glucose, growth complementation was observed after a few selective passages (**Note S2**). Expectedly, supplementing the medium with glyoxylate or glycine restored growth of all strains (**Fig. 9**g–h). Thus, M_2_ of the mSC resulted in comparable growth parameters to the STC in DBM medium with 20 mM glucose, with μ ∼ 0.2 h^−1^ and maximum cell density (OD_600_) ∼ 2 (**Fig. 9**g–h). These results underscore the potential for implementing the full mSC in *P. putida*.

### Endogenous and heterologous activities lead to a fully active serine-threonine cycle in engineeredP. putida

After successfully implementing all module variants for the three synthetic SC, we concluded that the STC had the highest *in vivo* potential in *P. putida*. To build a consolidated and stable methylotrophic *P*. *putida* strain, we engineered all necessary activities expressed from the genome of *P*. *putida*ΔSGG-C_1_. In addition to the C_1_ assimilation module from methylotrophic bacteria, we overexpressed both the native *ltaE* to regenerate glycine and *yiaY* to upregulate endogenous PQQ-dependent MeDHs (**Fig. 10**a–b). We also deleted *thiO*, a FAD-dependent glycine/D-amino acid oxidase that produces glyoxylate [32] and could decrease flux through the STC competing with biomass formation. Finally, since we observed substantial biomass aggregation in media containing high methanol concentrations, we deleted the genes encoding the biofilm-producing adhesins LapA and LapF [70,71].

**Figure 10.**
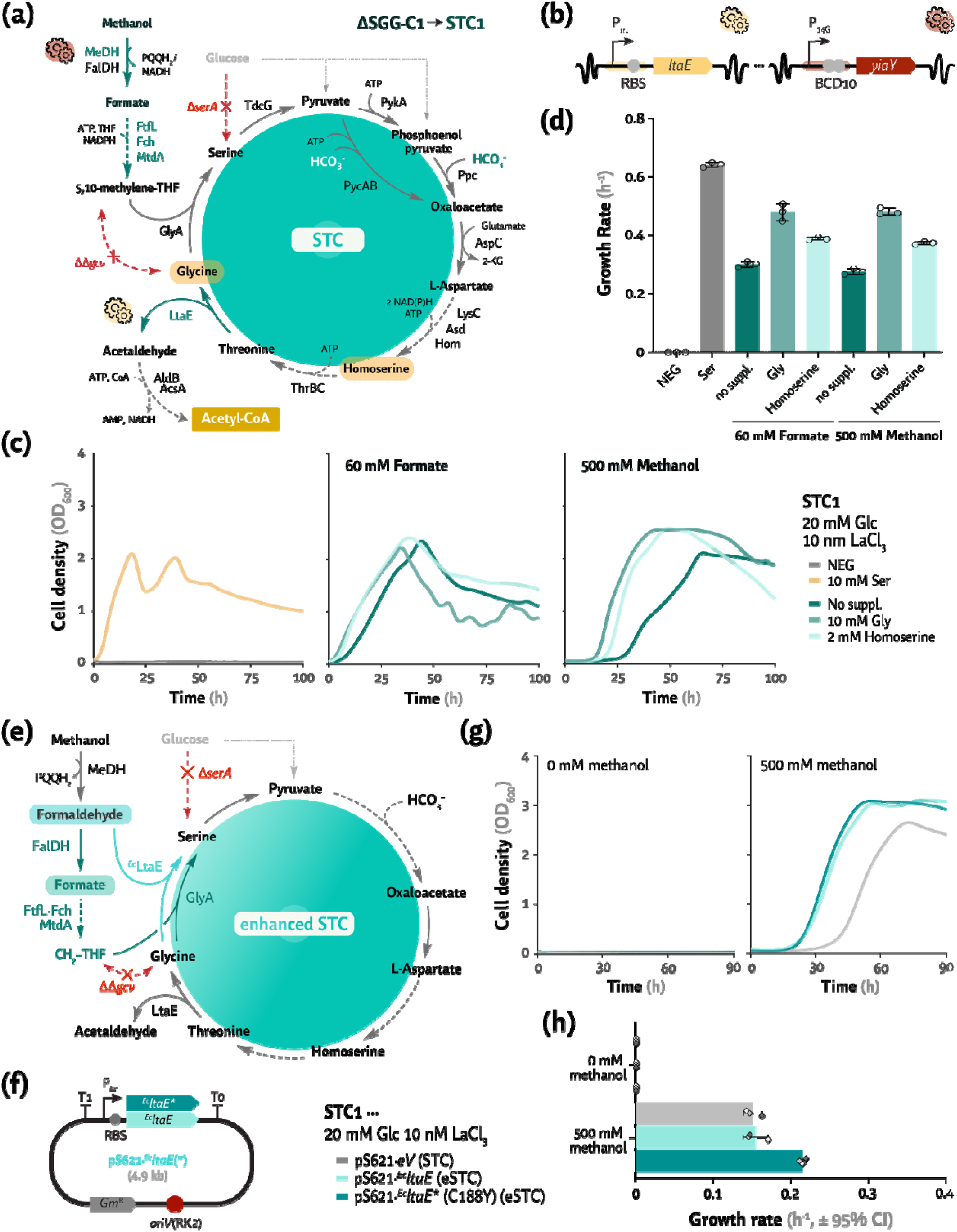
A combination of endogenous and heterologous activities activates the serine threonine cycle (STC) and a novel architecture for C_1_ assimilation, the enhanced STC. (a) A serine and glycine auxotrophic strain with further modifications (M_1_·STC, ΔSGG-C1) to overexpress the endogenous ltaE and yiaY. LtaE overproduction yields glycine from central metabolism whereas YiaY promotes methanol oxidation by derepressing the activity of endogenous PQQ-dependent alcohol dehydrogenases. This P. putida strain was termed STC1. (b) Genomic constitutive overexpression of the endogenous ltaE and yiaY via promoter exchange with the strong, constitutive P_trc_ and P_14G_ promoters, respectively. (c) Growth profile of strain STC1 in DBM medium with 20 mM glucose and 10 nM LaCl_3_ supplemented with 10 mM glycine, serine (positive control), or 2 mM homoserine. When needed, 60 mM formate or 500 mM methanol were also supplemented. (d) Specific growth rates corresponding to the growth profiles shown in panel (c). (e) Metabolic map for strain STC1 harboring the plasmids indicated in panel (f), encoding M_1_ of the HSC. Increasing the assimilation of C_1_ compounds directly via formaldehyde leads to a new synthetic metabolism, termed enhanced STC (eSTC). (f) Structure of plasmids used to overexpress ^Ec^ltaE and the new variant ^Ec^ltaE* (C188Y [72]) under the constitutive expression of P_trc_ and the canonical RBS. (g) Growth profiles of strain STC1 harboring pS621·^Ec^ltaE / ^Ec^ltaE* (C188Y) in DBM medium with 20 mM glucose, 10 nM LaCl_3_, and 500 mM methanol. The strain transformed with an empty vector (eV) is also plotted as a control. (h) Specific growth rates corresponding to the experiments shown in panel (g). Average values for bacterial growth (estimated as the optical density measured at 600 nm, OD_600_), and specific growth rate (μ, in h^−1^) ± standard deviation of three biological replicates are represented in all cases. Individual data points are shown. Error bars in panel (h) correspond to 95% confidence intervals (CI), with P-values < 0.05 for all pairwise comparisons where error bars do not overlap.

The resulting strain was termed *P*. *putida* STC1, and the detailed engineering steps are summarized in **Fig. S2**. We tested if *P*. *putida* STC1 can assimilate methanol into l-serine and regenerate glycine from pyruvate or oxaloacetate (i.e., M_1_ and M_2_ of STC, respectively, **Fig. 10**a). When strain STC1 was incubated in DBM medium with 20 mM glucose and 10 nM LaCl_3_, both formate and methanol were efficiently assimilated and promoted growth, with μ ∼ 0.3 h^−1^ and 0.28 h^−1^ and maximum OD_600_ ∼ 2.4 and 2.1, respectively (**Fig. 10**c–d). Thus, while the implementation of the STC was insufficient to support growth on methanol alone, the mixotrophic growth of *P*. *putida* STC1 when supplemented with C1 molecules indicates that the full STC is functionally active in this strain. STC was realized in *P. putida via* overproduction of the endogenous PQQ-MeDH and l-threonine aldolase as well as heterologous C_1_-assimilation *via* THF.

### An enhanced serine-threonine cycle, encompassing a serine aldolase activity, outperforms the parental pathway

Feedstock oxidation to CO_2_ cannot be completely abolished in an aerobic heterotroph, and we hypothesized that adding extra entry points for C_1_ moieties into central carbon metabolism could improve the ability of the engineered strain to utilize C_1_ substrates (**Fig. 10**e). Since the M_1_ variants from the STC or HSC are not mutually exclusive, including M_1_·HSC (i.e., serine aldolase activity) in the STC architecture could enable additional carbon assimilation at the formaldehyde level. To test this scenario, we transformed *P*. *putida* STC1, where all relevant STC genes are stably integrated in the chromosome, with plasmid pSEVA621·^Ec^*ltaE* (**Fig. 10**f). We also tested ^Ec^*ltaE*\*, a variant of the aldolase carrying the C188Y point mutation described to display faster kinetics [72].

When the fully engineered strain was incubated in DBM medium with 20 mM glucose, 10 nM LaCl_3_, and 500 mM methanol, all growth parameters improved compared to *P*. *putida* STC1. A decreased lag phase was observed, with faster growth (μ ∼ 0.21 h^−1^ for ^Ec^*ltaE*\*), and increased maximum cell density (OD_600_ ∼ 3, **Fig. 10**g–h). The wild-type ^Ec^*ltaE* supported a similar growth profile as ^Ec^*ltaE*\*, albeit the growth rate was equivalent to the STC1 strain (**Fig. 10**g–h). We named the new cycle *enhanced* serine-threonine cycle (eSTC). A calibration assay was performed to identify the concentration range at which methanol is assimilated (**Fig. S8**). We tested the STC1 strain harboring either ^Ec^*ltaE* or ^Ec^*ltaE\** in DBM medium with 20 mM glucose and 10 nM LaCl_3_ at different methanol concentrations (**Fig. S8**). No growth was observed for any of the variants when methanol was omitted, highlighting the need for a C_1_ substrate to complement the l-serine auxotrophy. Importantly, PedH, the native PQQ-MeDH, supported synthetic methylotrophy even at low (7.8 mM) methanol concentrations. Hence, both eSTC variants (with either ^Ec^*ltaE* or ^Ec^*ltaE\**) outperformed the parental STC in terms of maximum cell growth (OD_600_) and lag phase reduction, exhibiting more consistent growth profiles at lower methanol concentrations. Since specific growth rates and maximum cell densities were not substantially affected by the methanol concentration, further engineering efforts could be used to overcome other potential bottlenecks (e.g., glycine regeneration, **Fig. S8**b–c).

## DISCUSSION

The SC is a natural, oxygen-insensitive pathway that synthesizes acetyl-CoA from C_1_ molecules without carbon loss [16]. This is an attractive pathway to establish synthetic C_1_ assimilation, but engineering efforts thus far have been hindered by inherent bottlenecks, e.g., cycle efficiency, toxicity of intermediates, and interference with central metabolism in the host [16,47]. To address these challenges, we explored three synthetic SC variants, the STC, the HSC, and the mSC, systematically comparing their performance both *in silico* and *in vivo* through growth-coupled selection and modular engineering in a genome-reduced *P*. *putida* strain. *P*. *putida* has a highly flexible metabolism with a native architecture that aligns with implementing SCs. Indeed, the substantial fluxes through carboxylation reactions from pyruvate or PEP [73], producing oxaloacetate and assimilating CO_2_ (i.e., in M_2_ of the STC or mSC), favor the implementation of these synthetic C_1_ modules. Additionally, *P*. *putida* contains multiple copies of most endogenous enzymes involved in the cycles, including three homologs of serine deaminase (TdcG) and several C_1_ dehydrogenases [56].

Pathway modularization was essential for our engineering efforts. M_1_ covers carbon assimilation from glycine to serine, common to all synthetic cycles. This metabolic step is supported either by the THF-moiety from formate in the STC or mSC, or directly from formaldehyde *via* the serine aldolase reaction in the HSC. We engineered both variants in serine and glycine auxotrophs of *P*. *putida* using methanol and, as predicted by FBA (**Tables S6**–**S7**), both C_1_ entry points performed similarly well. However, the choice of MeDH significantly impacted the performance of the C_1_ assimilative module (**Table 1**). M_2_, on the other hand, involves the condensation of an additional C_1_ moiety (HCO_3_^−^ or formaldehyde) with pyruvate to produce oxaloacetate or homoserine. These intermediates are then cleaved by lyases to replenish the glycine pool and generate acetyl-CoA, which can be used for biomass formation and energy regeneration. Glycine is used to initiate a new cycle through M_1_, since the SCs are autocatalytic. To compare the three variants of this module, we utilized serine and/or glycine auxotrophies (**Table 1**). Our *in-silico* analysis indicated that M_2_ of the HSC should outperform the mSC variant, while the STC is the longest and most ATP- and NAD(P)H-demanding route. However, the STC leverages native pathways of *P*. *putida*, whereas the other cycle variants require extensive engineering [16,20,45–47]. Interestingly, *E. coli* GarL assimilated formaldehyde to complement the homoserine auxotrophy in the HSC (**Notes S3** and **S4**). The M_2_ variants of the STC and the mSC were predicted to operate with similar growth parameters, with the mSC being slightly more efficient. In a glycine auxotroph (ΔΔ*glyA*), the mSC variant showed better yields (cell density, OD_600_) compared to the STC (**Table 1**). However, the mSC requires extensive flux rewiring, since glycine is derived from the TCA cycle intermediates glyoxylate and malate. In contrast, M_2_ of the STC relies on existing pathways in *P*. *putida* and overexpressing the later enzyme in the module (LtaE) was sufficient to elicit module activity.

**Table 1.**
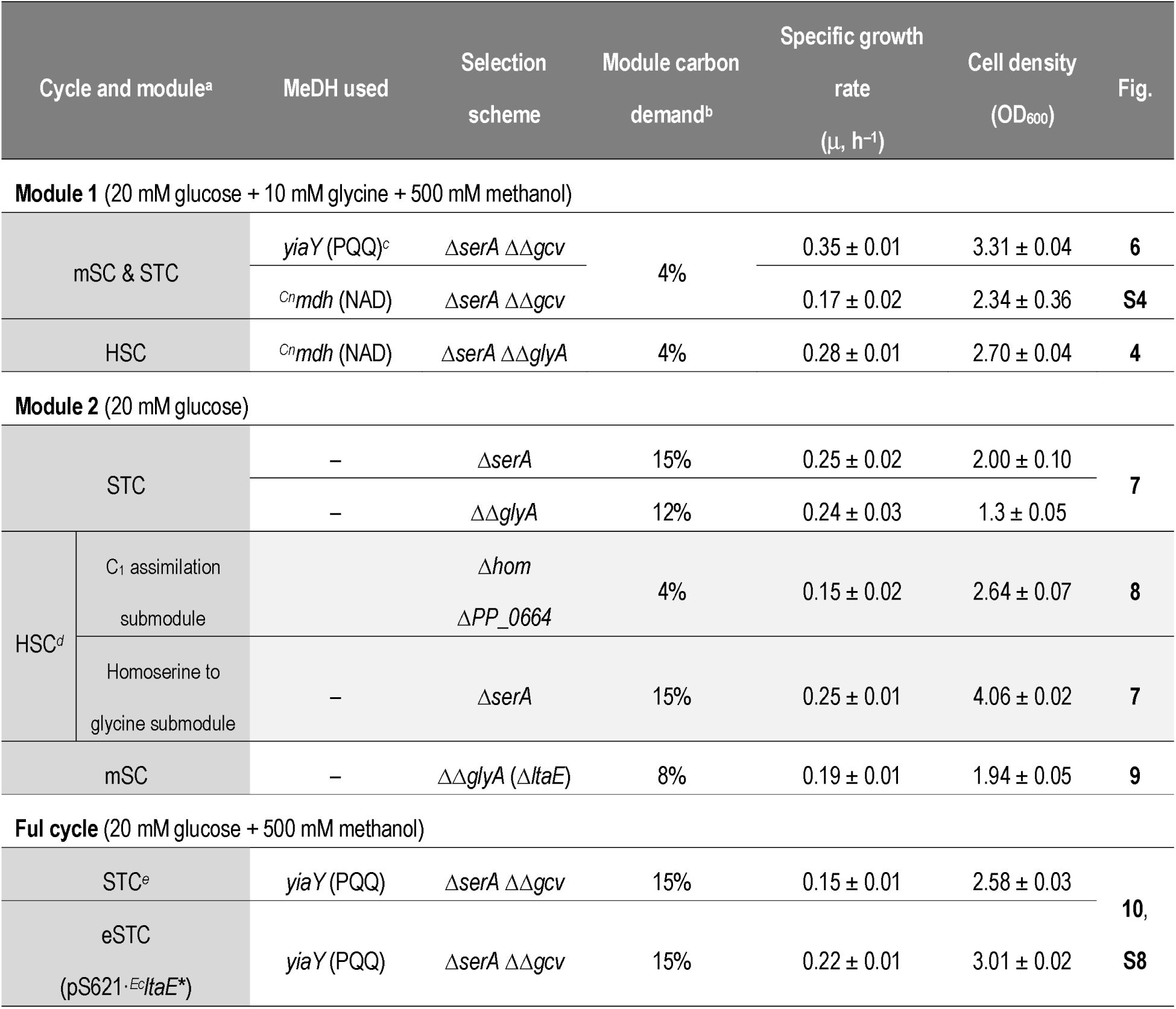
Module comparison of the three synthetic serine cycles tested in this work.

Another advantage of using *P. putida* as a host for synthetic methanol assimilation is its inherent alcohol oxidation capacities. Methanol oxidation is a major bottleneck in methylotrophic growth due to unfavorable thermodynamics, particularly since most synthetic methylotrophic engineering relies on NADH-dependent MeDH [16,20,45–47]. While NAD-MeDH may result in higher biomass yields, reaction thermodynamics are compromised, leading to a significant MDF decrease also observed during implementation of the HSC. Using PQQ-dependent MeDHs can overcome this bottleneck with increased thermodynamic favorability, resulting in higher growth rates [12,74,75]. We successfully engineered methanol oxidation using either a heterologous NAD-MeDH or the methanol activity of the native, broad-range PQQ-dependent alcohol dehydrogenases of *P. putida*(PedE and PedH). PQQ-MeDH activity was achieved by overexpressing *yiaY*, encoding its putative regulator [61]. Additionally, the presence of REEs (lanthanides) determined which dehydrogenase was expressed [33,62]. PedH activity, upregulated in the presence of lanthanides, yielded better growth parameters compared to PedE in the absence of LaCl_3_. No efforts to engineer full methylotrophy in synthetic hosts have used PQQ-dependent MeDHs. Only a recent study utilized PedE under mixotrophic conditions to implement a module of the reductive glycine pathway [76]. Unlike *E. coli*, which lacks the PQQ biosynthesis pathway, *P. putida* facilitates the engineering of methanol oxidation through PQQ-MeDH for both native and heterologous enzymes [77], including the *M. extorquen* A*s* M1 XoxFGJ reported in this work.

Building on the modularization efforts and endogenous methanol oxidation, we selected the STC as the highest potential for methylotrophy. Although energetically demanding, this cycle closely resembles the native metabolism, and it supported synthetic formatotrophy in engineered *E*. *coli* [47]. Hence, we constructed a streamlined, stable STC1 strain, incorporating the optimized STC modules into the genome of *P. putida*. This strain displayed efficient C_1_ assimilation (both formate and methanol) and fully replenished serine through the synthetic cycle. We showed that the STC relieved serine and glycine auxotrophies using its original module architecture. We envision that different module combinations, e.g., M_1_·HSC and M_2_·STC (or M_1_·HSC and M_2_·mSC), could also be implemented in *P. putida*.

Finally, we identified a new variant of the STC through module combination that outperforms the original STC. This new cycle, termed the *enhanced serine-threonine cycle*(eSTC), incorporates the serine aldolase activity from M_1_·HSC into the STC design. The eSTC not only mediated faster growth but also promoted higher cell densities. We infer that these improved growth parameters are due to increased C_1_ assimilation, as formaldehyde is also condensed directly by the eSTC. Therefore, increasing the scope of entry points for C_1_ substrates can enhance substrate availability, yields, and specific growth rates by reducing wasteful substrate oxidation. We also present the first direct comparison of engineered cycles in a heterotrophic organism. The results of this study, summarized in **Table 1**, offer interesting insights into the relative performance of these routes for C_1_ assimilation. Expanding the biochemical repertoire of *P*. *putida* to include C_1_ feedstocks paves the way for achieving a sustainable and truly circular carbon economy through synthetic assimilation. This approach will also guide future efforts in establishing synthetic metabolism in other heterotrophs of industrial interest.

## CONCLUDING REMARKS

Methanol, a cost-effective and renewable feedstock derived from CO₂ capture or renewable resources such as biomass, offers a promising alternative to petroleum-based substrates. In this sense, one-carbon feedstocks have strong potential to contribute to a circular carbon bioeconomy, enabling the production of bio-based plastics, green chemicals, and biofuels. When coupled with metabolically versatile and robust bacterial hosts, e.g. *P*. *putida*, these feedstocks hold promise for reducing the carbon footprint of industrial processes. Inspired by these features, in this work we demonstrated the use of three synthetic serine cycles integrated with the native activity of PQQ-dependent alcohol dehydrogenases, enabling the design of an enhanced cycle for methanol assimilation. Additionally, we expanded the synthetic biology toolkit by identifying and characterizing methanol-inducible promoters, which can be leveraged for engineering methylotrophy in non-canonical microbial hosts. These efforts constitute a substantial contribution to the metabolic engineering of *P*. *putida* for methanol utilization, introducing a novel serine cycle variant and contributing to a broader range of engineered microbes for sustainable biotechnology. The engineering strategies presented here pave the way for transforming methanol into valuable biochemicals, materials, and fuels through microbial biomanufacturing. This work addresses critical global challenges, including resource depletion, competition for food and feed, and climate change, by advancing the potential of methanol as a sustainable industrial feedstock.

## AUTHOR′S CONTRIBUTIONS

O.P. and P.I.N. conceived the project and designed the experiments. O.P. performed MDF. D.B. and O.P. performed and analyzed FBA. O.P., J.M.T, L.F.C., L.G. and H.S.M. performed the experiments leading to the results described in the article. All the authors analyzed the data and participated in the discussions included in this study. O.P. and P.I.N. wrote the manuscript, H.S.M. assisted in drafting the manuscript, with contributions from all the other authors.

## ACKNOWLEDGEMENTS

We thank Justine Turlin for fruitful discussions, Vittorio Rainaldi for providing the NADH-MeDH library, Lennart Schada von Borzyskowski for sharing ^Pd^bhcA, Hai He for feedback on implementation of M2·HSC, and Elad Noor for his help with MDF calculations. D.B. acknowledges funding from CSIRO through the CSIRO Postgraduate Top-Up Scholarship, supported by the Advanced Engineering Biology Future Science Platform. The financial support from The Novo Nordisk Foundation through TARGET (grant NNF21OC0067996) to P.I.N. is gratefully acknowledged.

## MATERIALS AND METHODS

### Bacterial strains, medium composition and culture conditions

All bacterial strains and plasmids are listed in **Table S8** and **Table S9**, respectively. *E*. *coli* DH5α λ*pir* [78] was used as cloning host, while the reduced-genome *P. putida* strain SEM11 [55] was selected for quantitative physiology and engineering purposes unless indicated otherwise. Lysogeny broth (LB) complex medium (containing 10 g L^−1^ tryptone, 5 g L^−1^ yeast extract, and 10 g L^−1^ NaCl) and de Bont minimal (DBM) medium were used for all cultivations [79]. DBM medium contained 3.88 g L^−1^ K_2_HPO_4_, 1.63 g L^−1^ NaH_2_PO_4_, 2 g L^−1^ (NH_4_)_2_SO_4_, and 0.1 g L^−1^ MgCl_2_·6H_2_O with the initial pH adjusted at 7.0 and supplemented with a trace elements solution [10 mg L^−1^ ethylenediaminetetraacetic acid (EDTA), 2 mg L^−1^ ZnSO_4_·7H_2_O, 1 mg L^−1^ CaCl_2_·2H_2_O, 5 mg L^−1^ FeSO_4_·7H_2_O, 0.2 mg L^−1^ Na_2_MoO_4_·2H_2_O, 0.2 mg L^−1^ CuSO_4_·5H_2_O, 0.4 mg L^−1^ CoCl_2_·6H_2_O, and 1 mg L^−1^ MnCl_2_·2H_2_O] [80]. When needed, kanamycin (Km) and gentamicin (Gm) were supplied at 50 μg mL^−1^ and 10 μg mL^−1^, respectively.

Overnight cultures in LB medium were diluted 1/100 to inoculate a 5-mL preculture of DBM medium with 20 mM glucose in a 50-mL culture tube and incubated at 30°C and 250 rpm for ca. 18 h. This overnight culture was washed with DBM medium without any carbon source prior to the inoculation of the main culture in 96-well microtiter plates with the appropriate carbon source(s) as described in the text. All growth assays were performed in the presence of antibiotics, unless the strain did not harbor any plasmid. For cultivations in 96-well microtiter plates, 150 μL of a cell suspension at an OD_600_ of 0.05 were incubated in an Epoch2 microtiter plate reader (BioTek Instruments Inc.; Winooski, VT, USA) with 50 μL of mineral oil to prevent evaporation. Measurements obtained with microtiter plate readers were calibrated against a tabletop spectrophotometer. The specific growth rate (μ) and, when relevant, the extension of the lag phase (λ) were calculated using *QurvE* (www.qurveanalysis.com) by performing a smooth spline fit on the growth data [81].

### Construction of (deletion) plasmids

The suicide plasmids and overexpression plasmids noted in **Table S9** were constructed using USER cloning [82–85]. For deletion plasmids, DNA fragments, consisting of ca. 500-bp upstream and downstream regions around the locus to be eliminated, were amplified with Phusion *U* Hot Start^TM^ DNA polymerase (ThermoFisher Scientific Co.) using uracil-containing primers. The pGNW2 backbone [86] was digested with *Dpn* I prior to mixing 1 μL of *Dpn* I-treated vector with 100 ng of each PCR fragment and 1 μL of USER^TM^ enzyme (New England BioLabs) in a final volume of 10 μL. The reaction was incubated for 30 min at 37°C, followed by a temperature decrease over 3 min (from 28°C to 20°C, 1°C per step) and a final incubation step at 10°C for at least 10 min. Finally, chemically-competent *E*. *coli* DH5α λ*pir* cells were transformed *via* heat shock with 5 μL of the USER mix; upon recovery, the cell suspension was plated onto selective LB medium agar plates containing the corresponding antibiotic.

### Construction of mutant ***P. p****utida* **strains**

The corresponding suicide pGNW2-derivative plasmid was delivered into the cells by triparental conjugation with the corresponding DH5α λ*pir* harboring the specific suicide plasmid, the *P. putida* strain of interest, and the *E. coli* helper strain *E. coli* HB101 carrying plasmid pRK2013 [87]. The three strains were incubated in LB plates for over 5 h at 30°C and subsequently plated in LB plates supplemented with the antibiotic of interest and Irgasan. Positive co-integration events were further transformed with pQURE6·H (**Table S9**), a conditionally-replicative plasmid bearing the meganuclease gene *I-SceI* [88]. I-SceI cuts pGNW2 co-integrants within the chromosome, thus forcing a second homologous recombination event. This was performed by electroporating 50 ng of plasmid DNA into 50 μL of freshly-prepared electrocompetent *P*. *putida* cells, previously washed three times with 300 mM sucrose. Electroporation was performed with a Gene Pulser XCell (Bio-Rad) set to 2.5 kV, 25 μF capacitance and 200 Ω resistance in a 2-mm gap cuvette. Cells were recovered in 1 mL of LB medium supplemented with 2 mM of 3-methylbenzoate (3-*m* Bz) for at least 3 h at 30°C and plated onto LB medium agar containing the corresponding antibiotic(s) and 1 mM 3-*m* Bz to induce both plasmid replication and I-*Sce* I expression. Positive clones were identified by colony PCR, verified by DNA sequencing, and cured from the resolving plasmid by serial dilution under non-selective conditions. All sequences used were native with the exception of ^Hs^*agxt1* and ^Sc^*agx1* which were codon-optimized.

### Maximum-minimum driving force (MDF) analysis

MDF analysis [89] was applied to evaluate and compare the different natural and modified SC in the context of *P. putida* KT2440 using eQuilibrator-API [90]. We also compared these metabolisms to the reductive glycine pathway (rGlyP) and the ribulose monophosphate (RuMP) pathway [17,32,91]. Metabolite concentrations were constrained to the range of 1-10 mM [89,92] with a few exceptions: (i) the upper bound for formaldehyde was constrained to 1 mM, the highest concentration tolerated by *P. putida* KT2440 [93]; (ii) l-glutamate and 2-ketoglutarate were set as amino-donors at 100 mM and 0.5 mM, respectively [20,94]; (iii) the intracellular pH was set to 7.8 [95]; (iv) the ionic strength and –log_10_[Mg^2+^] (pMg) were assumed to be 0.25 M and 3, respectively [20,92,96]; (v) since the MHPT oxidation of formaldehyde is not supported by *e*Quilibrator [20], we adopted the glutathione-independent oxidation reaction for the SCs instead (except for the HSC), (vi) all carboxylations were assumed to use CO_2_ as a substrate to simplify the calculations, given that such reactions are pH-independent, unlike those involving bicarbonate [92]. The low (ambient CO_2_) and high CO_2_ concentrations in solution were set to 10 μM and 1 mM, respectively.

High CO_2_ concentrations are needed, for instance, by the rGlyP [97]. All pathways used methanol as a sole carbon source and acetyl-CoA or pyruvate as a substrate. Given that the number of carbons differs in each substrate, the MDF per C-mol is reported in kJ C-mol^−1^ [91]. The scripts and further details on these calculations can be found at https://github.com/puiggene07/PubSuppl within the 2024_Ser_Cycles_P_putida directory.

### Flux Balance Analysis (FBA)

For *in silico* comparison of the different SC modules, FBA was performed with the COBRApy python package [98,99]. Simulations were run with a curated version of the latest genome-scale metabolic model available for strain KT2440 [51,52]. The model was further refined based on the following considerations: (i) the reversibility of GlyA (GHMT2) in glycine metabolism; (ii) our experimental evidence demonstrating that homoserine dehydrogenase (HSDy) operates irreversibly in *P*. *putida* during threonine regeneration, as observed in E. coli [20]; and (iii) the lack of natural formatotrophy *via* PurU in purine/pyrimidine biosynthesis in *P*. *putida*[32], leading to GARFT being set as irreversible. In addition, autotrophy based on CO_2_ assimilation *via* the lipoamide-dependent complexes (including AKGDa and PDHa) was prevented by setting the reactions as irreversible. The formaldehyde dismutase (FALDM) reaction was deactivated as this activity was never isolated in *P. putida*. Finally, genes borne by the TOL plasmid (reactions with pWW0 gene IDs) were eliminated and nickel (Ni^2+^) was removed from the biomass function, as *P. putida* does not have that requirement for growth [100]. For the C_1_ assimilation modules, the respective reactions were introduced in the model.

The *in-silico* comparison of the different synthetic modules was based on their capability of assimilating C_1_ substrates, their predicted maximal biomass formation rate, as well as the amount of directly oxidized C_1_ compounds (equal to the rate of the native formate dehydrogenase reaction), and the total CO_2_ emission rate. CO_2_ formation is a primary result of NAD(P)H generation; hence, it can be adopted as an indirect indicator for the energy demand of the respective pathway. Conditions were tested for co-utilization of glucose and methanol, full methylotrophy, and growth solely in glucose. Carbon equimolar uptake rates were set for the respective primary substrates (30 mmol g^−1^ h^−1^). For co-utilization of glucose and methanol, methanol uptake rates were additionally set to 5 mmol g^−1^ h^−1^ (high methanol uptake) or 2 mmol g^−1^ h^−1^ (low methanol uptake). Reactions were disabled according to the different *in vivo* selection schemes *via* the COBRApy function mode.genes.id.knockout().

### Chemicals and reagents

Chemicals were purchased from Sigma-Aldrich Co. (St. Louis, MO, USA) unless otherwise indicated, and oligonucleotides were synthesized by Integrated DNA Technologies Inc. (Coralville, IA, USA). DNA sequencing was performed at Eurofins Genomics (Ebersberg, Germany). All primers used in this study are listed in **Table S10**. PCR reactions were performed using Phusion *U* Hot Start^TM^ DNA polymerase, purchased from ThermoFisher Scientific Co. (Waltham, MA, USA). The commercial One*Taq*^TM^ master mix from New England BioLabs (Ipswich, MA, USA) was used for colony PCRs. 4-Hydroxy-2-oxobutanoate (HOB, in its lactone form) was purchased from Synthenova (France).

### Statistical analysis

Data analysis was performed using Prism 9.0.2 (GraphPad Software Inc.; San Diego, CA, USA). All reported values are indicated as averages ± standard deviation of at least three independent biological replicates as specified in the legend of the corresponding figures. Considering that most negative controls (i.e., strains transformed with empty vectors) did not show any growth, enzyme variants that restored methylotrophic growth were considered to be statistically significant [101].

## LIST OF ABBREVIATIONS

AGT: Alanine-glyoxylate transaminase
ALE: Adaptive laboratory evolution
C1: One-carbon (substrate/moiety)
DBM: medium de Bont minimal medium
EDEMP: cycle Entner-Doudoroff–Embden-Meyerhof-Parnas–pentose phosphate cycle
eSTC: Enhanced serine threonine cycle
FBA: Flux balance analysis
HOB: 4-Hydroxy-2-oxobutanoate
HAL: HOB aldolase
HAT: HOB transaminase
MDF: Maximum-minimum driving force
MeDH: Methanol dehydrogenase
mSC: Modified serine cycle
M1: Module 1
M2: Module 2
MTK: Malate thiokinase
MCL: Malyl-coenzyme A lyase
NAD(P)+/H: Nicotinamide adenine dinucleotide (phosphate) (oxidized/reduced)
NAD-MeDH: NAD(P)H-dependent methanol dehydrogenases
OD600: Maximal bacterial cell density
PDH: Pyruvate dehydrogenase
PQQ: Pyrroloquinoline quinone
PQQ-MeDH: PQQ-dependent methanol dehydrogenase
REEs: Rare earth elements
rGlyP: Reductive glycine pathway
RuMP pathway: Ribulose monophosphate pathway
SAL: Serine aldolase
SC: Serine cycle
sM1/2: Sub-module of Module 1/2
STC: Serine threonine cycle
TCA cycle: Tricarboxylic acid cycle
THF: Tetrahydrofolate
μ: Specific growth rate (h–1)

