## Supplemental Data for "Systematic engineering of synthetic serine cycles in Pseudomonas putida uncovers emergent topologies for methanol assimilation"

by

Òscar Puiggené[a], Jaime Muñoz-Triviño[a], Laura Civil-Ferrer[a], Line Gille[a], Helena Schulz-Mirbach[b], Daniel Bergen[c,d], Tobias J. Erb[b,e], Birgitta E. Ebert[c], and Pablo I. Nikel[a*]

[a] The Novo Nordisk Foundation Center for Biosustainability, Technical University of Denmark, Kongens Lyngby, Denmark

[b] Department of Biochemistry and Synthetic Metabolism, Max Planck Institute for Terrestrial Microbiology, Marburg, Germany

[c] Australian Institute for Bioengineering and Nanotechnology (AIBN), The University of Queensland, Brisbane, Queensland, Australia

[d] Advanced Engineering Biology Future Science Platform, CSIRO Ecoscience Precinct, Dutton Park, QLD, Australia

[e] Center for Synthetic Microbiology (SYNMIKRO), Marburg, Germany

**Running title:** Systematic implementation of methylotrophic cycles

**Keywords:** *Pseudomonas putida*; Growth-coupled selection; Metabolic engineering; Methylotrophy; Methanol; Serine cycle

* Correspondence to:

**Pablo I. Nikel**

The Novo Nordisk Foundation Center for Biosustainability,

Technical University of Denmark

2800 Kongens Lyngby

Denmark

Tel: (+45) 93 5119 18

### Supplemental Material

#### *Supplemental Figures*

Figure S1. Optimized thermodynamic profiles and maximum-minimum driving force (MDF) analysis. S3
Figure S2. Chart of all growth-coupled selection strains constructed and tested in this study. S6
Figure S3. Testing a library of low-specificity L-threonine aldolases in strain ΔserA ΔΔglyA ΔfrmAC. S7
Figure S4. Testing M1 of the serine-threonine and modified serine cycles via tetrahydrofolate. S8
Figure S5. Screening of transaminase activity on 4-hydroxy-2-oxobutanoate to yield homoserine. S9
Figure S6. Testing a library of putative HOB aldolases in a homoserine auxotroph. S10
Figure S7. Complementation of acetyl-CoA auxotrophy via malate thiokinase and malyl-CoA lyase. S12
Figure S8. The enhanced serine-threonine cycle (eSTC), using the native PQQ-dependent methanol dehydrogenases, enables methanol assimilation at low substrate concentrations. S13

#### *Supplemental Tables*

Table S1. Enzyme name, function, and reaction for each (modified) serine cycle tested in this study. S14
Table S2. Natural and modified serine cycles and their maximum-minimum driving force (MDF) compared to assimilative pathways producing $C_3$ intermediates S16
Table S3. Specific growth rate, C1 oxidation, and metabolic turnover ($CO_2$ emission) for the native serine cycle (SC) and its variants engineered in *P. putida* predicted by flux balance analysis. S17
Table S4. Specific growth rate, C1 oxidation, and metabolic turnover ($CO_2$ emission) for the native serine cycle (SC) and its variants engineered, as individual modules, in *P. putida* predicted by flux balance analysis at high methanol uptake rate. S18
Table S5. Specific growth rate, C1 oxidation, and metabolic turnover ($CO_2$ emission) for the native serine cycle (SC) and its variants engineered, as individual modules, in *P. putida* predicted by flux balance analysis in the presence of glucose. S19
Table S6. Specific growth rate, C1 oxidation, and metabolic turnover ($CO_2$ emission) for the native serine cycle (SC) and its variants engineered, as individual modules, in *P. putida* predicted by flux balance analysis in the presence of glucose and methanol (high uptake rate). S20
Table S7. Specific growth rate, C1 oxidation, and metabolic turnover ($CO_2$ emission) for the native serine cycle (SC) and its variants engineered, as individual modules, in *P. putida* predicted by flux balance analysis in the presence of glucose and methanol (low uptake rate). S21
Table S8. Bacterial strains used in this study. S22
Table S9. Plasmids used in this study S23
Table S10. Oligonucleotides used in this study. S27

#### *Supplemental Notes* S30

#### *Supplemental References* S31

**Fig. S1. Optimized thermodynamic profiles and maximum-minimum driving force (MDF) analysis.**

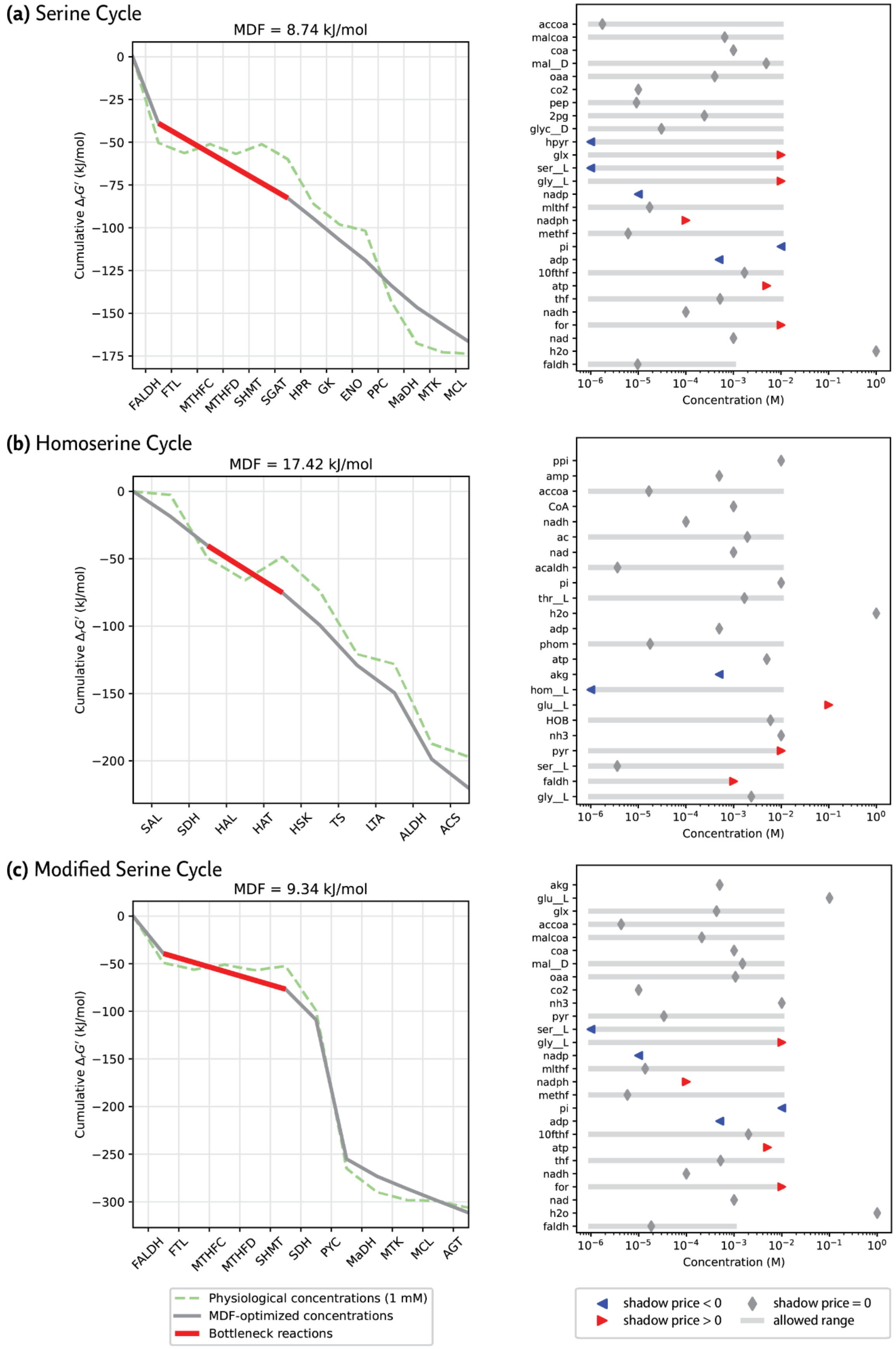

**(d)** Serine-Threonine Cycle

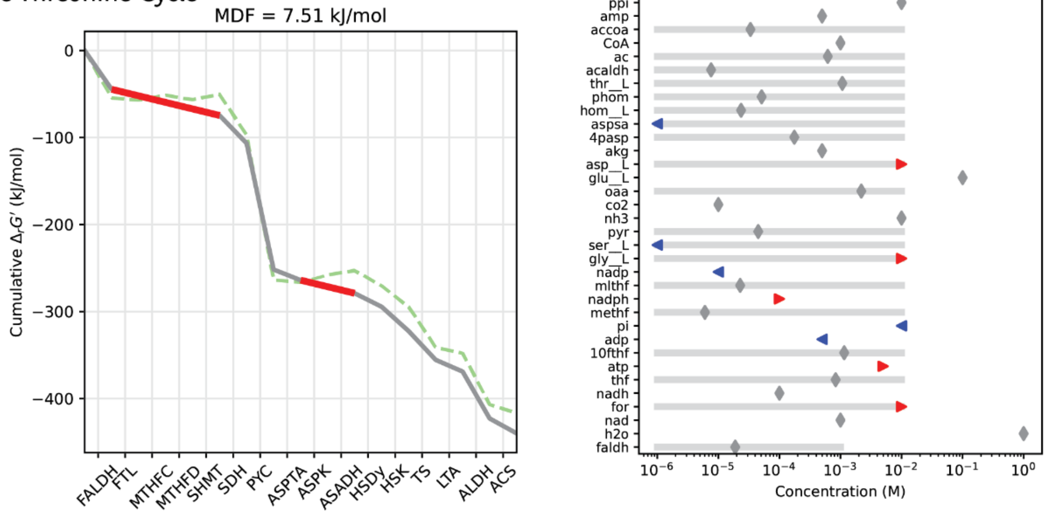


**(e)** Reductive Glycine Pathway

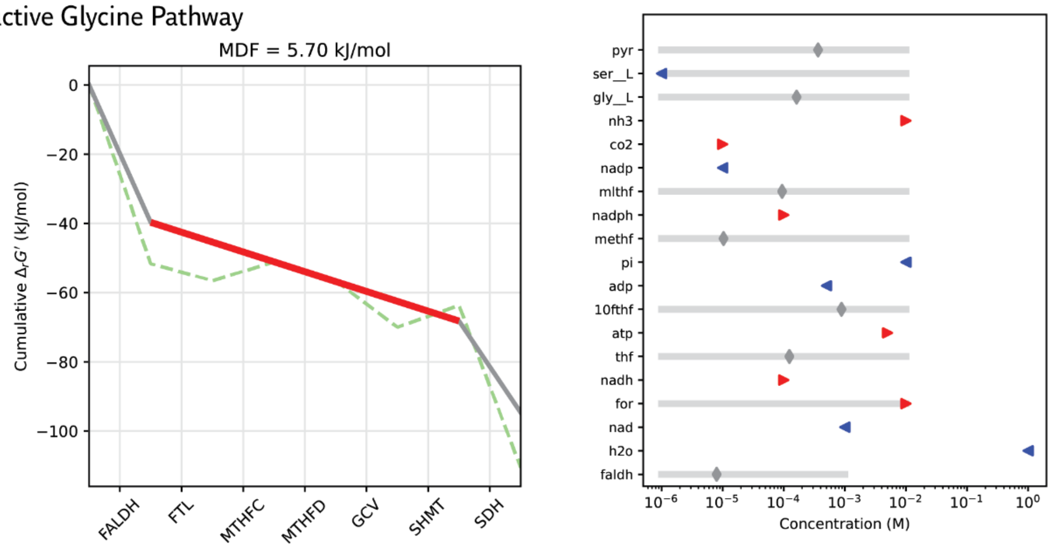


**(f)** Ribulose Monophosphate Pathway

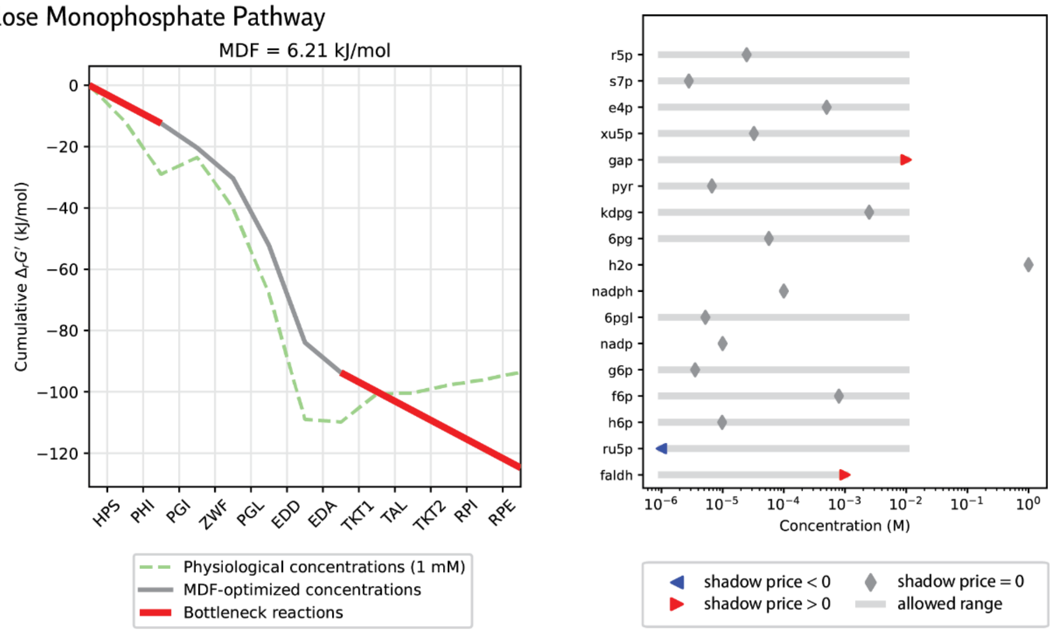

In all cases, the left panels display the energetic profile of the $C_1$ assimilation pathways, including the natural and modified serine cycles (a–d), in comparison to the reductive glycine pathway (e) and RuMP pathway (f). The right panels show metabolite concentrations within the physiological range (see *Materials and Methods*) after MDF optimization. Formaldehyde was used as the substrate for these simulations. Green dotted lines represent the $\Delta rG'_0$ values of pathway reactions at pH = 7.8. Grey lines (and red lines) indicate the $\Delta_r G'$ values of pathway reactions after MDF optimization, with red lines highlighting the predicted bottleneck reactions. Cofactors, such as NADH or ATP, were fixed at specified concentrations (see *Materials and Methods*). The shadow price is indicated in blue (shadow price < 0), grey (shadow price = 0), or red (shadow price > 0).

**Fig. S2. Chart of all growth-coupled selection strains constructed and tested in this study.**

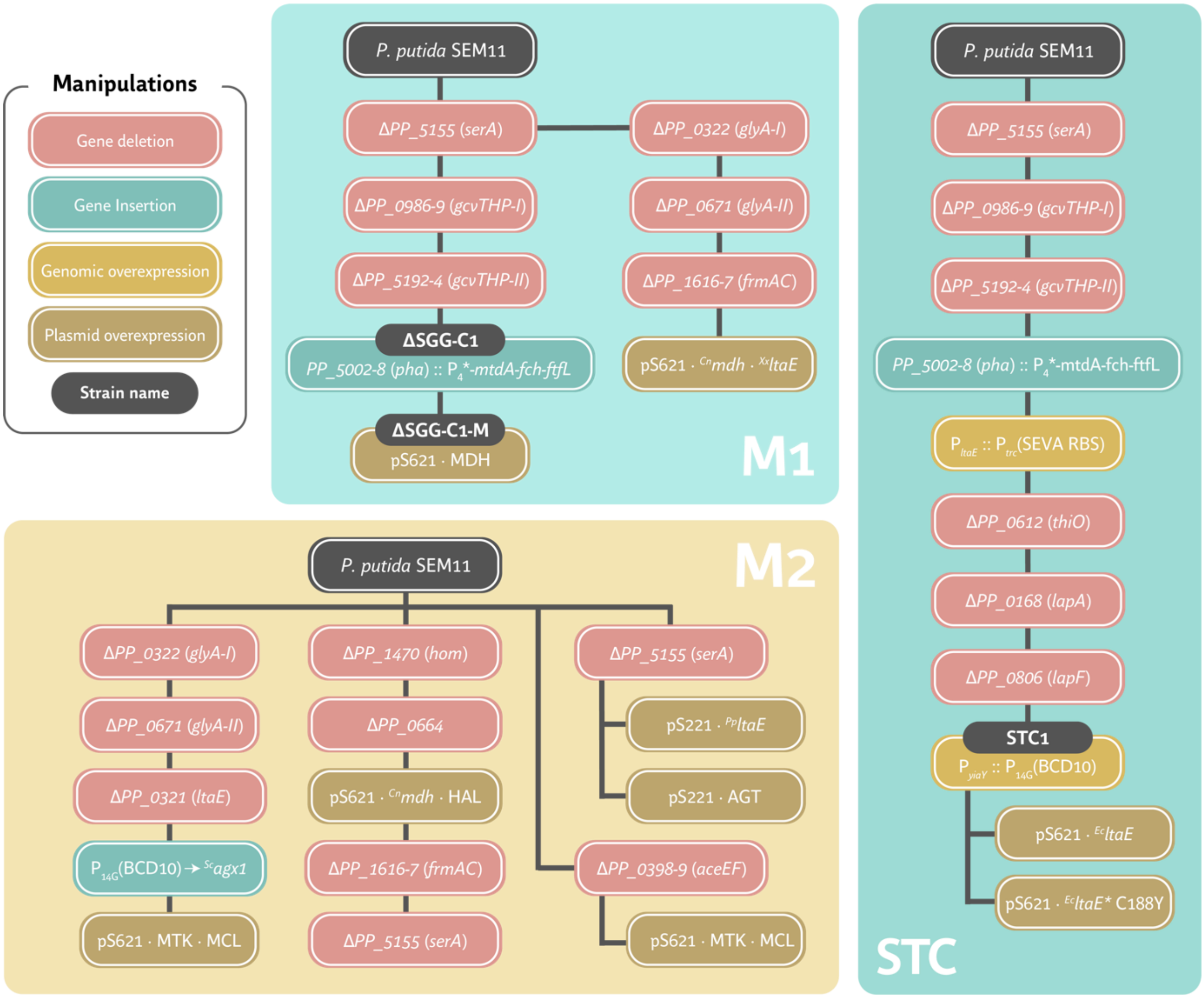


For details on the constructs and naming of strains and plasmids, please refer to **Tables S8-S9**.

**Fig. S3. Testing a library of low-specificity L-threonine aldolases in strain ΔserA ΔΔglyA ΔfrmAC.**

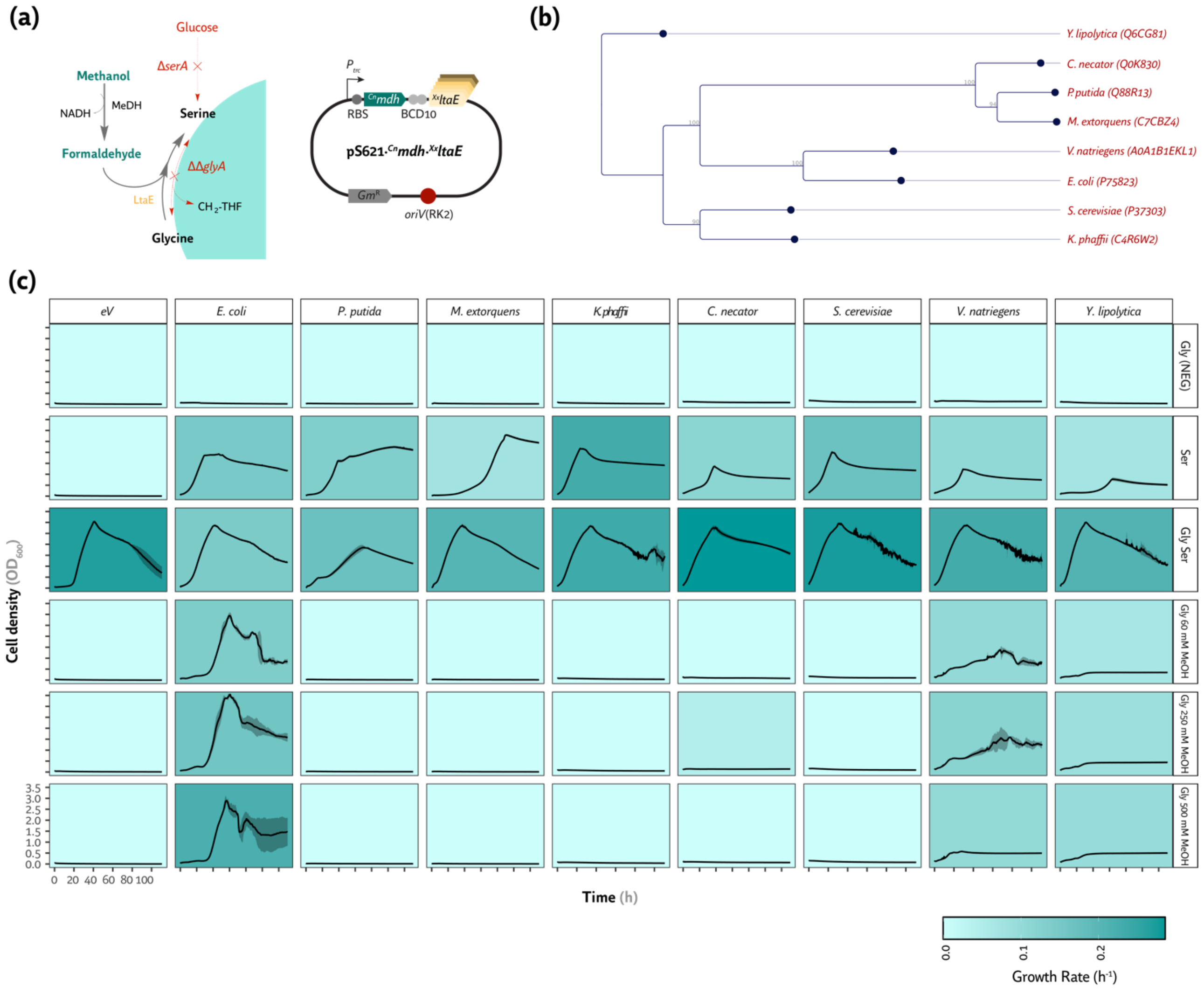


(a) Metabolic map of the auxotrophic strain Δ*serA* ΔΔ*glyA* Δ*frmAC*, used to test the side reaction of LtaE for assimilating formaldehyde into glycine to produce serine. A plasmid map is also provided; a canonical SEVA RBS was used (5'-AGG AGG AAA AAC AT-3' (Silva-Rocha et al., 2013)) for overexpression. The gene encoding the CT4-1 methanol dehydrogenase was overexpressed alongside the different LtaE variants. (b) Phylogram showing the phylogenetic relationships among LtaE variants (Uniprot code provided for each variant). (c) Growth profiles of strain Δ*serA* ΔΔ*glyA* Δ*frmAC* carrying these variant library plasmids in de Bont minimal medium supplemented with 20 mM glucose, 10 mM serine, 10 mM glycine, and [60, 250, or 500] mM methanol as indicated. Specific growth rates were also calculated. Average values for cell density (optical density at 600 nm, $OD_{600}$) and specific growth rates ($\mu$, in $h^{-1}$) ± standard deviation of three biological replicates are shown in all cases.

**Fig. S4. Testing M1 of the serine-threonine and modified serine cycles *via* tetrahydrofolate.**

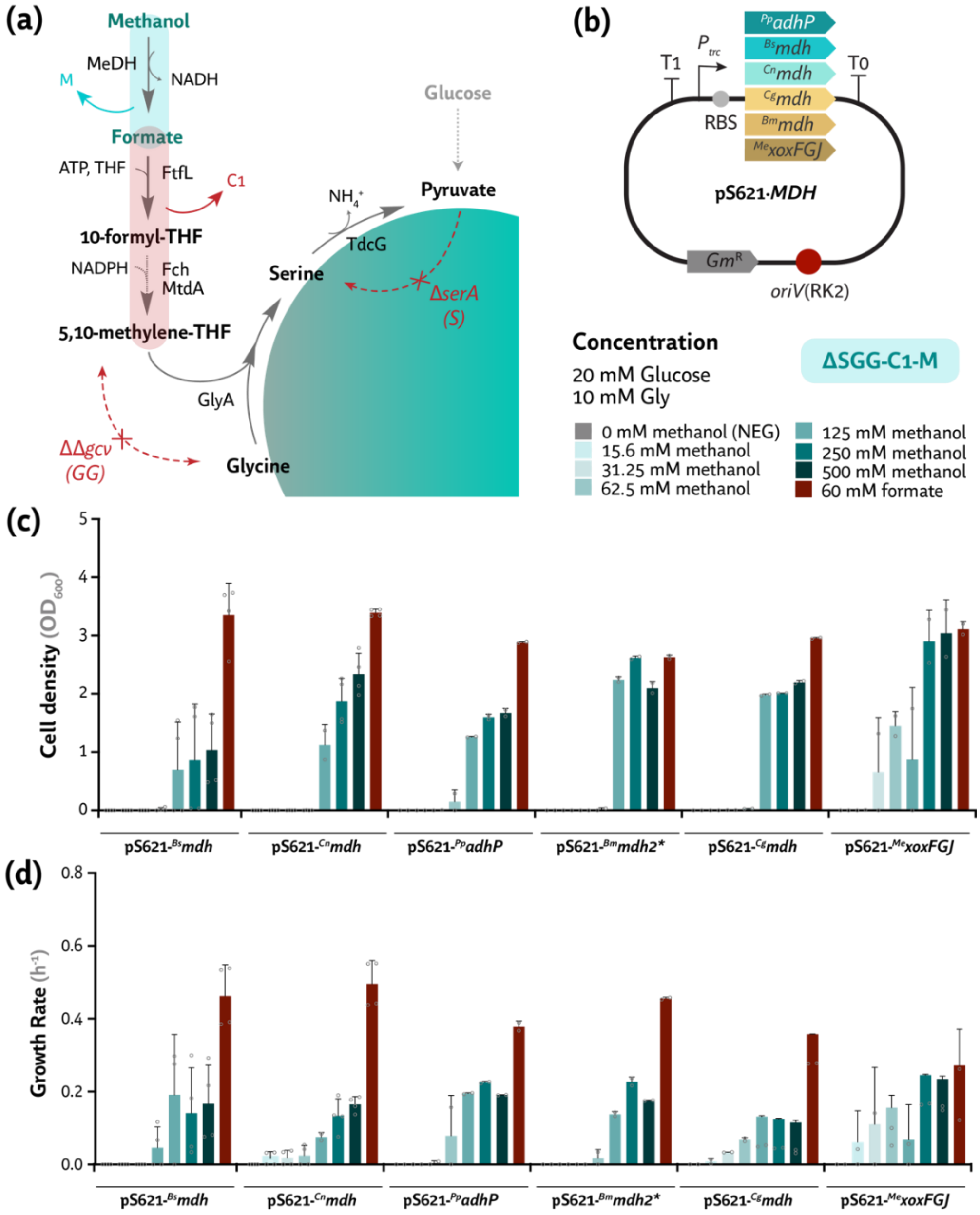


(a) Metabolic map of the auxotrophic strain Δ*serA* ΔΔ*gcvTHP* (referred to as Δ*SGG*) used to test one-carbon assimilation *via* FtfL, Fch, and MtdA (indicated as $C_1$) and methanol. Methanol oxidation was facilitated by overexpressing several endogenous putative methanol dehydrogenases (MDH, denoted as M). (b) Plasmid maps of the methanol dehydrogenase genes (MDH) overexpressed in strain ΔSGG-$C_1$ (yielding strain ΔSGG-$C_1$-M). (c) Maximum $OD_{600}$ and specific growth rates of the different MDH variants overexpressed in strain ΔSGG-$C_1$-M. Strains were grown in de Bont minimal medium with 20 mM glucose and 10 mM glycine, along with varying concentrations of methanol (or 60 mM formate, as a positive control). Average values for cell density (optical density measured at 600 nm, $OD_{600}$) and specific growth rates (μ, in $h^{-1}$) ± standard deviation of three biological replicates are represented in all cases. Individual data points are shown whenever relevant.

**Fig. S5. Screening of transaminase activity on 4-hydroxy-2-oxobutanoate to yield homoserine.**

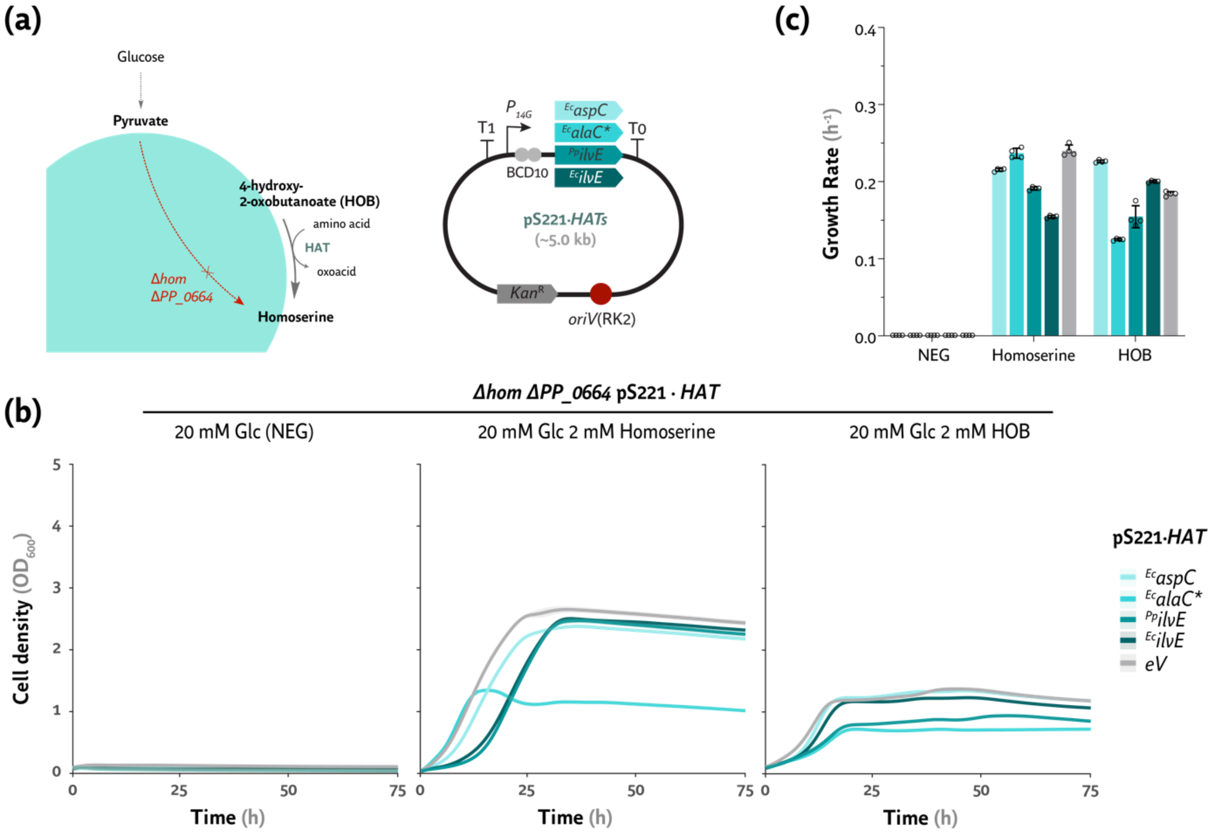


(a) *Left*: Metabolic map of the auxotrophic strain Δ*hom* Δ*PP_0664* (lacking homoserine dehydrogenase activity) used as a homoserine auxotroph to identify native or heterologous 4-hydroxy-2-oxobutanoate (HOB) transaminases (HAT). *Right*: Plasmid maps used to overexpress putative HOB transaminase genes (*aspC*, *alaC** carrying the A142P Y275D mutations, and *ilvE* from *E. coli*, as well as *ilvE* from *P. putida* KT2440 (Bouzon et al., 2017; He et al., 2020)) compared to an empty vector control (eV). (b) Growth profiles of the Δ*hom* Δ*PP_0664* selection strain in de Bont minimal medium with 20 mM glucose and supplemented with 2 mM homoserine or 2 mM 4-hydroxy-2-oxobutanoate (lactone form). The strains carried the corresponding pSEVA221 plasmid constitutively overexpressing the different HAT candidates or the empty vector (eV). (c) Specific growth rates corresponding to the growth profiles of panel (b). Average values for the cell density (optical density measured at 600 nm, $OD_{600}$) and specific growth rate ($\mu$, in $h^{-1}$) ± standard deviation of three biological replicates are represented in all cases, with individual data points shown when relevant.

**Fig. S6. Testing a library of putative HOB aldolases in a homoserine auxotroph.**

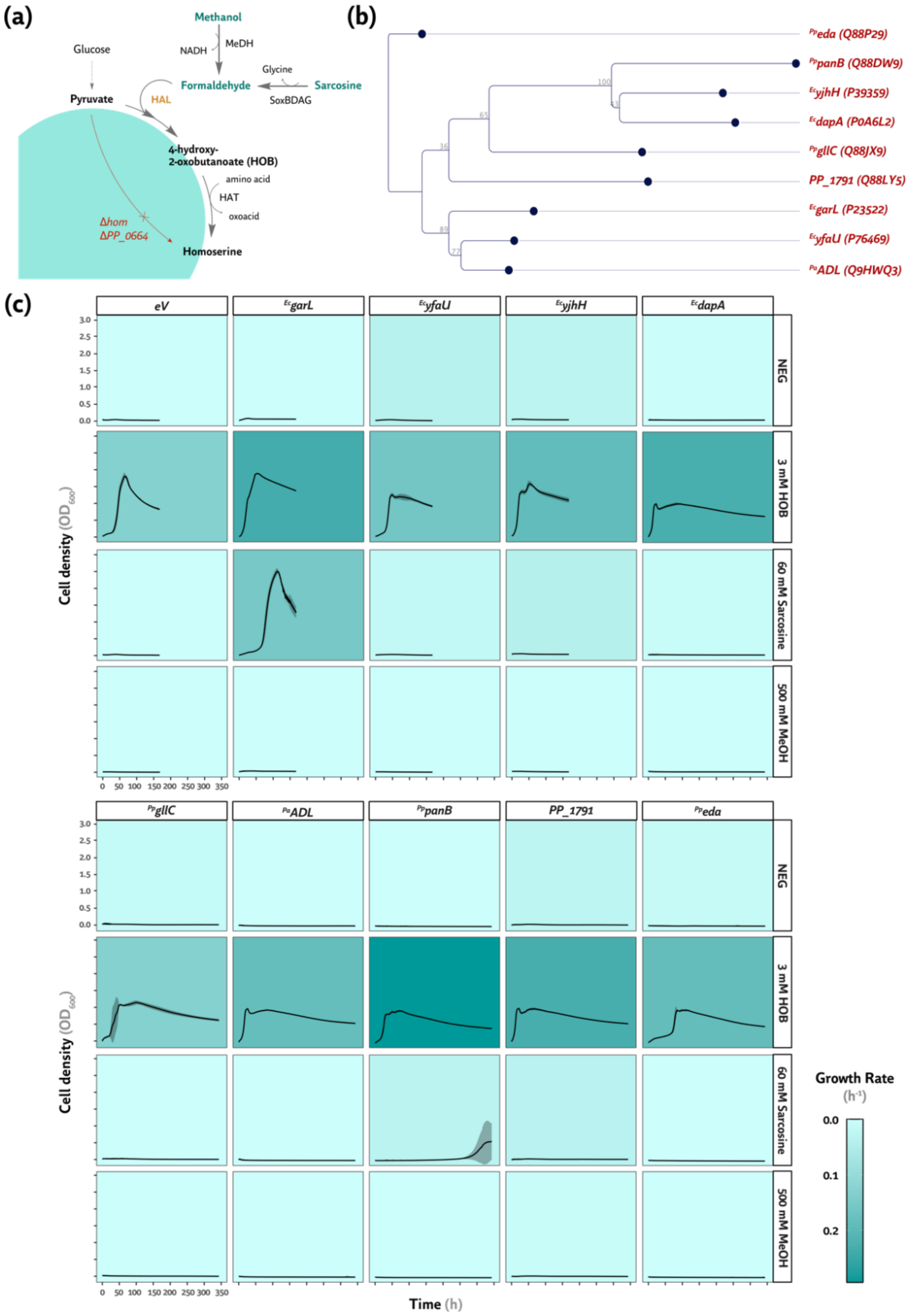


(a) Metabolic map of the auxotrophic strain Δ*hom* Δ*PP_0664* used to test the HOB aldolase (HAL) activity that assimilates formaldehyde into pyruvate to yield 4-hydroxy-2-oxobutanoate (HOB). The CT4-1 methanol dehydrogenase was also overexpressed alongside different HAL variants. (b) Phylogram representing the phylogeny of the putative HAL variants (the Uniprot code is given for each variant). (c)

Growth profiles of the Δ*hom* Δ*PP_0664* strain carrying plasmids encoding the variant library in de Bont minimal medium with 20 mM glucose supplemented with 3 mM (HOB, in its lactone form), 60 mM sarcosine, or 500 mM methanol. The specific growth rate was also calculated for each experiment and represented as a color grading. Average values for cell density (optical density measured at 600 nm, $OD_{600}$) ± standard deviation of three biological replicates are represented.

**Fig. S7. Complementation of acetyl-CoA auxotrophy *via* malate thiokinase and malyl-CoA lyase.**

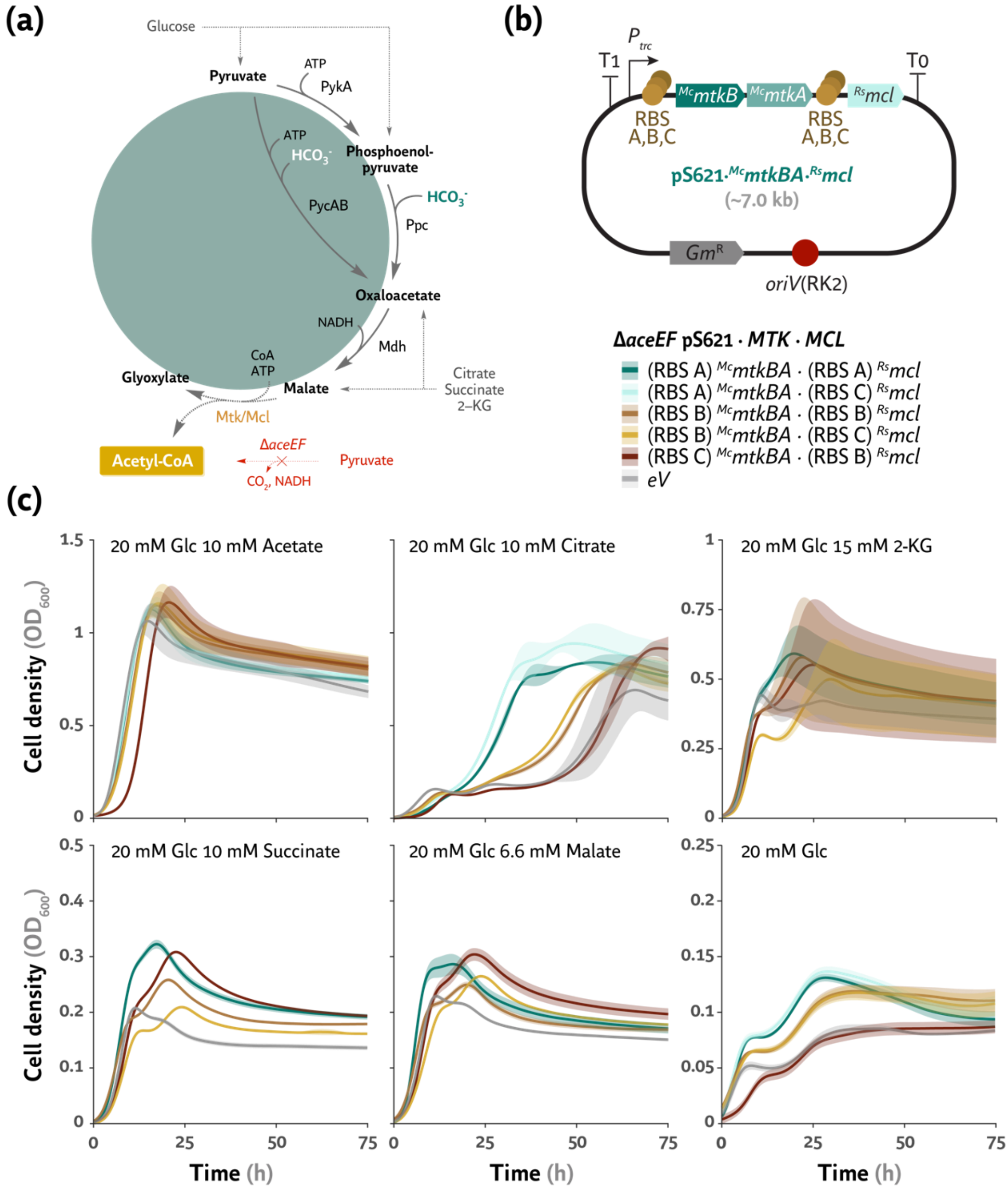


(a) Metabolic map of the Δ*aceEF* strain (lacking pyruvate dehydrogenase activity) used as an acetyl-CoA auxotroph. (b) Plasmid map of genes encoding the malate thiokinase subunits A and B (*mtkBA*) and malyl-CoA lyase (*mcl*). A small library of RBS was used for each gene with varying expression levels (A, strong; B, medium; C, weak). (c) Growth profiles of the Δ*aceEF* selection strain in de Bont minimal medium with 20 mM glucose (Glc) supplemented with acetate, citrate, 2-ketoglutarate (2-KG), succinate, or malate. All strains carried the corresponding pSEVA621 plasmid constitutively overexpressing *mtkBA* and *mcl* or the empty vector (eV). Average values for cell density (optical density measured at 600 nm, $OD_{600}$) ± standard deviation of three biological replicates are represented in all cases.

**Fig. S8. The enhanced serine-threonine cycle (eSTC), using the native PQQ-dependent methanol dehydrogenases, enables methanol assimilation at low substrate concentrations.**

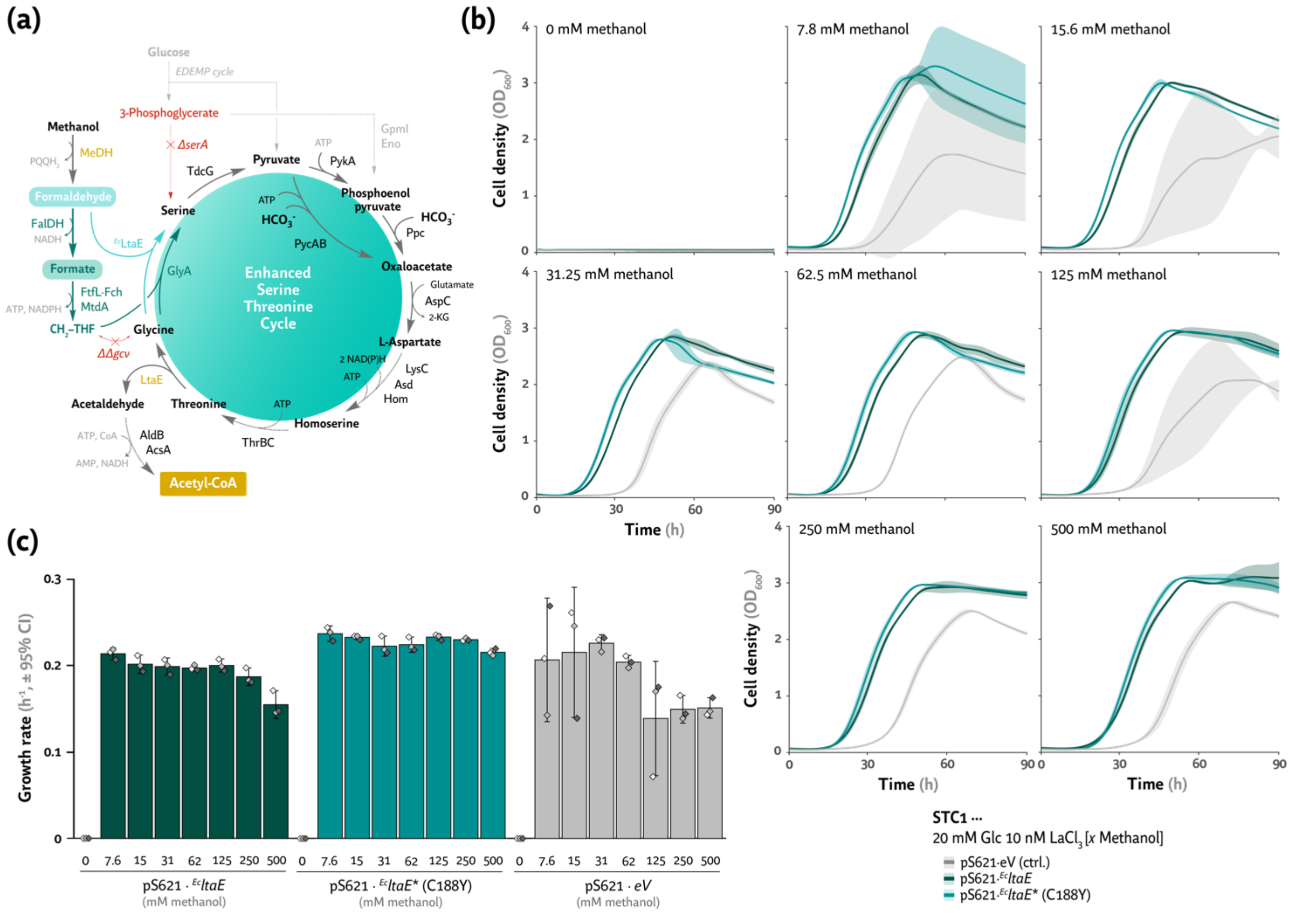


(a) Metabolic map of the serine and glycine auxotrophic strain (termed in $M_1$·STC: ΔSGG-C1) with further modifications as indicated, overexpressing the endogenous *ltaE* and *yiaY* genes. LtaE overproduction yields glycine from central carbon metabolism, whereas YiaY enables constitutive activity of endogenous PQQ-dependent alcohol dehydrogenases for methanol oxidation. This engineered *P. putida* strain was termed STC1. (b) Growth profiles of strain STC1 harboring pS621·$^{Ec}$*ltaE* / $^{Ec}$*ltaE** [C188Y (Schann et al., 2024)] in de Bont minimal medium with 20 mM glucose, 10 nM $LaCl_3$, and varying methanol concentrations. The strain carrying the empty vector (eV) is also plotted as a control. (c) Specific growth rates corresponding to the growth profiles of panel (b). Average values for the cell density (optical density measured at 600 nm, $OD_{600}$) ± standard deviation of three biological replicates are represented. Individual data points are shown. Error rates in panel (h) are plotted as specific growth rate ($\mu$, in $h^{-1}$) ± 95% confidence intervals; *P*-values were < 0.05 when error bars did not overlap.

**Table S1.** Enzyme name, function, and reaction for each (modified) serine cycle tested in this study.

| # | Enzyme[a] | Function | Reaction |
|---|---|---|---|
| **(a) Natural serine cycle (*M. extorquens* AM1)** | | | |
| 1 | MeDH | Methanol dehydrogenase | NAD + Methanol <=> **NADH** + Formaldehyde |
| 2 | Fae | 5,6,7,8-tetrahydromethanopterin hydro-lyase | 5,6,7,8-tetrahydromethanopterin + formaldehyde <=> 5,10-methylenetetrahydromethanopterin + $H_2O$ |
| 3 | MtdB | Methylenetetrahydromethanopterin dehydrogenase | 5,10-methylenetetrahydromethanopterin + NAD <=> 5,10-methenyl-5,6,7,8-tetrahydromethanopterin + **NADH** |
| 4 | Mch | Methenyltetrahydromethanopterin cyclohydrolase | 5,10-methenyl-5,6,7,8-tetrahydromethanopterin + $H_2O$ = $N_5$-formyl-5,6,7,8-tetrahydromethanopterin |
| 5 | Fhc | Formyltransferase/hydrolase complex | $N_5$-formyl-5,6,7,8-tetrahydromethanopterin <=> 5,6,7,8-tetrahydromethanopterin + Formate |
| 6 | FtfL | Formate tetrahydrofolate ligase | **ATP** + Formate + Tetrahydrofolate <=> ADP + Orthophosphate + 10-Formyltetrahydrofolate |
| 7 | Fch | Methenyltetrahydrofolate cyclohydrolase | 10-Formyltetrahydrofolate <=> 5,10-Methenyltetrahydrofolate + $H_2O$ |
| 8 | MtdA | Methylene-tetrahydrofolate synthase | **NADPH** + 5,10-Methenyltetrahydrofolate <=> NADP + 5,10-Methylenetetrahydrofolate |
| 9 | GlyA | Serine hydroxymethyltransferase (SHMT) | Glycine + 5,10-Methylenetetrahydrofolate + $H_2O$ <=> L-Serine + Tetrahydrofolate |
| 10 | Sgt | Serine-glyoxylate transaminase | Glyoxylate + L-Serine => Glycine + Hydroxypyruvate |
| 11 | Hpr | Glycerate dehydrogenase | Hydroxypyruvate + **NADH** <=> Glycerate + NAD |
| 12 | Gk | Glycerate kinase | **ATP** + Glycerate => ADP + 2-Phospho-D-glycerate |
| 13 | Eno | Enolase | 2-Phospho-D-glycerate <=> $H_2O$ + Phosphoenolpyruvate |
| 14 | Ppc | Phosphoenolpyruvate carboxylase | $HCO_3^-$ + Phosphoenolpyruvate + $H_2O$ <=> Orthophosphate + Oxaloacetate |
| 15 | Mdh | Malate dehydrogenase | **NADH** + Oxaloacetate <=> NAD + Malate |
| 16 | Mtk | Malate thiokinase | **ATP** + CoA + (S)-Malate <=> ADP + Orthophosphate + Malyl-CoA |
| 17 | Mcl | Malyl-CoA lyase | Malyl-CoA <=> Acetyl-CoA + Glyoxylate |
| **(b) Homoserine cycle (HSC)** | | | |
| 1 | MeDH | Methanol dehydrogenase | NAD + Methanol <=> **NADH** + Formaldehyde |
| 2 | SAL | Serine aldolase | Formaldehyde + Glycine <=> L-Serine |
| 3 | TdcG | Serine dehydratase | L-Serine <=> $NH_3$ + Pyruvate |
| 4 | HAL | HOB aldolase | Pyruvate + Formaldehyde <=> HOB |
| 5 | HAT | HOB aminotransferase | HOB + aminoacid <=> Homoserine + Oxoacid |
| 6 | ThrB | Homoserine kinase | Homoserine + **ATP** <=> O-phospho-homoserine + ADP |
| 7 | ThrC | Threonine synthase | O-phospho-homoserine + $H_2O$ <=> Threonine + $PO_4^-$ |
| 8 | LtaE | Low specificity L-threonine aldolase | Threonine <=> Glycine + Acetaldehyde |
| 9 | AldB | Aldehyde dehydrogenase | Acetaldehyde + **ATP** <=> Acetate + **AMP** |
| 10 | AcsA | Acetyl-coenzyme A synthetase | Acetate + CoA + NAD <=> Acetyl-CoA + **NADH** |
| **(c) Modified serine cycle (mSC)** | | | |
| 1 | MeDH | Methanol dehydrogenase | NAD + Methanol <=> **NADH** + Formaldehyde |
| 2 | FalDH | Formaldehyde dehydrogenase | NAD + Formaldehyde + $H_2O$ <=> **NADH** + Formate |
| 3 | FtfL | Formate tetrahydrofolate ligase | **ATP** + Formate + Tetrahydrofolate <=> ADP + Orthophosphate + 10-Formyltetrahydrofolate |
| 4 | Fch | Methenyltetrahydrofolate cyclohydrolase | 10-Formyltetrahydrofolate <=> 5,10-Methenyltetrahydrofolate + $H_2O$ |

| | | | |
|---|---|---|---|
| 5 | MtdA | Methylene-tetrahydrofolate synthase | **NADPH** + 5,10-Methenyltetrahydrofolate <=> NADP + 5,10-Methylenetetrahydrofolate |
| 6 | GlyA | Serine hydroxymethyltransferase (SHMT) | L-Glycine + 5,10-Methylenetetrahydrofolate + $H_2O$ <=> L-Serine + Tetrahydrofolate |
| 7 | TdcG | Serine dehydratase | L-Serine <=> $NH_3$ + Pyruvate |
| 8[b] | PycAB | Pyruvate carboxylase | **ATP** + Pyruvate + $CO_2$ <=> ADP + Orthophosphate + Oxalacetate |
| 8[c] | PykA | Phosphoenolpyruvate synthetase | **ATP** + Pyruvate + $H_2O$ <=> Orthophosphate + **AMP** + Phosphoenolpyruvate |
| 8[c] | Ppc | Phosphoenolpyruvate carboxylase | $CO_2$ + Phosphoenolpyruvate + $H_2O$ <=> Orthophosphate + Oxaloacetate |
| 9 | Mdh | Malate dehydrogenase | **NADH** + Oxaloacetate <=> NAD + Malate |
| 10 | Mtk | Malate thiokinase | **ATP** + CoA + Malate <=> ADP + Orthophosphate + Malyl-CoA |
| 11 | Mcl | Malyl-CoA lyase | Malyl-CoA <=> Acetyl-CoA + Glyoxylate |
| 12 | Agt | Alanine-glyoxylate transaminase/Glutamate-pyruvate transaminase | L-Alanine + Glyoxylate <=> Pyruvate + L-Glycine |
| **(d) Serine-threonine cycle (STC)** | | | |
| 1 | MeDH | Methanol dehydrogenase | NAD + Methanol <=> **NADH** + Formaldehyde |
| 2 | FalDH | Formaldehyde dehydrogenase | NAD + Formaldehyde + $H_2O$ <=> **NADH** + Formate |
| 3 | FtfL | Formate tetrahydrofolate ligase | **ATP** + Formate + Tetrahydrofolate <=> ADP + Orthophosphate + 10-Formyltetrahydrofolate |
| 4 | Fch | Methenyltetrahydrofolate cyclohydrolase | 10-Formyltetrahydrofolate <=> 5,10-Methenyltetrahydrofolate + $H_2O$ |
| 5 | MtdA | Methylene-tetrahydrofolate synthase | **NADPH** + 5,10-Methenyltetrahydrofolate <=> NADP + 5,10-Methylenetetrahydrofolate |
| 6 | GlyA | Serine hydroxymethyltransferase (SHMT) | Glycine + 5,10-Methylenetetrahydrofolate + $H_2O$ <=> L-Serine + Tetrahydrofolate |
| 7 | TdcG | Serine dehydratase | L-Serine <=> $NH_3$ + Pyruvate |
| 8[a] | PycAB | Pyruvate carboxylase | **ATP** + Pyruvate + $CO_2$ <=> ADP + Orthophosphate + Oxalacetate |
| 8[b1] | PykA | Phosphoenolpyruvate synthetase | **ATP** + Pyruvate + $H_2O$ <=> Orthophosphate + **AMP** + Phosphoenolpyruvate |
| 8[b2] | Ppc | Phosphoenolpyruvate carboxylase | $CO_2$ + Phosphoenolpyruvate + $H_2O$ <=> Orthophosphate + Oxaloacetate |
| 9 | AspC | Aspartate aminotransferase | Oxalacetate + Glutamate <=> L-Aspartate + 2-Ketoglutarate |
| 10 | LysC | Aspartate kinase | **ATP** + L-Aspartate <=> ADP + 4-phospho-L-aspartate |
| 11 | Asd | Aspartate-semialdehyde dehydrogenase | 4-phospho-L-aspartate + **NADPH** <=> L-Aspartate 4-semialdehyde + phosphate + NADP |
| 12 | Hom | Homoserine dehydrogenase | L-aspartate 4-semialdehyde + **NAD(P)H** <=> L-homoserine + NAD(P) |
| 13 | ThrB | Homoserine kinase | Homoserine + **ATP** <=> O-phospho-homoserine + ADP |
| 14 | ThrC | Threonine synthase | O-phospho-homoserine + $H_2O$ <=> L-Threonine + $PO_4^-$ |
| 15 | LtaE | Low specificity L-threonine aldolase | L-Threonine <=> L-Glycine + Acetaldehyde |
| 16 | AldB | Aldehyde dehydrogenase | Acetaldehyde + **ATP** <=> Acetate + **AMP** |
| 17 | AcsA | Acetyl-coenzyme A synthetase | Acetate + CoA + NAD <=> Acetyl-CoA + **NADH** |

[a] Enzymes and reactions taken from the literature [HSC, (He et al., 2020); mSC, (Yu and Liao, 2018); and STC, (Wenk et al., 2022; Yishai et al., 2017)]. Although not included in this enzyme list, given that the specific reaction remains unidentified, one NADH equivalent has been factored to replenish the amino donor for each transamination event.

[b] From methanol to acetyl-CoA *via* the shortest route (e.g., PycAB).

[c] From methanol to acetyl-CoA *via* alternative routes (e.g., PykA).

**Table S2.** Natural and modified serine cycles and their maximum-minimum driving force (MDF) compared to assimilative pathways producing $C_3$ intermediates.

| Cycle/Pathway | MDF (kJ mol$^{-1}$)[a] | | | | Product | Reaction (No MeDH)[b] |
|---|---|---|---|---|---|---|
| | No MeDH | NAD-MeDH | PQQ-MeDH | High $CO_2$ (No MeDH) | | |
| Serine cycle (SC) | 8.74 | 8.45 | 8.80 | 8.74 | Acetyl-CoA | 3 atp + co2 + coa + faldh + h2o + nadh + nadph = accoa + 3 adp + nad + nadp + 3 pi |
| Homoserine cycle (HSC) | 17.42 | 7.16 | 17.77 | – | Acetyl-CoA | CoA + 2 atp + 2 faldh + glu__L + 2 h2o + nad = accoa + adp + akg + amp + nadh + nh3 + pi + ppi |
| Modified serine cycle (mSC) | 9.34 | 9.05 | 9.41 | 9.34 | Acetyl-CoA | 3 atp + co2 + coa + faldh + glu__L + h2o + nadph = accoa + 3 adp + akg + nadp + nh3 + 3 pi |
| Serine-threonine cycle (STC) | 7.51 | 6.97 | 7.66 | 7.51 | Acetyl-CoA | CoA + 5 atp + co2 + faldh + glu__L + 3 h2o + 2 nad + 3 nadph = accoa + 4 adp + akg + amp + 2 nadh + 3 nadp + nh3 + 4 pi + ppi |
| Reductive glycine pathway (rGlyP) | 5.70 | 5.55 | 5.80 | 7.13 | Pyruvate | 2 atp + co2 + 2 faldh + h2o + nad + 2 nadph = 2 adp + nadh + 2 nadp + 2 pi + pyr |
| | 5.80 | – | – | 7.23 | Acetyl-CoA (*via* PDH) | CoA + 2 atp + 2 faldh + h2o + 2 nad + 2 nadph = accoa + 2 adp + 2 nadh + 2 nadp + 2 pi |
| Ribulose monophosphate (RuMP) pathway | 6.21 | 3.33 | 6.26 | – | Pyruvate | 3 faldh + nadp = nadph + pyr |
| | 6.26 | – | – | – | Acetyl-CoA *via* PDH) | CoA + 3 faldh + nad + nadp = accoa + co2 + nadh + nadph |

[a] The MDF value was calculated for the different $C_1$ assimilation pathways with or without the methanol oxidation reaction. When a methanol dehydrogenase (MeDH) was present, calculations considered the two types of MeDH: NADH-dependent and PQQ-dependent MeDHs. The resulting values are also plotted in **Fig. 1**g. Further details on thermodynamic profiles (No MeDH) or metabolite concentrations after MDF optimization are given in **Fig. S1** and *Materials and Methods*.

[b] The reaction given does not account for the costs of transamination or AMP phosphorylation.

**Table S3.** Specific growth rate, $C_1$ oxidation, and metabolic turnover ($CO_2$ emission) for the native serine cycle (SC) and its variants engineered in *P. putida* predicted by flux balance analysis.

| Cycle variant[a] | Module / Full Cycle | Specific growth rate ($\mu$, $h^{-1}$) | $\Delta\mu$ to SC[b] (%) | Oxidation of formate to $CO_2$ (%) | Metabolic turnover ($CO_2$ emission, mmol $g^{-1}$ $h^{-1}$) |
|---|---|---|---|---|---|
| Serine cycle (SC) | *Full cycle* | 0.606 | – | 48.1 | 5.6 |
| Homoserine cycle (HSC) | *Full cycle* | 0.621 | 2.48 | 0.0 | 5 |
| Modified serine cycle (mSC) | *Full cycle* | 0.602 | –0.66 | 48.5 | 5.8 |
| Serine-threonine cycle (STC) | *Full cycle* | 0.557 | –8.09 | 52.2 | 7.6 |

[a] Values predicted by flux balance analysis for the natural serine cycle (SC) and synthetic SC variants in the context of *P. putida* KT2440 with the methanol uptake rate set to 30 mmol $g^{-1}$ $h^{-1}$.

[b] Difference in the specific growth rate of the synthetic SC variants as compared to that supported by the natural SC, expressed as a percentage.

**Table S4.** Specific growth rate, $C_1$ oxidation, and metabolic turnover ($CO_2$ emission) for the native serine cycle (SC) and its variants engineered, as individual modules, in *P. putida* predicted by flux balance analysis at high methanol uptake rate.

| Cycle variant[a] | Module / Full cycle | Specific growth rate ($\mu$, $h^{-1}$) | Oxidation of formate to $CO_2$ (%) | Metabolic turnover ($CO_2$ emission, mmol $g^{-1}$ $h^{-1}$) |
|---|---|---|---|---|
| **Methanol uptake rate = 30 mmol $g^{-1}$ $h^{-1}$** | | | | |
| Template model (WT) | – | – | – | – |
| mSC & STC | *Module 1* | 0.496 | 57.5 | 10 |
| HSC | *Module 1* | 0.526 | 54.9 | 8.8 |
| mSC | *Module 2* | – | – | – |
| STC | *Module 2* | – | – | – |
| HSC | *Module 2* | – | – | – |
| mSC | *Full cycle* | 0.602 | 48.5 | 5.8 |
| STC | *Full cycle* | 0.557 | 52.2 | 7.6 |
| HSC | *Full cycle* | **0.564** | 0.0 | 7.3 |

[a] The specific growth rate comparison correspond to the indicated module variants and the template model (wild-type, WT). Abbreviations: mSC, modified serine cycle; STC, serine-threonine cycle; and HSC, homoserine cycle.

**Table S5.** Specific growth rate, $C_1$ oxidation, and metabolic turnover ($CO_2$ emission) for the native serine cycle (SC) and its variants engineered, as individual modules, in *P. putida* predicted by flux balance analysis in the presence of glucose.

| Cycle variant | Module / Full Cycle | Specific growth rate ($\mu$, $h^{-1}$) | $\Delta\mu$ to WT[a] (%) | Metabolic turnover ($CO_2$ emission, mmol $g^{-1}$ $h^{-1}$) |
|---|---|---|---|---|
| **Glucose uptake rate = 5 mmol $g^{-1}$ $h^{-1}$** | | | | |
| Template model (WT) | – | 0.486 | 0.00 | 10.4 |
| mSC & STC | *Module 1* | – | – | – |
| HSC | *Module 1* | – | – | – |
| mSC | *Module 2* | 0.483 | –0.62 | 10.5 |
| STC | *Module 2* | 0.474 | –2.47 | 10.9 |
| HSC | *Module 2* | – | – | – |
| mSC | *full cycle* | – | – | – |
| STC | *full cycle* | – | – | – |
| HSC | *full cycle* | – | – | – |
| **With △*aceEF* (pyruvate dehydrogenase) deletion included** | | | | |
| mSC | *Module 2* | 0.470 | –3.29 | |
| STC | *Module 2* | 0.466 | –4.12 | |

[a] The specific growth rate comparison correspond to the indicated module variants and the template model (wild-type, WT). Abbreviations: mSC, modified serine cycle; STC, serine-threonine cycle; and HSC, homoserine cycle.

**Table S6.** Specific growth rate, $C_1$ oxidation, and metabolic turnover ($CO_2$ emission) for the native serine cycle (SC) and its variants engineered, as individual modules, in *P. putida* predicted by flux balance analysis in the presence of glucose and methanol (high uptake rate).

| Cycle variant | Module / Full cycle | Specific growth rate ($\mu$, $h^{-1}$) | $\Delta\mu$ to WT[a] (%) | Oxidation of formate to $CO_2$ (%) | Metabolic turnover ($CO_2$ emission, mmol $g^{-1}$ $h^{-1}$) |
|---|---|---|---|---|---|
| **Uptake rates = 5 mmol $g^{-1}$ $h^{-1}$ (glucose) and 5 mmol $g^{-1}$ $h^{-1}$ (methanol)** | | | | | |
| Template model (WT) | – | 0.615 | 0.00 | 100.0 | 10.2 |
| mSC & STC | *Module 1* | 0.601 | –2.28 | 80.7 | 10.8 |
| HSC | *Module 1* | 0.597 | –2.93 | 88.9 | 11.0 |
| mSC | *Module 2* | 0.620 | 0.81 | 55.5 | 10.0 |
| STC | *Module 2* | 0.612 | –0.49 | 80.3 | 10.4 |
| HSC | *Module 2* | 0.606 | –1.46 | 80.5 | 10.6 |
| mSC | *full cycle* | 0.620 | 0.81 | 55.4 | 10.0 |
| STC | *full cycle* | 0.612 | –0.49 | 95.0 | 10.4 |
| HSC | *full cycle* | 0.602 | –2.11 | 95.1 | 10.8 |
| **With Δ*aceEF* (pyruvate dehydrogenase) deletion included** | | | | | |
| mSC | *Module 2* | 0.598 | –2.76 | 100.0 | 10.9 |
| STC | *Module 2* | 0.594 | –3.41 | 95.1 | 11.1 |
| HSC | *Module 2* | 0.616 | 0.16 | 55.7 | 10.2 |
| mSC | *full cycle* | 0.596 | –3.09 | 80.8 | 11.0 |
| STC | *full cycle* | 0.594 | –3.41 | 80.9 | 11.1 |
| HSC | *full cycle* | 0.616 | 0.16 | 21.1 | 10.2 |

[a] The specific growth rate comparison correspond to the indicated module variants and the template model (wild-type, WT). Abbreviations: mSC, modified serine cycle; STC, serine-threonine cycle; and HSC, homoserine cycle.

**Table S7.** Specific growth rate, $C_1$ oxidation, and metabolic turnover ($CO_2$ emission) for the native serine cycle (SC) and its variants engineered, as individual modules, in *P. putida* predicted by flux balance analysis in the presence of glucose and methanol (low uptake rate).

| Cycle variant | Module / Full cycle | Specific growth rate ($\mu$, $h^{-1}$) | $\Delta\mu$ to WT[a] (%) | Oxidation of formate to $CO_2$ (%) | Metabolic turnover ($CO_2$ emission, mmol $g^{-1}$ $h^{-1}$) |
|---|---|---|---|---|---|
| **Uptake rates = 5 mmol $g^{-1}$ $h^{-1}$ (glucose) and 2 mmol $g^{-1}$ $h^{-1}$ (methanol)** | | | | | |
| Template model (WT) | – | 0.537 | 0.00 | 100.1 | 10.4 |
| mSC & STC | *Module 1* | 0.525 | –2.23 | 57.8 | 10.9 |
| HSC | *Module 1* | 0.522 | –2.79 | 75.8 | 11.0 |
| mSC | *Module 2* | 0.535 | –0.37 | 89.0 | 10.5 |
| STC | *Module 2* | 0.526 | –2.05 | 89.2 | 10.8 |
| HSC | *Module 2* | 0.541 | 0.74 | 2.7 | 10.2 |
| mSC | *Full cycle* | 0.535 | –0.37 | 57.0 | 10.5 |
| STC | *Full cycle* | 0.530 | –1.30 | 57.4 | 10.7 |
| HSC | *Full cycle* | 0.542 | 0.93 | 2.6 | 10.2 |
| **With Δ*aceEF* (pyruvate dehydrogenase) deletion included** | | | | | |
| mSC | *Module 2* | 0.521 | –2.98 | 100.1 | 11.0 |
| STC | *Module 2* | 0.518 | –3.54 | 89.4 | 11.2 |
| HSC | *Module 2* | 0.535 | –0.37 | 0.1 | 10.5 |
| mSC | *Full cycle* | 0.519 | –3.35 | 58.3 | 11.1 |
| STC | *Full cycle* | 0.518 | –3.54 | 58.4 | 11.1 |
| HSC | *Full cycle* | 0.534 | –0.56 | 4.1 | 10.5 |

[a] The specific growth rate comparison correspond to the indicated module variants and the template model (wild-type, WT). Abbreviations: mSC, modified serine cycle; STC, serine-threonine cycle; and HSC, homoserine cycle.

**Table S8.** Bacterial strains used in this study.

| Strain | Relevant characteristics[a] | Reference or source |
|---|---|---|
| *Escherichia coli* | | |
| DH5α λ*pir* | Cloning host; F⁻ λ⁻ *endA1 glnX44(AS) thiE1 recA1 relA1 spoT1 gyrA96*(Nal$^R$) *rfbC1 deoR nupG* Φ*80(lacZ*Δ*M15)* Δ*(argF-lac)U169 hsdR17(*$r_K^-$ $m_K^+$*)*, λ*pir* lysogen | (Platt et al., 2000) |
| HB101 pRK2013 | Conjugative helper strain; *hsdR-M*$^+$*, proA2, leuB6, thi-1, recA*; harboring plasmid pRK2013, Kan$^R$ | (Figurski and Helinski, 1979) |
| *Pseudomonas putida* | | |
| EM42 | Reduced-genome derivative of *P. putida* KT2440 | (Martínez-García et al., 2014b) |
| EM42 $P_{yiaY}$::$P_{14G}$(BCD10) | Derivative of *P. putida* EM42, $P_{yiaY}$::$P_{14G}$(BCD10) | This work |
| SEM11 | Reduced-genome derivative of *P. putida* KT2440 (Belda et al., 2016) and EM42 (Martínez-García et al., 2014b) | (Wirth et al., 2023) |
| Δ*serA* | Derivative of *P. putida* SEM11, Δ*serA* (*PP_5155*) | This work |
| ΔΔ*glyA* | Derivative of *P. putida* SEM11, Δ*glyA-I* (*PP_0322*) and Δ*glyA-II* (*PP_0671*) | This work |
| Δ*ltaE* ΔΔ*glyA* | Derivative of *P. putida* ΔΔ*glyA,* Δ*ltaE* (*PP_0231*) | This work |
| Δ*ltaE* ΔΔ*glyA g*$^{Sc}$*agx1* | Derivative of *P. putida* ΔΔ*glyA* Δ*ltaE,* random integration of $P_{14G}$(BCD10)→$^{Sc}$*agx1;* Sm$^R$ | This work |
| Δ*serA* ΔΔ*glyA* Δ*frmAC* | Derivative of *P. putida* Δ*serA,* Δ*glyA-I* Δ*glyA-II* Δ*frmAC* (*PP_1616-7*) | This work |
| Δ*hom* Δ*PP_0664* | Derivative of *P. putida* SEM11, Δ*hom* (*PP_1470*) Δ*PP_0664* | This work |
| Δ*hom* Δ*PP_0664* Δ*frmAC* Δ*serA* | Derivative of *P. putida* Δ*hom* Δ*PP_0664,* Δ*frmAC* Δ*serA* | This work |
| ΔSGG-$C_1$ | Derivative of *P. putida* SEM11 Δ*serA,* Δ*gcvTHP-I* (*PP_0986-9*) Δ*gcvTHP-II* (*PP_5192-4*) *phaC1ZC2DFI* (*PP_5003-8*)::$P_4$*→*mtdA-fch-ftfL* (Turlin et al., 2022) | This work |
| Δ*aceEF* | Derivative of *P. putida* SEM11, Δ*aceEF* (*PP_0398-9*) | This work |
| STC1 | Derivative of *P. putida* ΔSGG- $C_1$, $P_{ltaE}$::$P_{trc}$ (SEVA RBS) Δ*thiO* (*PP_0612*) Δ*lapA* (*PP_0168*) Δ*lapF* (*PP_0806*) $P_{yiaY}$::$P_{14G}$(BCD10) | This work |

**Table S9.** Plasmids used in this study.

| Plasmid | Relevant characteristics[a] | Reference or source |
|---|---|---|
| pGNW2 | Suicide vector used for deletions in Gram-negative bacteria; *oriT*, *traJ*, *lacZ*$\alpha$, conditional RK6 replication origin, $P_{EM7}$→*msfGFP*; $Km^R$ | (Wirth et al., 2020) |
| pGNW2·$P_{yiaY}$::$P_{14G}$(BCD10) | Derivative of vector pGNW2 carrying homology regions to insert promoter P14G with BCD10 instead of endogenous promoter of *yiaY* (*PP_2682*); $Km^R$ | This work |
| pGNW2·Δ*serA* | Derivative of vector pGNW2 carrying homology regions to delete *serA* (*PP_5155*); $Km^R$ | (Turlin et al., 2022) |
| pGNW2·Δ*glyA-I* | Derivative of vector pGNW2 carrying homology regions to delete *glyA-I* (*PP_0322*); $Km^R$ | (Turlin et al., 2022) |
| pGNW2·Δ*glyA-II* | Derivative of vector pGNW2 carrying homology regions to delete *glyA-II* (*PP_0671*); $Km^R$ | (Turlin et al., 2022) |
| pGNW2·Δ*frmAC* | Derivative of vector pGNW2 carrying homology regions to delete *frmAC* (*PP_1616-7*); $Km^R$ | (Turlin et al., 2023) |
| pGNW2·*pha*::$P_4$*-*mtdA-fch-fttL* | Derivative of vector pGNW2 carrying homology regions to integrate $P_4$*→*mtdA-fch-fttL* from *M. extorquens* AM1 into the native *phaC1ZC2DFI* (*PP_5003-8*) locus (Turlin et al., 2022); $Km^R$ | This work |
| pGNW2·Δ*gcvTHP*-I | Derivative of vector pGNW2 carrying homology regions to delete *gcvTHP-I* (*PP_0986-9*); $Km^R$ | (Turlin et al., 2022) |
| pGNW2·Δ*gcvTHP*-II | Derivative of vector pGNW2 carrying homology regions to delete *gcvTHP-II* (*PP_5192-4*); $Km^R$ | (Turlin et al., 2022) |
| pGNW2·Δ*hom* | Derivative of vector pGNW2 carrying homology regions to delete *hom* (*PP_1470*); $Km^R$ | This work |
| pGNW2·Δ*PP_0664* | Derivative of vector pGNW2 carrying homology regions to delete *PP_0664*; $Km^R$ | This work |
| pGNW2·Δ*aceEF* | Derivative of vector pGNW2 carrying homology regions to delete *aceEF* (*PP_0338-39)*; $Km^R$ | (Wirth et al., 2022) |
| pGNW2·Δ*thiO* | Derivative of vector pGNW2 carrying homology regions to delete *thiO* (*PP_0612)*; $Km^R$ | (Turlin et al., 2022) |
| pGNW2·Δ*lapA* | Derivative of vector pGNW2 carrying homology regions to delete *lapA* (*PP_0168)*; $Km^R$ | (Calero et al., 2022) |
| pGNW2·Δ*lapF* | Derivative of vector pGNW2 carrying homology regions to delete *lapF* (*PP_0806)*; $Km^R$ | This work |
| pQURE6·H | Helper plasmid for gene deletions; conditionally-replicating vector carrying XylS/*Pm*→*I-SceI* and $P_{14G}$(BCD2)→*mRFP*; $Gm^R$ | (Volke et al., 2020) |
| pS221c | Control vector derivative of pSEVA221 (Silva-Rocha et al., 2013) harboring a $P_{14G}$ promoter and the translational coupler BCD10 followed by a *START* and *STOP* codon (ATGTAA); *oriT oriV*(RK2); $Kan^R$ | This work |
| pS221·$^{Pp}$*ltaE* | Derivative of vector pSEVA221 harboring a constitutive promoter and translational coupler $P_{14G}$(BCD10)→ $^{Pp}$*ltaE* (PP_0321); *oriT oriV*(RK2); $Km^R$ | This work |
| pS221·$^{Pp}$*ilvE* | Derivative of vector pSEVA221 harboring a constitutive promoter and translational coupler $P_{14G}$(BCD10)→ $^{Pp}$*ilvE* (PP_3511); *oriT oriV*(RK2); $Km^R$ | This work |
| pS221·$^{Ec}$*ilvE* | Derivative of vector pSEVA221 harboring a constitutive promoter and translational coupler $P_{14G}$(BCD10)→ $^{Ec}$*ilvE* (JW5606); *oriT oriV*(RK2); $Km^R$ | This work |
| pS221·$^{Ec}$*alaC** | Derivative of vector pSEVA221 harboring a constitutive promoter and translational coupler $P_{14G}$(BCD10)→ $^{Ec}$*alaC* A142P Y275D (Bouzon et al., 2017); *oriT oriV*(RK2); $Km^R$ | This work |
| pS221·$^{Ec}$*aspC* | Derivative of vector pSEVA221 harboring a constitutive promoter and translational coupler $P_{14G}$(BCD10)→ $^{Ec}$*aspC* (JW0911); *oriT oriV*(RK2); $Km^R$ | This work |
| pS221·$^{Pp}$*thiO* | Derivative of vector pSEVA221 harboring a constitutive promoter and translational coupler $P_{14G}$(BCD10)→ $^{Pp}$*thiO* (PP_0612); *oriT oriV*(RK2); $Km^R$ | This work |

| | | |
|---|---|---|
| pS221·$^{Hs}$*agxt1* | Derivative of vector pSEVA221 harboring a constitutive promoter and translational coupler $P_{14G}$(BCD10)→*agxt1* from *H. sapiens* (codon-optimized) (Yu and Liao, 2018); *oriT oriV*(RK2); Km$^{R}$ | This work |
| pS221·$^{Sc}$*agt1* | Derivative of vector pSEVA221 harboring a constitutive promoter and translational coupler $P_{14G}$(BCD10)→*agt1* from *S. cerevisiae* (codon-optimized) (Yu and Liao, 2018); *oriT oriV*(RK2); Km$^{R}$ | This work |
| pS221·$^{Pb}$*bhcA* | Derivative of vector pSEVA221 harboring a constitutive promoter and translational coupler $P_{14G}$(BCD10)→*bhcA* from *P. denitrificans* (*Pden_3921*) (Schada von Borzyskowski et al., 2023); *oriT oriV*(RK2); Km$^{R}$ | This work |
| pS621c | Control vector derivative of pSEVA621 (Silva-Rocha et al., 2013) harboring a $P_{trc}$ promoter and the canonical SEVA ribosome binding site (RBS) followed by a *START* and *STOP* codon (ATGTAA); *oriT oriV*(RK2); Gm$^{R}$ | This work |
| pS621·$^{Bs}$*mdh* | Derivative of vector pSEVA621 harboring a $P_{trc}$ promoter and the canonical SEVA ribosome binding site (RBS)→methanol dehydrogenase from *Bacillus stearothermophilus* (Whitaker et al., 2017); *oriT oriV*(RK2); Gm$^{R}$ | This work |
| pS621·$^{Cn}$*mdh* | Derivative of vector pSEVA621 harboring a $P_{trc}$ promoter and the canonical SEVA ribosome binding site (RBS)→methanol dehydrogenase CT4-1 from *C. necator* N-1 (Wu et al., 2016); *oriT oriV*(RK2); Gm$^{R}$ | This work |
| pS621·*yiaY* | Derivative of vector pSEVA621 harboring a $P_{trc}$ promoter and the canonical SEVA ribosome binding site (RBS)→*yiaY* (*PP_2682*); *oriT oriV*(RK2); Gm$^{R}$ | This work |
| pS621·*pedE* | Derivative of vector pSEVA621 harboring a $P_{trc}$ promoter and the canonical SEVA ribosome binding site (RBS)→*pedE* (*PP_2674*); *oriT oriV*(RK2); Gm$^{R}$ | This work |
| pS621·*pedH* | Derivative of vector pSEVA621 harboring a $P_{trc}$ promoter and the canonical SEVA ribosome binding site (RBS)→*pedH* (*PP_2679*); *oriT oriV*(RK2); Gm$^{R}$ | This work |
| pS621·$^{Pp}$*adhP* | Derivative of vector pSEVA621 harboring a $P_{trc}$ promoter and the canonical SEVA ribosome binding site (RBS)→*adhP* (*PP_3839*); *oriT oriV*(RK2); Gm$^{R}$ | This work |
| pS621·$^{Cg}$*mdh* | Derivative of vector pSEVA621 harboring a $P_{trc}$ promoter and the canonical SEVA ribosome binding site (RBS)→methanol dehydrogenase from *C. glutamicum* (Uniprot: A4QHJ5) (Kotrbova-Kozak et al., 2007); *oriT oriV*(RK2); Gm$^{R}$ | This work |
| pS621·$^{Bm}$*mdh2* | Derivative of vector pSEVA621 harboring a $P_{trc}$ promoter and the canonical SEVA ribosome binding site (RBS)→methanol dehydrogenase from *B. methanolicus* carrying mutations Q5L A363L (Uniprot: I3DVX6) (Roth et al., 2019); *oriT oriV*(RK2); Gm$^{R}$ | This work |
| pS621·$^{Me}$*xoxFGJ* | Derivative of vector pSEVA621 harboring a $P_{trc}$ promoter and the canonical SEVA ribosome binding site (RBS)→methanol dehydrogenase complex *xoxFGJ* from *M. extorquens* AM1 (*MexAM1_META1p1740-2*) (Featherston et al., 2019; Good et al., 2019); *oriT oriV*(RK2); Gm$^{R}$ | This work |
| pS621·$^{Ec}$*ltaE* | Derivative of vector pSEVA621 harboring a $P_{trc}$ promoter and the canonical SEVA ribosome binding site (RBS)→ $^{Ec}$*ltaE*; *oriT oriV*(RK2); Gm$^{R}$ | This work |
| pS621·$^{Ec}$*ltaE** C188Y | Derivative of vector pSEVA621 harboring a $P_{trc}$ promoter and the canonical SEVA ribosome binding site (RBS)→ $^{Ec}$*ltaE** (C188Y) (Schann et al., 2024); *oriT oriV*(RK2); Gm$^{R}$ | This work |
| pS621·$^{Cn}$*mdh*·$^{Ec}$*ltaE* | Derivative of vector pSEVA621 harboring a $P_{trc}$ promoter and the canonical SEVA ribosome binding site (RBS)→$^{Cn}$*mdh:*BCD10*:* $^{Ec}$*ltaE*; *oriT oriV*(RK2); Gm$^{R}$ | This work |
| pS621·$^{Cn}$*mdh*·$^{Pp}$*ltaE* | Derivative of vector pSEVA621 harboring a $P_{trc}$ promoter and the canonical SEVA ribosome binding site (RBS)→ $^{Cn}$*mdh:*BCD10*:* $^{Pp}$*ltaE*; *oriT oriV*(RK2); Gm$^{R}$ | This work |
| pS621·$^{Cn}$*mdh*·$^{Yl}$*ltaE* | Derivative of vector pSEVA621 harboring a $P_{trc}$ promoter and the canonical SEVA ribosome binding site (RBS)→ $^{Cn}$*mdh:BCD10:ltaE* from *Y. lipolytica* (Uniprot Q6CG81); *oriT oriV*(RK2); Gm$^{R}$ | This work |

| | | |
|---|---|---|
| pS621·$^{Cn}$*mdh*·$^{Cn}$*ltaE* | Derivative of vector pSEVA621 harboring a $P_{trc}$ promoter and the canonical SEVA ribosome binding site (RBS)→ $^{Cn}$*mdh:BCD10:ltaE* from *C. necator* (Uniprot Q0K830); *oriT oriV*(RK2); Gm$^{R}$ | This work |
| pS621· $^{Cn}$*mdh*·$^{Me}$*ltaE* | Derivative of vector pSEVA621 harboring a $P_{trc}$ promoter and the canonical SEVA ribosome binding site (RBS)→$^{Cn}$*mdh:BCD10:ltaE* from *M. extorquens* AM1 (Uniprot C7CBZ4); *oriT oriV*(RK2); Gm$^{R}$ | This work |
| pS621·$^{Cn}$*mdh*·$^{Vn}$*ltaE* | Derivative of vector pSEVA621 harboring a $P_{trc}$ promoter and the canonical SEVA ribosome binding site (RBS)→$^{Cn}$*mdh:BCD10:ltaE* from *V. natriegens* (Uniprot A0A1B1EKL1); *oriT oriV*(RK2); Gm$^{R}$ | This work |
| pS621·$^{Cn}$*mdh*·$^{Sc}$*ltaE* | Derivative of vector pSEVA621 harboring a $P_{trc}$ promoter and the canonical SEVA ribosome binding site (RBS)→$^{Cn}$*mdh:BCD10:ltaE* from *S. cerevisiae* (Uniprot P37303); *oriT oriV*(RK2); Gm$^{R}$ | This work |
| pS621·$^{Cn}$*mdh*·$^{Kp}$*ltaE* | Derivative of vector pSEVA621 harboring a $P_{trc}$ promoter and the canonical SEVA ribosome binding site (RBS)→$^{Cn}$*mdh:BCD10:ltaE* from *K. phaffii* (Uniprot C4R6W2); *oriT oriV*(RK2); Gm$^{R}$ | This work |
| pS621·$^{Cn}$*mdh*·$^{Pp}$*eda* | Derivative of vector pSEVA621 harboring a $P_{trc}$ promoter and the canonical SEVA ribosome binding site (RBS)→$^{Cn}$*mdh:BCD10:eda* (*PP_1024*); *oriT oriV*(RK2); Gm$^{R}$ | This work |
| pS621·$^{Cn}$*mdh*·$^{Pp}$*panB* | Derivative of vector pSEVA621 harboring a $P_{trc}$ promoter and the canonical SEVA ribosome binding site (RBS)→$^{Cn}$*mdh:BCD10:panB* (*PP_4699*); *oriT oriV*(RK2); Gm$^{R}$ | This work |
| pS621·$^{Cn}$*mdh*·$^{Ec}$*yjhH* | Derivative of vector pSEVA621 harboring a $P_{trc}$ promoter and the canonical SEVA ribosome binding site (RBS)→$^{Cn}$*mdh:BCD10:yjhH* from *E. coli* (Uniprot P39359) (He et al., 2020); *oriT oriV*(RK2); Gm$^{R}$ | This work |
| pS621·$^{Cn}$*mdh*·$^{Ec}$*dapA* | Derivative of vector pSEVA621 harboring a $P_{trc}$ promoter and the canonical SEVA ribosome binding site (RBS)→$^{Cn}$*mdh:BCD10:dapA* from *E. coli* (Uniprot P0A6L2); *oriT oriV*(RK2); Gm$^{R}$ | This work |
| pS621·$^{Cn}$*mdh*·$^{Pp}$*gllC* | Derivative of vector pSEVA621 harboring a $P_{trc}$ promoter and the canonical SEVA ribosome binding site (RBS)→$^{Cn}$*mdh:BCD10:gllC* (*PP_2514*); *oriT oriV*(RK2); Gm$^{R}$ | This work |
| pS621·$^{Cn}$*mdh*·*PP_1791* | Derivative of vector pSEVA621 harboring a $P_{trc}$ promoter and the canonical SEVA ribosome binding site (RBS)→$^{Cn}$*mdh:BCD10:PP_1791*; *oriT oriV*(RK2); Gm$^{R}$ | This work |
| pS621·$^{Cn}$*mdh*·$^{Ec}$*garL* | Derivative of vector pSEVA621 harboring a $P_{trc}$ promoter and the canonical SEVA ribosome binding site (RBS)→$^{Cn}$*mdh:BCD10:garL* from *E. coli* (Uniprot P23522); *oriT oriV*(RK2); Gm$^{R}$ | This work |
| pS621·$^{Cn}$*mdh*·$^{Ec}$*yfaU* | Derivative of vector pSEVA621 harboring a $P_{trc}$ promoter and the canonical SEVA ribosome binding site (RBS)→$^{Cn}$*mdh:BCD10:yfaU* from *E. coli* (Uniprot P76469); *oriT oriV*(RK2); Gm$^{R}$ | This work |
| pS621·$^{Cn}$*mdh*·$^{Pa}$*ADL* | Derivative of vector pSEVA621 harboring a $P_{trc}$ promoter and the canonical SEVA ribosome binding site (RBS)→$^{Cn}$*mdh:BCD10:ADL* from *P. aeruginosa PAO1* (Uniprot Q9HWQ3); *oriT oriV*(RK2); Gm$^{R}$ | This work |
| pBAMD1-4 | Tn*5* delivery plasmid; conditional RK6 replication origin; Sm$^{R}$, Amp$^{R}$ | (Martínez-García et al., 2014a) |
| pBAMD1-4-$P_{14G}$ (BCD10)→$^{Sc}$*agx1* | Derivative of vector pBAMD1-4 carrying a $P_{14G}$(BCD10)→$^{Sc}$*agx1* module from *S. cerevisiae* (codon-optimized); Sm$^{R}$, Amp$^{R}$ | This work |
| pS621 · (RBS A) $^{Mc}$*mtkBA* · (RBS A) $^{Rs}$*mcl* | Derivative of vector pSEVA621 harboring a $P_{trc}$ promoter and strong RBS (A)→$^{Mc}$*mtkB;* RBS A→$^{Mc}$*mtkB;* RBS A→$^{Rs}$*mcl*, codon-optimized from *M. capsulatus* (Uniprot Q607L9; Q607L8) and *R. sphaeroides* (Uniprot Q3J5L6), respectively ((as in (Orsi et al., 2024)); Gm$^{R}$ | This work |
| pS621 · (RBS A) $^{Mc}$*mtkBA* · (RBS C) $^{Rs}$*mcl* | Derivative of vector pSEVA621 harboring a $P_{trc}$ promoter and strong RBS (A)→$^{Mc}$*mtkB;* RBS A→$^{Mc}$*mtkB;* RBS C→$^{Rs}$*mcl,* codon-optimized from *M.* | This work |

| | | |
|---|---|---|
| | *capsulatus* (Uniprot Q607L9; Q607L8) and *R. sphaeroides* (Uniprot Q3J5L6), respectively (as in (Orsi et al., 2024)); $Gm^R$ | |
| pS621 · (RBS B) $^{Mc}$*mtkBA* · (RBS B) $^{Rs}$*mcl* | Derivative of vector pSEVA621 harboring a $P_{trc}$ promoter and medium RBS (B)→$^{Mc}$*mtkB;* RBS B→$^{Mc}$*mtkB;* RBS B→$^{Rs}$*mcl,* codon-optimized from *M. capsulatus* (Uniprot Q607L9; Q607L8) and *R. sphaeroides* (Uniprot Q3J5L6), respectively (as in (Orsi et al., 2024)); $Gm^R$ | This work |
| pS621 · (RBS B) $^{Mc}$*mtkBA* · (RBS C) $^{Rs}$*mcl* | Derivative of vector pSEVA621 harboring a $P_{trc}$ promoter and medium RBS (B)→$^{Mc}$*mtkB;* RBS B→$^{Mc}$*mtkB;* RBS C→$^{Rs}$*mcl,* codon-optimized from *M. capsulatus* (Uniprot Q607L9; Q607L8) and *R. sphaeroides* (Uniprot Q3J5L6), respectively (as in (Orsi et al., 2024)); $Gm^R$ | This work |
| pS621 · (RBS C) $^{Mc}$*mtkBA* · (RBS B) $^{Rs}$*mcl* | Derivative of vector pSEVA621 harboring a $P_{trc}$ promoter and weak RBS (C)→ $^{Mc}$*mtkB;* RBS C→$^{Mc}$*mtkB;* RBS B→$^{Rs}$*mcl,* codon-optimized from *M. capsulatus* (Uniprot Q607L9; Q607L8) and *R. sphaeroides* (Uniprot Q3J5L6), respectively (as in (Orsi et al., 2024)); $Gm^R$ | This work |

[a] Antibiotic markers: Gm, gentamicin; Km, kanamycin; and Sm, streptomycin.

**Table S10.** Oligonucleotides used in this study.

| Name | DNA sequence (5'→3') | Use |
|---|---|---|
| pGNW2_P4*_F | ATAATGGAUCCAGCTCTAGAATAGATATCCTAT | Construction of pGNW2·*pha*::P4$^{evo}$-*mtdA-fch-ftfL* |
| pGNW2_P4*_R | ATCCATTAUACAGAAAAATTTTCCTGATGTCAA | |
| hom_US_U_F | AGATCCUACGGCCTCTATGAAGTCA | Construction of pGNW2·*hom* |
| hom_US_U_R | ACACTGUGAACTCCCCATTGAACGG | |
| hom_DS_U_F | ACAGTGUAACAAATATTGCCGACGG | |
| hom_DS_U_R | AGGTCGACUGATGTTGTGGATGTTGTCG | |
| yiaY_gProm_Up_F | AGATCCUGCAAGCTCGTTCCAAAGT | Construction of pGNW2·$P_{yiaY}$::$P_{14G}$(BCD10) |
| yiaY_gProm_Up_R | ATGGGCCGGUAAGCCTGTTCTTATTGTTCTG | |
| 14gBCD10_yiaY_gProm_Up_F | ACCGGCCCAUTGACAAGGCTCTCGCGGC | |
| 14gBCD10_yiaY_gProm_Up_R | AGAAACGAUCCTCCGCATGATTAAGATGTTTCA | |
| yiaY_gProm_down_F | ATCGTTTCUAATGAGCCAGAGTTTCAGCC | |
| yiaY_gProm_down_R | AGGTCGACUCTCGATGGCATGCACCAA | |
| PP_0664_US_U_F | AGATCCUTTCATGCCTTGTTGACCCG | Construction of pGNW2·*PP_0664* |
| PP_0664_US_U_R | ACATGGUTTGCTGTCCTGTCCAGGC | |
| PP_0664_DS_U_F | ACCATGUGAGGAGCAACGTTCATGAG | |
| PP_0664_DS_U_R | AGGTCGACUTTTCAGGGCAGAAAGCGG | |
| pS221_PPLTAE_F | AACTAGUCTTGGACTCCTGTTGATAGAT | Construction of pS221· $^{Pp}$*ltaE* |
| pS221_PPLTAE_R | AGAAACGAUCCTCCGCATGATTAAGATG | |
| PPLTAE_pS221_F | ATCGTTTCUAATGACAGACAAGAGCCAAC | |
| PPLTAE_pS221_R | ACTAGTUCAGCCACCAATGATCGTG | |
| pS221_PPILVE_F | AACTAGUCTTGGACTCCTGTTGATAGAT | Construction of pS221· $^{Pp}$*ilvE* |
| pS221_PPILVE_R | AGAAACGAUCCTCCGCATGATTAAGATG | |
| PPILVE_pS221_F | ATCGTTTCUAATGAGCAACGAAAGCATTAATT | |
| PPILVE_pS221_R | ACTAGTUCAGGCGACCTTGACGATC | |
| pS221_ECILVE_F | ATAAACUAGTCTTGGACTCCTGTTGATAG | Construction of pS221· $^{Ec}$*ilvE* |
| pS221_ECILVE_R | AGAAACGAUCCTCCGCATGATTAAGATG | |
| ECILVE_pS221_F | ATCGTTTCUAATGACCACGAAGAAAGCT | |
| ECILVE_pS221_R | AGTTTAUTGATTAACTTGATCTAACCAGC | |
| pS221_ECALAC*_F | ATAAACUAGTCTTGGACTCCTGTTGATAGA | Construction of pS221· $^{Ec}$*alaC** |
| pS221_ECALAC*_R | AGCCATUAGAAACGATCCTCCGCATGA | |
| ECALAC*_pS221_F | AATGGCUGACACTCGCCCTGAACGT | |
| ECALAC*_pS221_R | AGTTTAUCCGCGTTTTCGTGAATATGT | |
| pS221_ECASPC_F | AAACTAGUCTTGGACTCCTGTTGATAGATCCA | Construction of pS221· $^{Ec}$*aspC* |
| pS221_ECASPC_R | AAACATUAGAAACGATCCTCCGCATGA | |
| ECASPC_pS221_F | AATGTTUGAGAACATTACCGCCGCT | |
| ECASPC_pS221_R | ACTAGTTUACAGCACTGCCACAATCG | |
| pS221_THIO_F | AACTAGUCTTGGACTCCTGTTGATAGAT | Construction of pS221· $^{Pp}$*thiO* |
| pS221_THIO_R | AGAAACGAUCCTCCGCATGATTAAGATG | |
| THIO_pS221_F | ATCGTTTCUAATGAGCAAGCAAGTAGTGGTGGTT | |
| THIO_pS221_R | ACTAGTUCAGCCCAAACGCCCTTCTG | |
| pS221_HSAGXT1_F | ACTGTAAACUAGTCTTGGACTCCTGTTGATAGA | Construction of pS221· $^{Hs}$*agxt1* |
| pS221_HSAGXT1_R | AGGCCATUAGAAACGATCCTCCGCATGA | |
| HSAGXT1_pS221_F | AATGGCCUCCCACAAATTGCTGGTTA | |
| HSAGXT1_pS221_R | AGTTTACAGUTCTTCTTTGGGCAGTGT | |
| pS221_SCAGT1_F | AAGTAAACUAGTCTTGGACTCCTGTTGATAGA | Construction of pS221· $^{Sc}$*agt1* |
| pS221_SCAGT1_R | AGAAACGAUCCTCCGCATGATTAAGATG | |
| SCAGT1_pS221_F | ATCGTTTCUAATGACCAAAAGCGTCGATACG | |
| SCAGT1_pS221_R | AGTTTACTUTTTGCGTTGCAGGGCTAG | |
| pS221_BHCA_F | AACTAGUCTTGGACTCCTGTTGATAGAT | Construction of |

| | | |
|---|---|---|
| pS221_BHCA_R | AGAAACGAUCCTCCGCATGATTAAGATG | pS221· $^{Pb}$*bhcA* |
| BHCA_ pS221_F | ATCGTTTCUAATGACCAGCCAGAACCCGATCT | |
| BHCA_ pS221_R | ACTAGTUCAGGCGGCTTTCTTCTGCG | |
| pS621_BSMDH_F | AGGATTAAUCTAGAGTCGACCTGCAGG | |
| pS621_BSMDH_R | ATATGTUTTTCCTCCTGGGAATTCG | Construction of |
| BSMDH_ pS621_F | AACATAUGAAAGCCGCGGTGGTCAA | pS621· $^{Bs}$*mdh* |
| BSMDH_ pS621_R | ATTAATCCUCTTTGAGCTTGAGCACGAT | |
| pS621_CNMDH_F | AGGATTAAUCTAGAGTCGACCTGCAGG | |
| pS621_CNMDH_R | ATATGTUTTTCCTCCTGGGAATTCG | Construction of |
| CNMDH_ pS621_F | AACATAUGAAAGCCGCGGTGGTCAA | pS621· $^{Cn}$*mdh* |
| CNMDH_ pS621_R | ATTAATCCUCTTTGAGCTTGAGCACGAT | |
| pS621_YIAY_F | AGGCCCTCUAATCTAGAGTCGACCTGC | |
| pS621_YIAY_R | ATATGTUTTTCCTCCTGGGAATTCG | Construction of |
| YIAY_ pS621_F | AACATAUGAGCCAGAGTTTCAGCCC | pS621·*yiaY* |
| YIAY_ pS621_R | AGAGGGCCUCGCCATAGACCACCTCGA | |
| pS621_PEDE_F | ACGTTGAUCTAGAGTCGACCTGCAGG | |
| pS621_PEDE_R | ATATGTUTTTCCTCCTGGGAATTCG | Construction of |
| PEDE_pS621_F | AACATAUGACAATAAGATCGCTACCC | pS621·*pedE* |
| PEDE_pS621_R | ATCAACGUTGTGCAGTCTTGTTGTCC | |
| pS621_PEDH_F | ATAATCUAGAGTCGACCTGCAGGCA | |
| pS621_PEDH_R | ATATGTUTTTCCTCCTGGGAATTCG | Construction of |
| PEDH_pS621_F | AACATAUGACCCGATCCCCACGTCG | pS621·*pedH* |
| PEDH_pS621_R | AGATTAUGGCTTGACGCTTGCCGTT | |
| pS621_ADHP_F | AGGCTGAUCTAGAGTCGACCTGCAGG | |
| pS621_ADHP_R | ATATGTUTTTCCTCCTGGGAATTCGC | Construction of |
| ADHP_pS621_F | AACATAUGAAAGCTGCTGTCGTTGCA | pS621· $^{Pp}$*adhP* |
| ADHP_pS621_R | ATCAGCCUTCGAACTGTATCACCATCCGG | |
| pS621_CGMDH_F | ACTAATCUAGAGTCGACCTGCAGGCA | |
| pS621_CGMDH_R | ATATGTUTTTCCTCCTGGGAATTCGC | Construction of |
| CGMDH_pS621_F | AACATAUGCATCATCACCATCACCA | pS621· $^{Cg}$*mdh* |
| CGMDH_pS621_R | AGATTAGUAACGGATAGCAACACGAC | |
| pS621_BMMDH_F | ATGTAATCUAGAGTCGACCTGCAGGCA | |
| pS621_BMMDH_R | ATATGTUTTTCCTCCTGGGAATTCGC | Construction of |
| BMMDH_pS621_F | AACATAUGCATCATCACCATCACCA | pS621· $^{Bm}$*mdh*$^{*}$ |
| BMMDH_pS621_R | AGATTACAUAGCGTTTTTGATGATCTG | |
| pS621_MEXOXFGJ_F | ATGATCUAGAGTCGACCTGCAGGCA | |
| pS621_MEXOXFGJ_R | ATATGTUTTTCCTCCTGGGAATTCGC | Construction of |
| MEXOXFGJ_pS621_F | AACATAUGCATCATCACCACCACCATCGTGC | pS621· $^{Me}$*xoxFGJ* |
| MEXOXFGJ_pS621_R | AGATCAUTCGTCGGCCGCGTCGAGA | |
| pS621_ECLTAE_F | ATCTAGAGUCGACCTGCAGGCATGCAA | |
| pS621_ECLTAE_R | ATCATATGUTTTTCCTCCTGGGAATTCGC | Construction of |
| ECLTAE_pS621_F | ACATATGAUTGATTTACGCAGTGATACCGTTA | pS621· $^{Ec}$*ltaE* |
| ECLTAE_pS621_R | ACTCTAGAUTAACGCGCCAGGAATGCA | |
| pS621_CNMDH_ECLTAE_F | ACCAGGCAUCAAATAAAATAGAGTCGACCTG | |
| pS621_CNMDH_ECLTAE_R | AGAAACGAUCCTCCGCATGATTAAGAT | Construction of |
| CNMDH_ECLTAE_pS621_F | ATCGTTTCUATGATTGATTTACGCAGTGATACC | pS621· $^{Cn}$*mdh* · $^{Ec}$*ltaE* |
| CNMDH_ECLTAE_pS621_R | ATGCCTGGUTAAGCTTAACGCGCCAGGAATGC | |
| GG_PPLTAE_F | AACGTCTCGCTCGAATGACAGACAAGAGCCAACA | Construction of |
| GG_PPLTAE_R | TTCGTCTCCCTCAAAGCGCCACCAATGATCGTGCG | pS621· $^{Cn}$*mdh* · $^{Pp}$*ltaE** |
| pS621_CNMDH_YLLTAE_F | ATCTAAGCUTAACCAGGCATCAAATAAAATAGA | |
| pS621_CNMDH_YLLTAE_R | AGAAACGAUCCTCCGCATGATTAAGAT | Construction of |
| CNMDH_YLLTAE_pS621_F | ATCGTTTCUATGACTGCCTGCTCTCAC | pS621· $^{Cn}$*mdh* · $^{Yl}$*ltaE* |
| CNMDH_YLLTAE_pS621_R | AGCTTAGAUGTAGCCACTAGTCTTAAGCT | |
| pS621_CNMDH_CNLTAE_F | ACCAGGCAUCAAATAAAATAGAGTCGACCTG | |
| pS621_CNMDH_CNLTAE_R | AGAAACGAUCCTCCGCATGATTAAGAT | Construction of |
| CNMDH_CNLTAE_pS621_F | ATCGTTTCUATGCAGCACTTCGCCTCTGACAAC | pS621· $^{Cn}$*mdh* · $^{Cn}$*ltaE* |
| CNMDH_CNLTAE_pS621_R | ATGCCTGGUTCAGGCGCCGCAAGCCGC | |

| | | |
|---|---|---|
| pS621_CNMDH_MELTAE_F | ACCAGGCAUCAAATAAAATAGAGTCGACCTG | Construction of pS621·$^{Cn}$*mdh*·$^{Me}$*ltaE* |
| pS621_CNMDH_MELTAE_R | AGAAACGAUCCTCCGCATGATTAAGAT | |
| CNMDH_MELTAE_ pS621_F | ATCGTTTCUGTGGCCGAACAGCAATTCGCCAGC | |
| CNMDH_MELTAE_ pS621_R | ATGCCTGGUTCACGCCGCCGCGGGCAG | |
| pS621_CNMDH_VNLTAE_F | ATTCTCUGCTTAACCAGGCATCAAATAA | Construction of pS621·$^{Cn}$*mdh*·$^{Vn}$*ltaE* |
| pS621_CNMDH_VNLTAE_R | AGTCCAUAGAAACGATCCTCCGCAT | |
| CNMDH_VNLTAE_ pS621_F | ATGGACUTTCGATCTGATACAGTAA | |
| CNMDH_VNLTAE_ pS621_R | AGAGAAUAGATTTCAACGCAGATAA | |
| pS621_CNMDH_SCLTAE_F | ACTGAGCTUAACCAGGCATCAAATAAAATAGA | Construction of pS621·$^{Cn}$*mdh*·$^{Sc}$*ltaE* |
| pS621_CNMDH_SCLTAE_R | AGAAACGAUCCTCCGCATGATTAAGAT | |
| CNMDH_SCLTAE_ pS621_F | ATCGTTTCUATGACTGAATTCGAATTG | |
| CNMDH_SCLTAE_ pS621_R | AAGCTCAGUATTTGTAGGTTTTTATTTCG | |
| pS621_CNMDH_KPLTAE_F | ACTAAGCTUAACCAGGCATCAAATAAAATAGA | Construction of pS621·$^{Cn}$*mdh*·$^{Kp}$*ltaE* |
| pS621_CNMDH_KPLTAE_R | AGAAACGAUCCTCCGCATGATTAAGAT | |
| CNMDH_KPLTAE_ pS621_F | ATCGTTTCUATGACAAAGGAAGATTTT | |
| CNMDH_KPLTAE_ pS621_R | AAGCTTAGUTTTTCAAAACTGGAGAAT | |
| GG_EDA_F | AACGTCTCGCTCGAATGCCCATGAGCCAAGGA | Construction of pS621·$^{Cn}$*mdh*·$^{Pp}$*eda** |
| GG_EDA_R | TTCGTCTCCCTCAAAGCGTTGGCGTCCAGCAGTGC | |
| GG_PANB_F | AACGTCTCGCTCGAATGCCTGAAGTAACCCTT | Construction of pS621·$^{Cn}$*mdh*·$^{Pp}$*panB** |
| GG_PANB_R | TTCGTCTCCCTCAAAGCTGCACTGAACCCGTGTTC | |
| pS621_CNMDH_YJHH_F | AGTCTGAGCUTAACCAGGCATCAAATAAAATAGA | Construction of pS621·$^{Cn}$*mdh*·$^{Ec}$*yjhH* |
| pS621_CNMDH_YJHH_R | AGAAACGAUCCTCCGCATGATTAAGAT | |
| CNMDH_YJHH_ pS621_F | ATCGTTTCUATGAAAAAATTCAGCGGCATTATT | |
| CNMDH_YJHH_ pS621_R | AGCTCAGACUGGTAAAATGCCCTGCGCG | |
| pS621_CNMDH_DAPA_F | AGCTTTGCUGCTTAACCAGGCATCAAATAA | Construction of pS621·$^{Cn}$*mdh*·$^{Ec}$*dapA* |
| pS621_CNMDH_DAPA_R | AGAAACGAUCCTCCGCATGATTAAGAT | |
| CNMDH_DAPA_ pS621_F | ATCGTTTCUATGCCGCAGTCCGCGTTGT | |
| CNMDH_DAPA_ pS621_R | AGCAAAGCUTGAGCTGTTGCAGCAGGGTTT | |
| GG_GLLC_F | AACGTCTCGCTCGAATGAGCGGGCTGATCGGC | Construction of pS621·$^{Cn}$*mdh*·$^{Pp}$*gllC** |
| GG_GLLC_R | TTCGTCTCCCTCAAAGCGCCTTCCAGGTCGGTGAG | |
| GG_PP1791_F | AACGTCTCGCTCGAATGAAGCATCTGGACTGC | Construction of pS621·$^{Cn}$*mdh*·*PP_1791** |
| GG_PP1791_R | TTCGTCTCCCTCAAAGCTTTCAGTGGCCCATAAACGC | |
| pS621_CNMDH_GARL_F | ATAAGCTUAACCAGGCATCAAATAAAATAGA | Construction of pS621·$^{Cn}$*mdh*·$^{Ec}$*garL* |
| pS621_CNMDH_GARL_R | AGAAACGAUCCTCCGCATGATTAAGAT | |
| CNMDH_GARL_ pS621_F | ATCGTTTCUATGAATAACGATGTTTTC | |
| CNMDH_GARL_ pS621_R | AAGCTTAUTTTTTAAAGGTATCAGCC | |
| GG_YFAU_F | AACGTCTCGCTCGAATGAACGCATTATTAAGC | Construction of pS621·$^{Cn}$*mdh*·$^{Ec}$*yfaU** |
| GG_YFAU_R | TTCGTCTCCCTCAAAGCATAACTACCTTTTATGCGTG | |
| pS621_CNMDH_PAADL_F | ACGGCTGAGCUTAACCAGGCATCAAATAAAATAGA | Construction of pS621·$^{Cn}$*mdh*·$^{Pa}$*ADL* |
| pS621_CNMDH_PAADL_R | AGGTCCAUAGAAACGATCCTCCGCAT | |
| CNMDH_PAADL_ pS621_F | ATGGACCUGCCCGTCAATCGCTTCAAG | |
| CNMDH_PAADL_ pS621_R | AGCTCAGCCGUAGACCGAGGAACTGGCGG | |
| pS621-Cnmdh-GG(ltaEs)_F | ATGGCCGCUGCGAAGATAGCCATGACTTG | Construction of pS621·$^{Cn}$*mdh*·$^{GG}$*ltaEs** |
| pS621-Cnmdh-GG(ltaEs)_R | AAATAGAGUCGACCTGCAGGCATGCAA | |
| GG(ltaE)_pS621-Cnmdh_F | AGCGGCCAUGTAAAGAGGGGCCCAAGTTCAC | |
| GG(ltaE)_pS621-Cnmdh_R | ACTCTATTUTATTTGATGCCTGGTTAAGC | |
| pS621_Ptrc_MTK_MCL _F | ATCTAGAGUCGACCTGCAGGCATGCAA | Construction of pS621·$^{Mc}$*mtkBA*·$^{Rs}$*mcl* |
| pS621_Ptrc_MTK_MCL _R | AGTGGTGUGAAATTGTTATCCGCTCACAA | |
| MTK_MCL_pS621_Ptrc_F | ACACCACUGACCGAATTCACTAGTTAA | |
| MTK_MCL_pS621_Ptrc_R | ACTCTAGAUTAAGCACTAATCATCTCG | |
| pBAMD1-4_Scagx1_F | AGTAATCUAGAGTCGACCTGCAGGCA | Construction of pBAMD1-4·$^{Sc}$*agx1* |
| pBAMD1-4_Scagx1_R | AGAAACGAUCCTCCGCATGATTAAGATG | |
| Scagx1_ pBAMD1-4_F | ATCGTTTCUAATGACCAAAAGCGTCGATACG | |
| Scagx1_pBAMD1-4_R | AGATTACTUTTTGCGTTGCAGGGCTAG | |

#### Supplemental Notes

**Note S1.** MDF refers to the minimal metabolic driving force calculated by optimizing metabolite concentrations to render the pathway reactions as favorable as possible (Bachleitner et al., 2023). MDF is expressed as the inverse Gibbs free energy ($-\Delta_r G'$) and represents the feasibility as well as the flux carried through a given pathway. Metabolic pathways with low MDF values carry low fluxes and require increased metabolic engineering, while pathways with higher MDFs are favored (Bachleitner et al., 2023).

**Note S2.** This longer adaptation prevented us from testing the module in the Δ*serA* strain, as it was unclear whether the selection could be circumvented prior to the adaptation to produce glycine from glyoxylate. Using a Δ*ltaE* ΔΔ*glyA* auxotroph demonstrated long-term strain stability, required when implementing reactions with draining or reverse fluxes, as represented by this module.

**Note S3.** The HSC functionality was demonstrated using *E. coli* GarL, assimilating formaldehyde from sarcosine to complement the homoserine auxotrophy. This auxotrophy was obtained through the deletion of the homoserine dehydrogenase gene, which circumvents the addition of diaminopimelate or isoleucine (He et al., 2020). Homoserine can be metabolized to glycine without overexpressing *ltaE*, the *P. putida* gene encoding L-threonine aldolase (although upregulation increases the submodule efficiency). Since the metabolism of sarcosine yields one glycine besides formaldehyde, further characterization of the module was not carried out with this substrate.

**Note S4.** GarL was the only functional HOB aldolase in *P. putida*, although YfaU and YjhH have shown to be active in *E. coli* (He et al., 2020). We recently reported that *P. putida* is notably efficient at oxidizing $C_1$ molecules, harboring several dehydrogenases for each oxidation step (Turlin et al., 2023). HOB aldolase reaction relies on a side-reaction of the tested aldolases, and the substrate affinity (formaldehyde) has been determined to be rather low in variants such as Eda and YfaU (both from *E. coli*) or ADL from *P. aeruginosa* ($K_M$~25-500 mM). The concentration of formaldehyde produced from methanol could be too low to carry enough flux through HOB aldolase, due either to limited methanol oxidase activity or efficient formaldehyde detoxification (Bosch et al., 2021; Jeong et al., 2023; Wang et al., 2019).
